# Blueprinting expandable nanomaterials with standardized protein building blocks

**DOI:** 10.1101/2023.06.09.544258

**Authors:** Timothy F. Huddy, Yang Hsia, Ryan D. Kibler, Jinwei Xu, Neville Bethel, Deepesh Nagarajan, Rachel Redler, Philip J. Y. Leung, Alexis Courbet, Erin C. Yang, Asim K. Bera, Nicolas Coudray, S. John Calise, Fatima A. Davila-Hernandez, Connor Weidle, Hannah L. Han, Zhe Li, Ryan McHugh, Gabriella Reggiano, Alex Kang, Banumathi Sankaran, Miles S. Dickinson, Brian Coventry, TJ Brunette, Yulai Liu, Justas Dauparas, Andrew J. Borst, Damian Ekiert, Justin M. Kollman, Gira Bhabha, David Baker

## Abstract

A wooden house frame consists of many different lumber pieces, but because of the regularity of these building blocks, the structure can be designed using straightforward geometrical principles. The design of multicomponent protein assemblies in comparison has been much more complex, largely due to the irregular shapes of protein structures^1^. Here we describe extendable linear, curved, and angled protein building blocks, as well as inter-block interactions that conform to specified geometric standards; assemblies designed using these blocks inherit their extendability and regular interaction surfaces, enabling them to be expanded or contracted by varying the number of modules, and reinforced with secondary struts. Using X-ray crystallography and electron microscopy, we validate nanomaterial designs ranging from simple polygonal and circular oligomers that can be concentrically nested, up to large polyhedral nanocages and unbounded straight “train track” assemblies with reconfigurable sizes and geometries that can be readily blueprinted. Because of the complexity of protein structures and sequence-structure relationships, it has not been previously possible to build up large protein assemblies by deliberate placement of protein backbones onto a blank 3D canvas; the simplicity and geometric regularity of our design platform now enables construction of protein nanomaterials according to “back of an envelope” architectural blueprints.

## Introduction

There has been considerable recent progress in the design of protein nanomaterials including cyclic oligomers^2–4^, polyhedral nanocages^5–8^, one dimensional fibers^9, 10^, two dimensional sheets^11, 12^, and three dimensional crystals^10, 13^ by docking together^8^ or fusing^6, 14^ protein monomers or cyclic oligomers. While powerful, these methods have two limitations which arise from the irregularity of almost all protein structures. First, because the shapes of the constituent components are generally complex, they cannot be assembled into higher order structures based on simple geometric principles; instead, large scale sampling calculations are required for each case, and there is no guarantee that designable interfaces can be found. Second, like the myriad protein complexes in nature, the size of a designed protein assembly cannot be readily scaled; it is nearly impossible to make a smaller or larger but otherwise nearly identical version of assemblies made using current methods. In contrast, designed materials which extend along just one dimension, such as alpha helical coiled coils and repeat proteins, can be grown or shrunk by simply varying the length of the chain. There is a rich history of designing coiled coils using simple geometric principles; this extensibility and designability have made them widely used constituents of designed protein materials^15^.

We set out to develop a general approach for designing expandable higher order protein nanomaterials with the simplicity and programmability of coiled coil engineering. We reasoned that if a modular and regular tool-kit of building blocks and interactions could be generated consisting of (i) straight, ***Linear*** building blocks constructed from repeating sequence elements that extend without twisting as additional sequence repeats are added (Fig. 1B, 1C top, 1D), (ii) ***Curved*** building blocks that trace out arcs of circles of different radii (Fig. 1C bottom, 1D), and (iii) noncovalent arrangements that hold two building blocks in pre-specified relative orientations (Fig. 1D), then building up new nanostructures could in principle be done by inspection in a manner analogous to blueprinting a house frame (Fig. 1A). As with a house frame, the regular structures of the constituent building blocks could enable scaling the dimensions of the final architecture (area or volume) by simply altering the size of the constituent monomers, and structural reinforcement by placement of additional buttressing elements.

**Fig 1.**
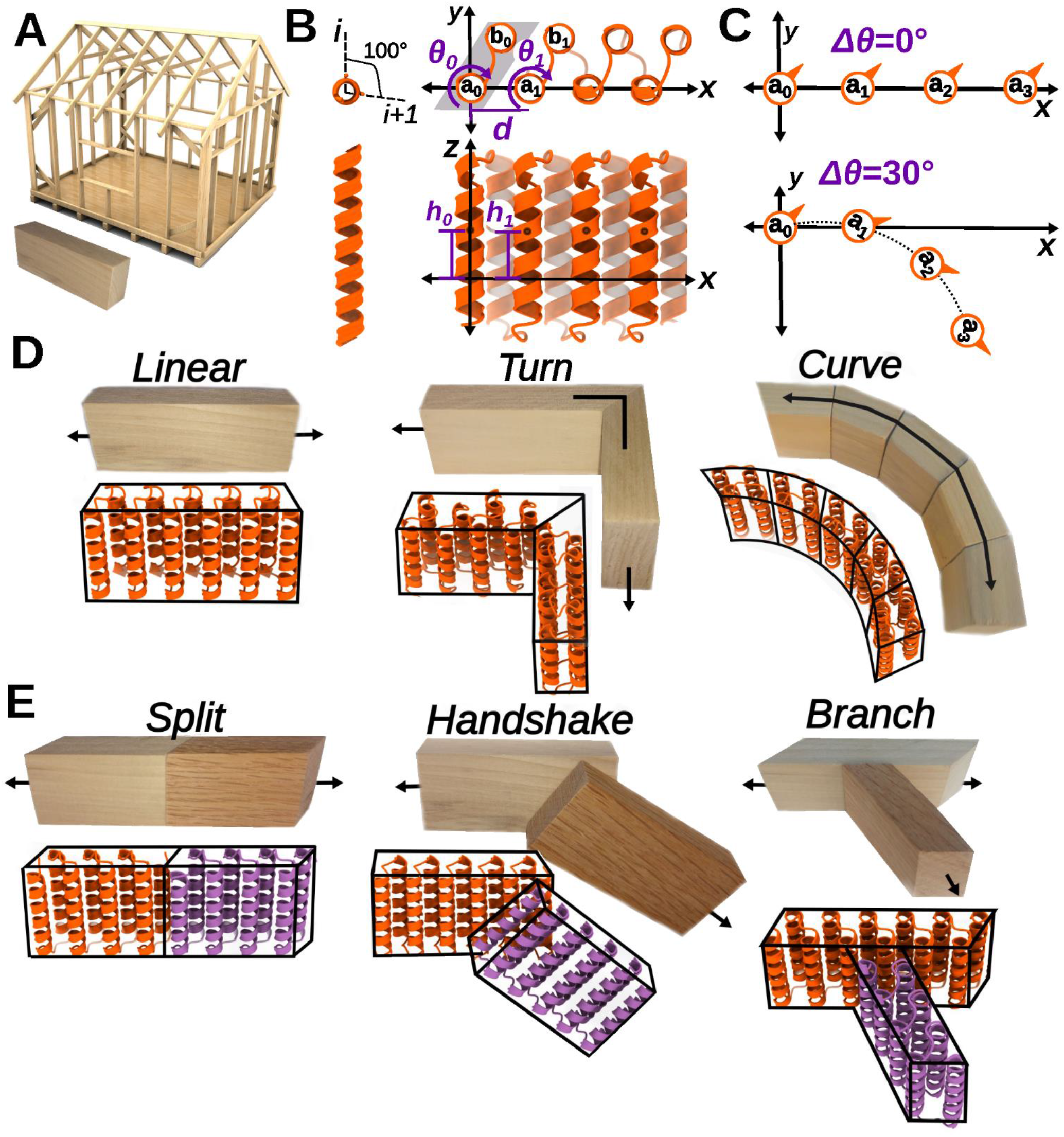
Overview of twistless helix repeat (THR) protein blocks and interaction modules. (A) Building a house frame using standardized wooden building blocks. (B) THR internal geometry. Blocks are constructed from idealized straight alpha helices with an angle of rotation between adjacent helices of Δθ; the remaining degrees of freedom that contribute to the repeat trajectory are also indicated. (C) Changing Δθ (while holding other parameters constant) specifically changes the curvature of the repeat trajectory. (D) Single chain THR modules. (E) THR interaction modules.

### Design of twist-less helix repeat (THR) protein blocks

Natural and previously designed proteins exhibit a wide range of helical geometries with local irregularities, kinks, and deviations from linearity ^16^ that make it difficult to achieve the properties illustrated in Fig 1 which enable simple nanomaterial scaling (beyond the one dimension accessed by varying the number of repeats in a repeat protein or coiled coil). To achieve these properties, we designed a series of new building blocks constructed from ideal alpha helices with all helical axes aligned. Restricting helical geometry to ideal straight helices with zero helical twist in principle considerably limits what types of structures could be built, but this is more than compensated by the great simplification of downstream material design, as illustrated below. We construct twist-less helix repeat (THRs) protein blocks from identical straight alpha helices (typically 2-4 helices in each unit); the length of the blocks can be varied simply by varying the number of repeat units. In contrast to existing natural and design repeat proteins^17^, THRs are constructed to enable modular nanomaterial design: ***Linear*** blocks are perfectly straight, allowing nanomaterials to be extended and contracted with no alteration in the angles between the constituent monomers, C***urve*** blocks have smoothly curving trajectories that stay in-plane, and ***Turn*** as well as interaction modules enable placement of two blocks in precise relative orientations with angles appropriate for regular material design.

We blueprint THRs by explicit placement of these straight helix structural elements using an extension of the principles used in coiled-coil and helical bundle design^16, 18^. A first helix ***a_0_***, part of the 0th repeat, is placed at the origin and aligned to the Z axis. A copy of ***a_0_*** called ***a_1_*** is then placed at a new location to set the rigid body transformation between the 0th and 1st (and all subsequent) repeat units. After this, any other helices (***b_0_, c_0_*** *, …*) that will be part of the repeating unit are placed as appropriate between ***a_0_*** and ***a_1_*** to provide more helices to pack against for stability, and the helices are connected with loops^19^; repetition of this basic unit then generates backbones with the desired geometries (Fig 1B, 1C)^17^. Since the helices are perfectly straight and parallel to the Z axis, the overall repeat protein trajectory is fully defined by the following transformation parameters from ***a_0_*** to ***a_1_***: the distance of displacement in the XY plane from helical axis to helical axis (***d***), the change in displacement in the Z axis direction (Δ***h***), and change in helix phase (Δ***θ***) (Fig 1B). The remaining degrees of freedom (DOFs) for the positions of helices ***b_0_, c_0_*** *, …*, which define the internal geometry of the repeat, are extensively sampled, sequences are designed using Rosetta FastDesign or ProteinMPNN^19, 20^, and designs are selected for experimental characterization based on packing and sequence-structure consistency metrics (see Methods). We obtained synthetic genes encoding the selected designs, expressed them in *Escherichia coli*, and purified the proteins using nickel-nitrilotriacetic acid (Ni-NTA) immobilized metal affinity chromatography (IMAC). Designs that were solubly expressed were analyzed by size exclusion chromatography (SEC) to determine oligomerization state, and in the case of assemblies a subset was analyzed by negative stain electron microscopy (nsEM). Experimental success rates and structural homogeneity for different classes of designs are summarized in Supplemental Figures 1 and 2, and Supplemental Discussion.

**Fig 2.**
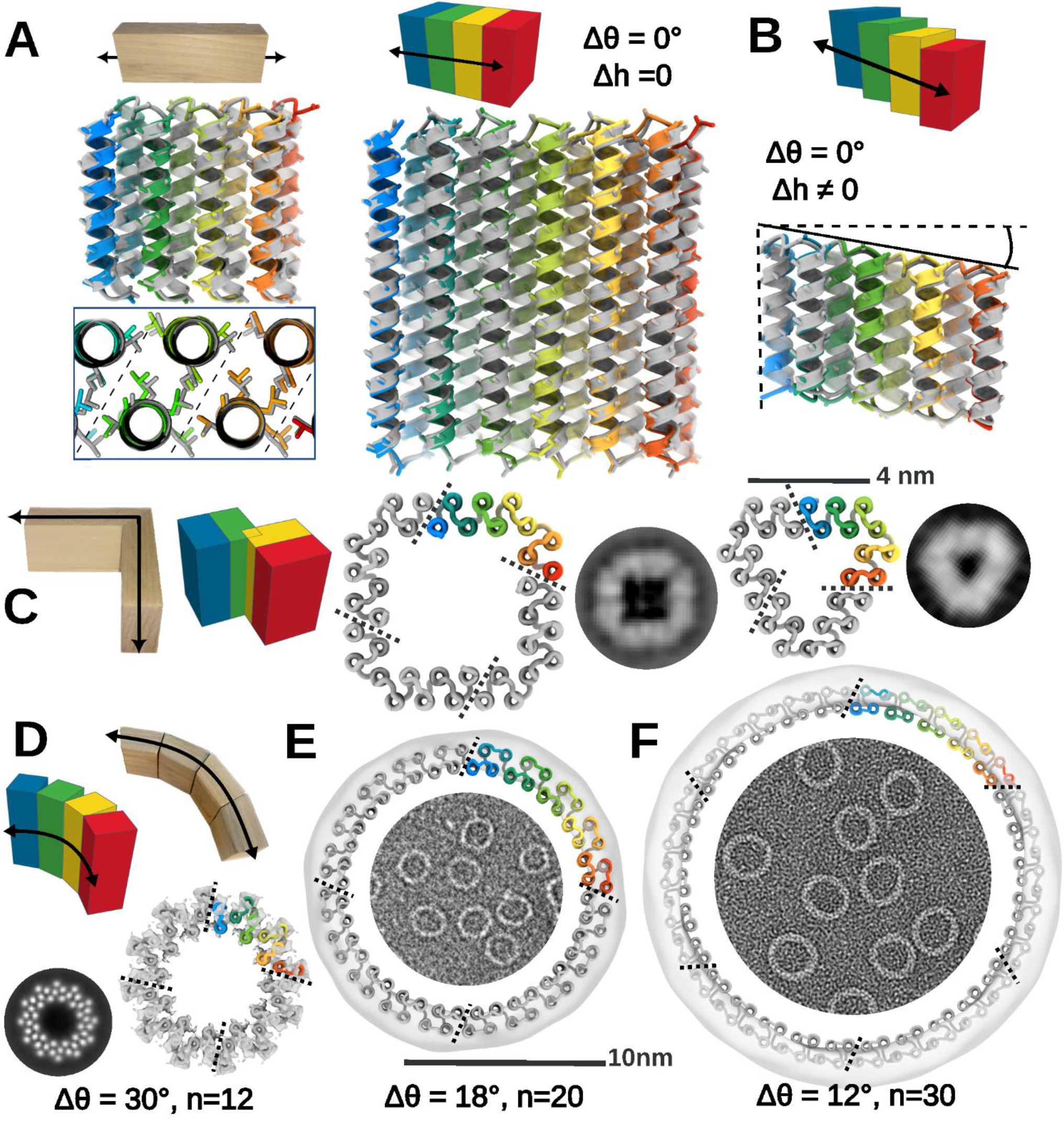
One-and two-dimensional shapes from THR blocks. (A,B) Linear THR design models (in rainbow) are nearly identical to the experimentally determined structures (grey). CA and CB atom sidechain sticks are shown to indicate helical phasing. (A) Left: 2.5 Å resolution crystal structure of short, linear THR1 has 0.8 Å CA RMSD to design model. Inset shows repeat packing in the THR interior. Middle/Right: 2.7 Å resolution crystal structure of tall, linear THR5 has 0.6 Å CA RMSD to design model. (B) Comparison of stair-stepping linear THR4 design model to cryo-EM structure (determined as part of a nanocage assembly; fig. S16). The CA RMSD between cryo structure and the design model is 1.0 Å. (C) C4 and C3 polygons generated from 4-helix Turn module THRs as illustrated on the left. Middle: C4 square 90_C4_B and right: C3 triangle 120_C3_A oligomers with representative nsEM 2D class averages for comparison (raw EM micrographs are in fig. S1F). Chain breaks are at the ends of the rainbow sections. The 4 nm scale bar is for the design models; class averages are not to scale. (D) Uncapped Curve THRs generate cyclic ring oligomers. The 12 repeat ring design R12B has a cryo-EM 3D reconstruction overlaid on the model; the two are nearly identical. A 2D class average with the individual straight helices resolved is shown left of the ring. (E) The 20 repeat ring design R20A has a nsEM reconstruction density overlaid on the model, and a raw micrograph is shown inside the ring. (F) The 30 repeat ring design R30A represented in similar manner to panel E. In D-F, the 10 nm scale bar is for the design models with reconstruction maps overlaid; class averages are not to scale. The asymmetric unit is colored in rainbow.

To generate straight, ***Linear*** THRs, we set ***Δθ*** to zero. As illustrated in Figure 2A and B, this results in perfectly straight repeat proteins in which each repeat unit is translated but not rotated relative to the previous unit. There are two subclasses: setting ***Δh*** = 0 generates repeat proteins with each repeat unit simply displaced in the X-Y plane (Fig. 2A), whereas setting ***Δh*** to a non-zero value generates repeat proteins that also step along the Z axis (Fig. 2B). We tested 33 ***Linear*** THRs (with ***Δh*** = 0) with helices either ∼20 or ∼40 residues in height; 23 of 33 tested designs were solubly expressed, and 13 of 19 designs analyzed by SEC were primarily monomeric as designed (fig. S1A-B, S2). Structural characterization of the ***Linear*** building blocks by X-ray crystallography individually and/or cryo electron microscopy (cryo-EM) in the context of assemblies (see below) revealed that both the detailed internal structures and the overall straight linear geometry were successfully achieved. The backbone RMSDs between the design models and crystal structures of three 20-residue helix designs (*THR1*, *THR2*, and *THR3*) and two 40-residue helix designs (*THR5* and *THR6*) were 0.8, 0.8, 0.4, 0.6, and 1.3 Å respectively, and in all five cases the relative rotation of successive repeats is nearly zero (Fig. 2A, S6A). We found that we could not only control ***Δθ*** =0, but also program values of the inter-repeat distance ***d***: the crystal structure of a design with ***d*** set to a compact helix packing value of 8.7 Å had a very close value of 8.6-8.8 Å at its central interior (*THR3*), in contrast to most others designed at 10.0 Å (fig. S6B). For a design with non-zero ***Δh***, the cryo-EM structure of an assembly constructed from such a block (*THR4*) exhibited a stair-stepping along Z but linear repeat structure nearly identical to the design model, backbone RMSD of 1.0 Å (Fig. 2B, S1A).

To generate ***Turn*** blocks, we blueprint an additional helix ***c_0_*** lined up with ***a_0_*** and ***a_1_*** that can be assigned any specified phase difference, which can be utilized in fusion operations to produce a ***Turn*** that is equal to ***θ_c_ - θ_a_*** (fig. S5D,E). As for all the THR blocks described here, because of the ideality of the block construction, the same sequence interactions can be used for the intra-block and inter-block interactions; we refer to blocks in which the terminal repeats have identical sequences as the internal repeats “uncapped”, and those in which the terminal helices have polar outward facing residues to prevent self-association (like the linear blocks above), as “capped”. We experimentally characterized uncapped ***Turn*** modules that generate rotations of 360/n where n is 3, 4, 5 or 6; if the geometry is correct these should oligomerize to form closed polygons with n subunits. nsEM 2D class averages of the n=3 designs clearly show the designed triangular shape with flattened corners (Fig. 2C, S1F), and for n=4, the designed square shapes (Fig. 2C, S1F) including fine details such as the lower density around the corner helix are observed. For n=5 and n=6, success rates were lower, likely because their hinge regions involved less extensive helix-helix interactions, but we did obtain designs with the expected polygonal structures for both after using reinforced corners on the C6 (fig. S1F, S1G, Supplemental Discussion). Thus, by controlling the phase rotations between adjacent helices, turns can be encoded while maintaining overall parallel helical architecture. We also made polygonal designs with combinations of ***Linear*** THRs and new straight helix-heterodimer corner junctions instead of ***Turn*** modules (Supplemental discussion, fig. S1G, S9, S10).

To generate ***Curve*** THRs, we incorporate a phase (***Δθ***) change between repeating elements (Fig. 1C) which generates a curved trajectory rather than a linear one. We choose ***Δθ*** to be a factor of 360 degrees so that perfectly closed rings can be generated. The parameters ***Δθ*** and the distance ***d*** between repeats can then be modulated to control the size of the ring (Supp Figure S7). To access a broad range of ***d*** parameter values, we add additional helices to the repeat unit; for circular rings we used 4 helices per repeat unit. A full ***Curve*** THR ring with ***n*** repeats can be divided into smaller chains each with ***m*** repeats, where ***m*** is a factor of ***n*** ; ***n / m*** uncapped repeats can associate to generate the full ring with cyclic symmetry^21^. To facilitate gene synthesis and protein production, we characterized such ***Split*** oligomeric versions of the rings rather than synthesizing very long single chains. We designed rings with 12, 18, 20, and 30 repeats ranging from 9 to 22 nm in outside diameter. The 12 and 20 repeat rings were tested as C4 designs, while the 18 and 30 repeat rings were tested as C6 designs. Designs for all four ring sizes were remarkably uniform with negative stain electron micrographs densely covered with circular assemblies with few to no defects or alternative structures present (fig S7). 2D class averages showed that designs for all four sizes were close to the intended size (Fig 2D; 10, 1, and 9 unique designs yielded distinct ring shapes for 18, 20 and 30 repeat rings; fig. S1E, S2)). The smallest rings with 12 repeats have solvent-exposed helices exterior to the ring placed to facilitate outward-facing fusions without disrupting the core packing of the ring; these are clearly visible in the 2D class averages and 5.3 Å resolution cryo-EM reconstruction of *R12B* (Fig. 2D, S1E) that shows correct patterning of the helices. Negative stain EM of the 18, 20, and 30 repeat rings (with outside diameters of 12, 14 and 22 nm respectively) showed that many designs formed remarkably monodisperse populations of ring-like structures closely consistent with the design models (Fig. 2E, F, S1E). Negative stain EM class averages of these designs had the smooth and round shape of the design models and were in most but not all cases homogeneous (some designs assembled into closed ring species that ranged by +/-1 chain of the desired number, resulting in some slightly oblong shapes (fig. S1E)). These designs highlight the control over ring curvature that can be achieved by specifying repeat parameters.

The simplicity of our blocks in principle enables the reinforcing of designed materials using struts rigidly linking distinct structural elements. As a first test of this, we sought to build concentric ring assemblies from pairs of rings that have different sizes but repeat numbers that share large common denominators. For example, 2 repeat units of a 20-repeat ring can be combined with 3 repeat units of a 30 repeat ring since ten copies generate a complete ring in both cases (Fig. 3A, left). Rings were segmented into matching cyclic symmetries, the rotation and Z displacement of one ring relative to the other was sampled, and linear THRs were placed to connect the inner and outer rings. We constructed single component C10 concentric ring assemblies by connecting a three repeat unit curved block and a two repeat unit curved block that both generate a 36° (360°/10) rotation with a radially oriented strut. 2D class averages of nsEM images of the designed strutted assemblies show both rings clearly present (Fig. 3A, right; some 11 subunit rings were observed in addition to the target 10 subunit structure). We similarly connected three repeat units with a 20° rotation per repeat, and five repeat units with a 12° rotation per repeat, with a radial strut; the resulting composite subunits map out a 60° rotation of inner and outer rings such that six subunits generate a full 360° ring. The resulting two component C6 strutted assembly yielded 2D class averages that showed both rings with all chains present, and a 5.1 Å cryo-EM reconstruction was very close to the design model (RMSD 2.7 Å) with very similar outer diameter (19.7 nm vs 20.1 nm, Fig. 3B, S8C). The helix positioning in the inner ring and the strut are also very close to the design model (fig. S8C inset boxes). Thus, the modularity of the THRs enables designing complex structures by inspection and enables buttressing to increase structural robustness (see Supplemental Discussion and Supplemental Figure S8).

**Fig 3.**
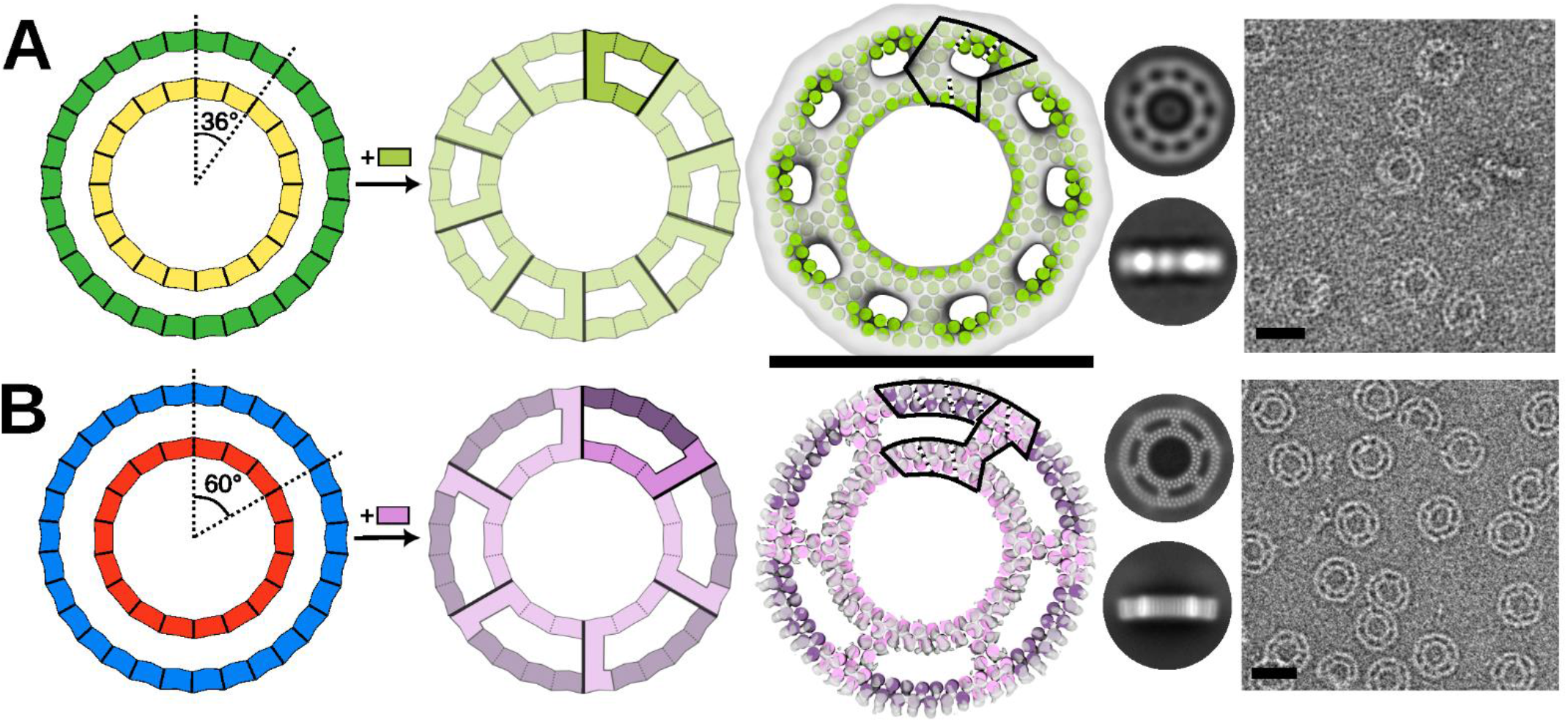
Design of strutted double rings. (A-B) 2 different size rings using ***Curve*** THRs for which integral multiples generate the same rotation can be concentrically nested and connected by struts. (A) Three repeats of an outer ring (12° per repeat) are combined with 2 repeats of an inner ring (18° per repeat) which both generate a 36° rotation. Connection of the 2 pieces with a ***Linear*** THR generates a C10 single component ring (called *strut_C10_8*); an asymmetric unit is highlighted in the second ring image. A nsEM 3D reconstruction in C10 symmetry is shown overlaid with the design model next to 2D class averages and a representative micrograph. (B) Five repeats of an outer ring (12° per repeat) are combined with 3 repeats of an inner ring (20° per repeat) which both generate a 60° rotation. Connection of the 2 pieces with a ***Linear*** THR and an additional chain break in the outer ring generates a C6 two component ring (called *strut_C6_21*); an asymmetric unit is highlighted in the second ring image. A cryo-EM 3D reconstruction in C6 symmetry is shown overlaid with the design model next to cryo-EM 2D class averages and a representative nsEM micrograph (Additional cryo-EM details in Supplemental Figure S8C). (A-B) Scale bars are 20 nm. The asymmetric unit is outlined on top of the design model and repeats are sectioned with dashed lines.

### Expandable nanomaterials

The regularity of our blocks in principle enables scaling the size of nanomaterial designs simply by changing the number of repeats in the constituent THRs without altering any of the inter-block interfaces. How the THRs must be aligned to enable expandability differs for each architecture, as described below.

To construct expandable cyclic assemblies, the ***Linear*** THRs must be placed such that the propagation axis is normal to the cyclic symmetry axis. For cyclic designs with this property (those built from ***Turn*** modules or heterodimers (see Supplemental Discussion)), adding or removing repeats simply changes the length of the oligomer edge without affecting the interface between monomers. We tested this expandability with a C4 “square” (*sC4*) oligomer for which we had obtained a cryo-EM reconstruction with 1.6 Å backbone RMSD (fig. S10). This subunit consists of a central ***Linear*** THR flanked by straight-helix heterodimers that produce a 90° turn. To expand this structure, we inserted two additional repeat units (6 helices) into the ***Linear*** THR portion of the subunit. Cryo-EM 2D class averages for both the original and expanded square show close agreement to the design models and clear expansion; the helices clearly remain aligned to the Z axis as designed (Fig. 5A).

Architectures with polyhedral nanocage symmetry can be similarly expanded provided that the ***Linear*** THR propagation axis can be arranged parallel to the plane formed by the two symmetry axes that are being spanned by the THR (fig. S11A). To generate such architectures, and enable further access to construction in three dimensions, we designed out-of-plane interactions between building blocks. We first focused on designing C2 symmetric interfaces in which the angles between ***Linear*** THRs correspond to the angles needed to generate regular polyhedral symmetry (Fig. 4) when combined with planar C3 or C4 components, while also satisfying expandability criteria. For an octahedral “cube” (O4) built from flat objects with C4 symmetry that lie on the “cube faces”, this angle is 90 degrees. For tetrahedra (T3), octahedra (O3), and icosahedra(I3) built from flat C3 symmetric objects, the out-of-plane ***Handshake*** angles that are needed to join the flat objects are 70.5°, 109.5°, and 138.2°, respectively ^22, 23^. ***Handshake*** C2 homodimers were generated by fixing this out-of-plane angle and keeping the ***Linear*** THR propagation axes parallel to each other, only sampling the offset spacing between the THRs ^8^ (fig. S12).

To generate expandable nanocages, flat cyclic components that form the “faces” of the cages were linked via the appropriate angle ***Handshake***. For the flat cyclic component, we used a ring design with 12 repeats (*R12B*, Fig 2C) constructed from ***Curve*** units, and split the 12 repeats into either three subunits with four repeats each (C3) or four subunits with three repeats each (C4) (fig. S3D), depending on the desired polyhedral symmetry architecture. To facilitate connection via the C2 ***Handshakes***, we fused ***Linear*** THR arms onto each subunit constrained to point outward parallel to a radial vector emanating from the symmetry axis, but offset such that when the C2 interface is formed, the C2 axis is along a radial vector (Fig. 4A, S12). Tetrahedral, octahedral “cubic” and icosahedral structures with C3 rings at respective axes (T3, O3, and I3), and octahedral structures with C4 rings at the respective axes (O4) were constructed by incorporation of the appropriate C2 interface. For example, to make a “cubic” octahedral nanocage, we incorporate into the C4 ring arm the 90° C2 ***Handshake*** module (Fig. 4E) by simple sequence concatenation. Synthetic genes were obtained for 13 nanocage designs; all 13 expressed solubly, 10 had SEC elution profiles that suggested cage formation, 8 yielded particles with the expected size by nsEM, and 7 gave 2D class averages and symmetric 3D reconstructions that resembled the design models. Successful designs for each architecture are shown in Fig. 4 B,C,D,E, S1J, S12. The designed geometric features including the spindle-like 2-fold ***Handshake*** interface and the flat “in-plane” ring areas with distinct holes are clearly evident. For the T3 and O4 cages, the correct species dominated, but in O3 and I3 cages there were noticeable populations of species that were either partially formed or broken under nsEM conditions (fig. S13). A 7.5 Å cryo-EM reconstruction and experimental model were obtained for the cubic cage built from tetrameric rings on the faces (*cage_O4_34*) which was very close to the design model with the straight helices clearly evident and only very slight deviations in the arm alignment (Fig. 4E, S14). These results illustrate the robustness of structures that can be assembled from our regularized building blocks using simple “snapping together” of complementary pieces, and suggest that there are surprisingly few limitations on the designable interface angles between THRs, as the selected angles were chosen without any consideration of “knob-and-hole” helical packing or other features ^24^.

**Fig 4.**
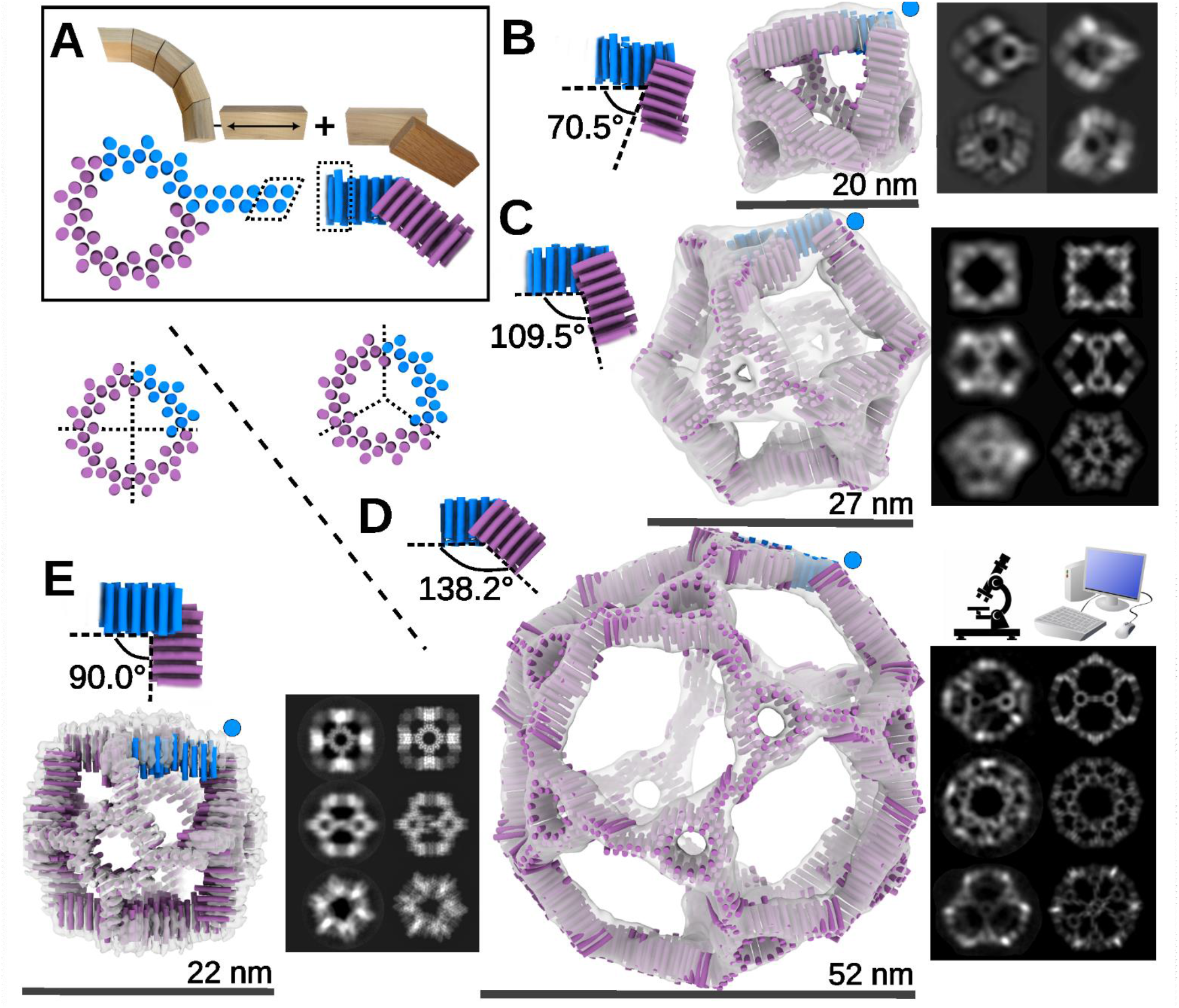
Modular construction of protein nanocages from THRs. (A) Regular nanocages are constructed from ***Curve*** THR rings with ***Linear*** arms projecting outward that can be linearly extended (left) and designed ***Handshake*** C2 interfaces (right) that hold the rings at the angle required for the desired polyhedral symmetry. THR chains that are fused into one chain are shown in blue, with identical backbone and sequence areas used for concatenation indicated in the dashed boxes. (B-E) Nanocage designs from the components shown in A. C2 ***Handshake*** modules generating the required angles (top left in each panel) are combined with either C3 (B, C, D) or C4 (E) versions of the ring (panels below boxed region in A). Design models are shown as helix cylinders, with the asymmetric unit in blue, and the remaining copies in purple. Each cage is overlaid with the 3D nsEM reconstruction, with representative paired 2D classes on the right (left is experimental, right is simulated from design model). Blue dots indicate the location of the handshake angle in the cage. (B) T3 tetrahedral design *cage_T3_101* uses a C3 ring and 70.5° C2 handshake. (C) O3 octahedral design *cage_O3_20* uses a C3 ring and 109.5° C2 handshake. (D) I3 icosahedral design *cage_I3_8* uses a C3 ring and a 138.2° C2 handshake. (E) O4 octahedral design *cage_O4_34* uses a C4 ring and a 90.0° C2 handshake.

We tested the expansion in all three dimensions of the cubic design (Fig 4E, S14) by increasing the number of repeat units in the linear arm. We generated four different sizes of the *cage_O4_34* by increasing the number of THR helices in the arm by +0, +4, +8, or +12 helices (Fig. 5B, S13). For all sizes, nsEM 2D class averages (Fig. 5B, bottom row) show all three symmetrical views with clearly increasing size but otherwise close preservation of architecture. 3D nsEM reconstructions were consistent with corresponding design models, with the overall cube shape and ring circular pore clearly visible in each of the sizes (Fig. 5B, top row). The first three sizes of cage show primarily intact assemblies across the nsEM grids; for the largest size (+12) some incomplete assembly was also observed (fig. S13). Additional single-component expandable nanocage designs are described in Supplemental Figures S13, S16, S17, and Supplemental Discussion.

We next designed two-component expandable nanocages by locking the rotation DOF of a THR-containing building block to maintain the expandability constraint (see Methods), and then docking it against a freely sampling partner oligomer to form an O43 architecture (Fig. 5C, Supplemental Discussion, fig. S19-23). Expandability over four different sizes was achieved with *cage_O43_129* (+0, +4, +8, +12 helices). The internal structure of the oligomers is clearly resolved in nsEM reconstructions; the distance between the center of mass (COM) of the tetramer component to the COM of the trimer component across the different sizes is 8.1, 9.1, 10.0, and 11.1 nm respectively (Fig. 5C, S21). Views down each of the three symmetry axes (2-fold, 3-fold, and 4-fold) are clear for each size (except for 3-fold view in the largest size) with rotational deviations of the 4-fold cyclic component compared to the design model, while the rotation of the 3-fold cyclic component holding the THR remains unperturbed as designed (Fig 5C, Supp Figure S22. A fifth size (16 additional helices) assembled into cage-like structures but the populations were too heterogeneous for detailed characterization (fig. S21).

**Fig 5.**
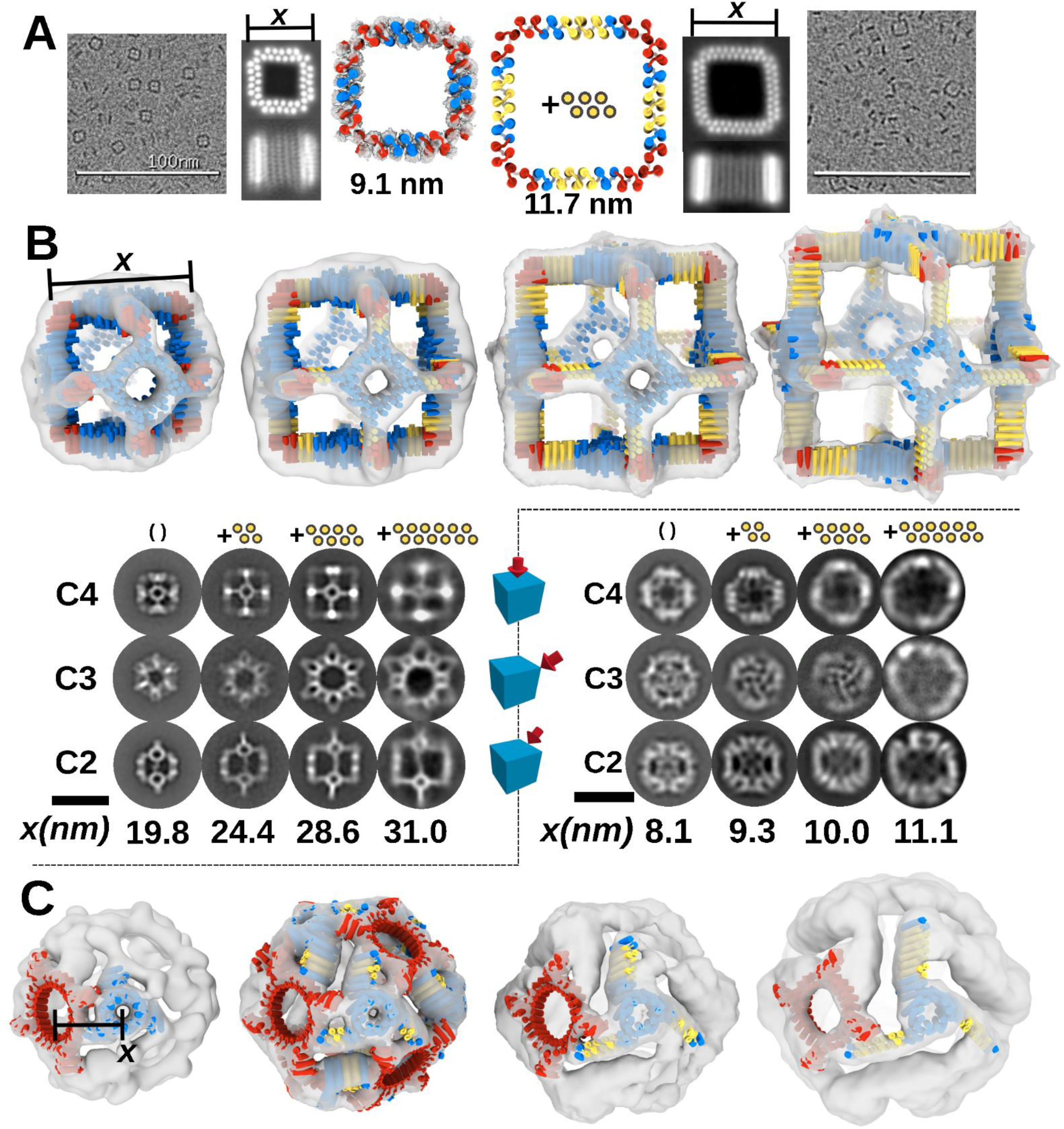
Expandable THR-based nanomaterials. (A) Expandable C4 square. Left: base design (*sC4)*; Right, expanded version with 6 additional helices per chain (*sC4+6)*. Far left and right: representative cryo EM micrographs (100 nm scale bars) with adjacent 2D class averages. A 3D reconstruction is shown superimposed on the base size design model. Chain breaks are in the red heterodimer region. A dimension labeled “*x”* is measured across the width of the square on the 2D class averages. (B) Expandable O4 octahedral handshake nanocage, *cage_04_34*. Top left to right, design models and negative stain reconstructions of designs extended by 0, 4, 8 and 12 helices. Bottom/left: Class averages along the three symmetry axes (rows) for increasing size designs (columns). The number of inserted helices is shown above each column. Scale bar (30 nm) for 2D class averages is located under “C2’’ text. A dimension labeled “*x”* is measured between outside corners of handshake density from the nsEM map volumes set to be as tall as the design model helices. (C) Expandible two-component O43 nanocage *cage_043_129* . Bottom row: nsEM reconstruction density overlaid on the corresponding design model for four different sizes. The extendable trimer component is in blue and the constant tetramer component in red. Top row (right side): 2D class averages along the 3 symmetry axes for each of the four sizes. In the +8 and +12 structures the tetramer component is slightly rotated, and was re-aligned into the ns EM map for clarity. The indicated distance “x” is between the experimentally determined center of mass of neighboring C3 and C4 components. Scale bar for 2D class averages is 25nm. Protein design models are in cylindrical helix representation, yellow represents regions that were extended by locally repeating the linear THR structure/sequence.

For unbounded architectures which extend along one or more axes, extensibility requires that the linear THR propagation axes be parallel to the extension axes. We constructed an anti-parallel assembly with an overall “train track” shape from THR modules (Fig. 6A). The “rails” of the track are ***Linear*** THRs that are uncapped to allow for unbounded linear assembly end-to-end, and **C2** “ties” dock onto ***Branch*** interfaces on the sides of the rails, organizing them into strutted antiparallel pairs. Adding repeats to the rails increases the spacing between ties (along the helical axis) and adding repeats to the ties increases the separation distance between rails along a different axis (Fig. 6B). We used 12 helix addition to the rail to double the spacing between ties, and 8 helix addition to the tie to roughly double the length of the tie. For the 4 combinations of component sizes, we obtained nsEM 2D class averages consistent with the design models (compare Fig. 6B and D). Train track assembly was robust to fusion of *mScarlet-i* on rails both at termini and in an internal loop (fig S24B), and *sfGFP* on the ties ^25, 26^, as monitored by nsEM, with density observed for the *GFP*, fig. S24C).

**Fig 6.**
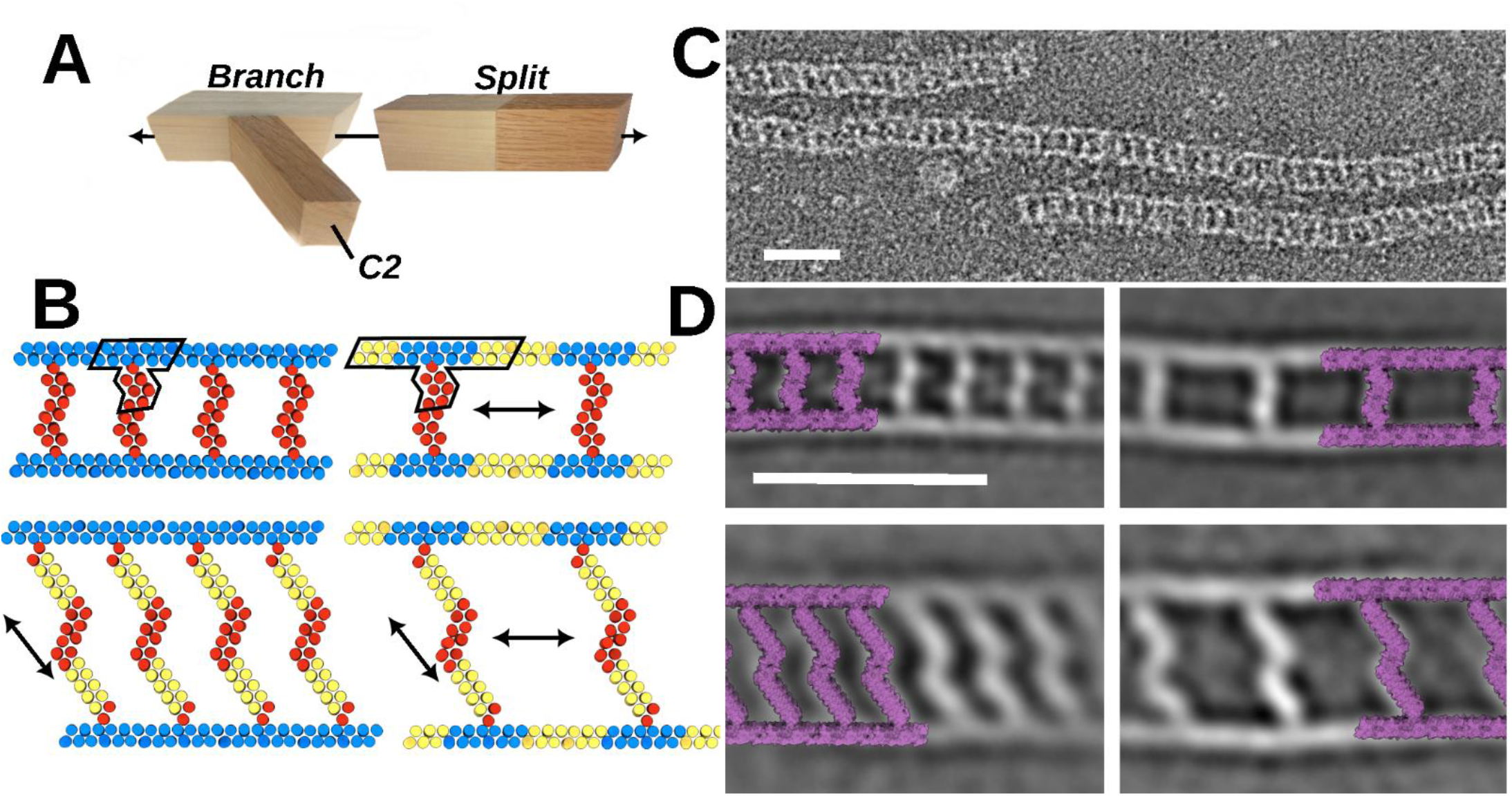
Designed “train track” fibers. **(A)** Components of the train track designs: a ***Branch*** module, a ***Split*** module, and a C2 interface module. (B) Train track designs. The asymmetric unit is outlined in the top left design; red and blue are unique protein chains. Taking advantage of the extensibility of the building blocks, four different train track designs were created, starting with the design at the top left. Top right: increase of spacing between rungs by expanding the ***Split*** module. Bottom left, increasing the length of the rungs by expanding the C2 module. Bottom right: increasing both rung spacing and length by expanding both the ***Split*** and C2 modules. Inserted segments are in yellow. (C) nsEM micrograph of design at top left of B. Scale bar = 25 nm (D) nsEM 2D class averages of the four designs in B, shown in the same overall layout. The design models are overlaid on the class averages at 1:1 scale. Scale bar = 25 nm.

## Discussion

Upon determining the first low-resolution model of a globular protein’s structure (myoglobin), John Kendrew wrote in 1958 that “Perhaps the most remarkable features of the molecule are its complexity and its lack of symmetry. The arrangement seems to be almost totally lacking in the kind of regularities which one instinctively anticipates”^27^. Over six decades of structural biology research have shown this to be a generally appropriate description of protein structure^1^. Figs 2-5 show that this complexity is not an inherent feature of the polypeptide chain: the simplicity and regularity of our designed materials approaches that of the wooden beams used for constructing a house frame. This enables the resizing of designed materials in 2 and 3 dimensions simply by changing the numbers of repeat units in the THR modules with little or no need for detailed design calculations; previously this has only been possible with coiled coils and repeat proteins with open helical symmetries (propagating along a single axis) ^9, 15, 28^. The flat surfaces and regular geometry have immediate applications to the design of bio-mineralizing systems: THR monomers presenting carboxylate groups in regular arrays nucleate the mineralization of carbonate into calcite^29^ and expandable THR systems such as the cubic assemblies in Fig 3 presenting such arrays could provide a route to hierarchical protein-mineral hybrid materials.

There are exciting paths forward to further increase the capabilities of our programmable THR platform. First, our current multi-subunit assemblies have high symmetry, and assembly of arbitrary nanostructures would require breaking symmetry – one approach to achieving this would be to build heterodimeric and heterotrimeric interfaces between THRs, which would enable considerable shape diversification and addressability of each protein chain^30^. This would allow access to a broader range of asymmetric nanostructures, as with DNA nanotechnology bricks, tiles, and slats^31–34^, but with the higher precision and greater functionality of proteins. Second, the materials generated here all form through self-assembly, but as the number of components increases the overall yield of the desired product could decrease. This limitation could potentially be overcome by stepwise solid phase assembly with cross linking after addition of each THR component (as in solid phase peptide or DNA synthesis, but in 3D with location of addition specified by non-covalent interactions between the THRs; the analogue in construction is nailing lumber pieces together after alignment). The combination of symmetry breaking and stepwise assembly would enable the design of a very wide range of protein nanomaterials based on simple geometric sketches which could be readily genetically modified to present a wide variety of functional domains in precisely controllable relative orientations.

## Supporting information

sC4_structure

## Acknowledgments

We thank F. Busch and V. Wysocki at Ohio State University for providing native mass spectrometry experiments that helped debug some of our early designs. Thanks to J. Decarreau for support in trying out optical microscopy with fibers. Thanks to J. Quispe and S. Dickinson at University of Washington and Veer S. Bhatt at Thermo Fisher Scientific for their assistance with cryo-EM data collection. Thanks to L. Milles and B. Wicky for wet lab assistance. Thanks to D. Hicks, H. Pyles, and W. Sheffler for computational assistance. Thanks to S. Boyken, C. Hague, J. Bai, and L. Stewart for helpful perspective and discussion. Thanks to F. Praetorius for manuscript editing.

We thank the Arnold and Mabel Beckman Cryo-EM Center at the University of Washington for electron microscope use.

We want to thank the Advanced Light Source (ALS) beamlines 8.2.1/ 5.0.3/ 8.2.2 at Lawrence Berkeley National Laboratory for X-ray crystallography data collection. The Berkeley Center for Structural Biology is supported in part by the National Institutes of Health (NIH), National Institute of General Medical Sciences, and the Howard Hughes Medical Institute. The ALS is supported by the Director, Office of Science, Office of Basic Energy Sciences and US Department of Energy (DOE) (DE-AC02-05CH11231).

## Funding

The IPD (Institute for Protein Design) Breakthrough Fund, for “De novo design of 100nm scale protein assemblies” (T.H., Y.H., R.D.K., D.B.), and “De novo design of selective pores” (Y.L., D.B.), The Audacious Project at the Institute for Protein Design (T.H., J.X., E.C.Y., A.J.B., H.H., Z.L., R.M., A.K., D.B.), The Open Philanthropy Project Improving Protein Design Fund (Y.H., R.R., P.J.Y.L., A.K.B., D.E., G.B., D.B.), National Science Foundation (NSF) award CHE-1629214 (D.N., D.B.), the Helen Hay Whitney Foundation (S.J.C), a gift from Microsoft (D.J., J.D., D.B.), The Donald and Jo Anne Petersen Endowment for Accelerating Advancements in Alzheimer’s Disease Research (T.J.B., D.B.), and the Howard Hughes Medical Institute (N.B., A.C., B.C., D.B.).

SAXS data were collected at the Advanced Light Source (ALS) SIBYLS beamline on behalf of US DOE-BER, through the Integrated Diffraction Analysis Technologies (IDAT) program.

Part of the cryo-EM data were collected on a Glacios TEM (Thermo Fisher Scientific, MA, USA) from NIH award S10OD023476 (J.K.).

Parts of the cryo-EM work were supported by Open Philanthropy subcontract via UW.

Parts of the cryo-EM data processing were supported by the High Performance Computing (HPC) facility at NYU School of Medicine.

Some of the cryo-EM grids were screened at Cryo-Electron Microscopy Laboratory Core at NYU School of Medicine (RRID: SCR_019202) and we thank the cryo-EM core staff for their assistance.

Some of the cryo-EM data acquisition was performed at the Simons Electron Microscopy Center and National Resource for Automated Molecular Microscopy (SEMC) and National Center for cryo-EM Access and Technology (NCCAT) located at the New York Structural Biology Center, supported by grants from the Simons Foundation (SF349247) and the NIH National Institute of General Medical Sciences (GM103310, U24 GM129539).

## Author contributions

Conceptualization: TFH, YH, TB, DB, RDK, JX, DN, PJYL

Methodology: TFH, YH, JX, RDK, PJYL, ECY, BC, TB, JD

Investigation: TFH, YH, JX, RDK, NB, DN, RR, PJYL, SJC, AC, AJB, FADH, AKB, HLH, CW, ZL, RM, GR, AK, BS, MSD, YL

Visualization: TFH, YH, RDK, NC

Funding acquisition: DB, GB, DE, TFH, RDK, YH, AJB

Supervision: DB, GB, JMK, DE

Writing – original draft: TFH, YH

Writing – review & editing: TFH, DB, YH, RDK, ECY, AB, GB, PJYL, SJC, NC

## Data and materials availability

All data are available in the main text or the supplementary materials. EM maps have been deposited in the Electron Microscopy Data Bank (*strut_C6_21:* EMD-29893, *cage_04_34*: EMD-29915, *sC4:* EMD-29974, *cage_03_10*: EMD-40070 (C1 asymmetric) and EMD-40071 (octahedral symmetric), *cage_T3_5*: EMD-40075 (C1 asymmetric) and EMD-40074 (tetrahedral symmetric) and EMD-40073 (C1, 1 chain missing) and EMD-40073 (1 trimer missing), *cage_T3_5_+2*: EMD-40076). Crystallographic datasets have been deposited in the PDB (*THR1:***8G9J***, THR2:* **8G9K***, THR5:***8GA7***, THR6:* **8GA6**). Experimental model from cryo-EM of “square” *sC4* is deposited in the PDB as **8GEL**. An example script and input for generating THR building blocks are provided at (https://github.com/tfhuddy/2023-manuscript-materials).

## Supplementary materials

### Materials and methods

#### Computational design

##### Placement of straight alpha helices for THRs (both linear and curved) and SHDs

The Rosetta protein design software suite has an implementation of Crick-parameter ^16^ generated secondary structures. The MakeBundle or BundleGridSampler movers in RosettaScripts ^18^ can both be used to generate preset idealized helices. We use both the “alpha_helix” and “alpha_helix_100” parameter presets; if blocks will be combined by fusion later, then it could be beneficial to use the same parameter sets for all blocks. The “alpha_helix_100” preset also features the repeating helical phase every 18 residues, so structures can be modularly lengthened or shortened at the level of helix length (fig. S4).

Both movers offer functionality for placing helices in the X-Y plane with specific phase and position. For a simple THR design, it is easiest to place an initial helix at the origin, and then place other helices relative to that one. We find that distances from the center of 1 helix to the center of an adjacent helix can be in the range of 8.5 Å to 11.5 Å to offer favorable packing solutions between them. The most favorable spacing to find solutions seems to be near 10.5 Å. Sampling the phases of adjacent helices is primarily how different packing solutions are found (although sometimes DOFs are locked because distance/phase relationships are used to establish repeat trajectory). It is also reasonable to use Python/PyRosetta ^35^ to position copies of a template ideal helix.

For making a repeat protein, all helices of the first repeat unit plus one copy of the first helix are generated with explicitly defined positioning. Then, the ConnectChainsMover is used to install loops between helices using helix-loop-helix structural fragments (available for download at: files.ipd.uw.edu/pub/modular_repeat_protein_2020/ss_grouped_vall_all.h5) that were harvested from the PDB^17^. Once this looped “repeat unit +1” structure is made, then the RepeatPropagationMover is used to generate perfect repeat structures that contain however many repeats are desired.

An example script for this building block generation is provided at https://github.com/tfhuddy/2023-manuscript-materials. This example uses a dummy input to an annotated RosettaScripts xml script, which will be used with the Rosetta executable to produce a ***Linear*** THR backbone.

##### Sequence design on protein backbones

We find ProteinMPNN to be sufficient for all our sequence design operations ^20^. For repeat proteins and homomeric oligomers, it is possible to restrict sequences to be identical between the structural elements where that is desired, using the “--tied_positions” arguments described at https://github.com/dauparas/ProteinMPNN.

##### Structure prediction on designed sequences

We primarily use AlphaFold2 ^36^ to judge if our designed sequences will likely fold or assemble as desired. Models 4 and 5 tend to give the best confidence for these all alpha-helical designed proteins, and often these are the only models used. Designs are judged to be good if they have pLDDT > 90, good pTM score (typically > 0.80) and low CA RMSD (typically < 1.5 Å) to the ideal design model.

##### Generation of protein assemblies

Curved THR rings and train-track rails are made by producing a monomer length as needed with RepeatPropagationMover, and leaving the outer ends of the proteins with intact repeat sequence so they can use the repeat interactions between chains.

Angle-encoded THRs being used to make cyclic oligomers can theoretically be done easily by hand in PyMOL, but we used the WORMS protocol to find the intended idealized closure solutions. Inputs to worms were split THRs (an THR with a loop region deleted such that it became a 2-chain input) fused to copies of themselves as described in the “Crown” structures in the reference publication ^6^.

THRs can also be combined with SHDs or other proteins via the WORMs protocol to make similar structures, such as in design “sC4”.

RPXDock can also be used to generate cyclic oligomers without requiring any fusion operations; just by docking THRs against each other, such as in design “hex_C6” (fig. S1G) ^8^.

For the C2 helical bundle design at the center of the “TT_C2” design, helical placement can be done with the intention that symmetric copies can be generated with appropriate rotation about the Z axis with the SetupForSymmetry mover, as detailed in previous work with these movers ^37^.

Handshake C2 designs were made in RPXDock by first positioning the THRs such that their repeat axes were parallel to the X axis but offset from it as shown in figure S12. These were then treated as if they were cyclic oligomers already (or an arm extension on an “invisible” cyclic oligomer) and docked into the desired nanocage symmetry that would correspond to the desired THR handshake angle. Designs were selected that maintained the 2 copies of THR at the handshake with their repeat axes parallel to each other and were then designed with proteinMPNN and evaluated with Alphafold2 as dimers.

Arm fusions to designs were done with either: (A) the HelixFuse wrapper for the MergePDBMover, as was done with fuse_2, fuse_3, and fuse_19 (*5*) or (B) RPXDock was used to dock arm to the outside of cyclic oligomer (treating arm as a fake cyclic oligomer as mentioned above, and then using RPXDock Axle sampler ^8^; multiple arm position inputs were used for this) and then solutions with chain termini at appropriate positions were looped together with ConnectChainsMover, as was done for the ring fusions in Figure 4 and with the struts added to rings in Figure 3. For the concentric ring designs in Figure 3, the RPXDock Axle sampling was repeated after the first round yielded inner ring + strut so that the outer ring height and rotation could be sampled to dock against the strutted inner pieces.

For the 2 component nanocages shown, RPXDock was used with constraints. Cyclic oligomers with THR arms on them were first pre-oriented such that the arm propagation was aligned to the x-axis. For each symmetry, the x-axis aligned oligomer was additionally rotated to match the symmetry axis of the partner symmetry; for example, in the O43 case, the THR-containing oligomer was rotated +45° around the z-axis. This component was then restricted from rotating during the RPXDock sampling using the --fixed_rot option, while the other component was free to sample all normally available DOFs.

For designing interactions between THRs in the case of the train track fibril designs, it did not matter how the tie was docked against the rail so long as the helices remained parallel. For this, samplings of the rigid body positions between the 2 pieces can be obtained by using helix fusion protocols as previously described^6^, where the stationary structure is the rail, and the mobile structure is the tie. Outputs of helix fusion then had the fusion helix truncated off of the rail, and the tie was restored to its original length such that the resulting poses feature a tie butting up against the rail, making contacts with one helix of the tie as if it were a continuation of the THR in the rail.

#### Experimental methods

##### Construction of synthetic genes

Synthetic genes were ordered from Genscript Inc. (Piscataway, NJ, USA) or Integrated DNA Technologies, Inc. (Coralville, IA, USA) and cloned in pET29b+ *E. coli* expression vector. Most genes feature a 6-Histidine tag for affinity purification. When protein sequences are in the case of a desired N terminal tag, we add that to the sequence and then add a stop codon at the end of the protein sequence so that the vector tag is not expressed. In some cases, such as the 2 component concentric rings, bicistronic expression was used by including a stop and ribosome binding site between 2 protein sequences included in the gene.

##### Protein production

E. coli expression strain BL21(DE3*) (New England Biolabs, MA, USA) was transformed with plasmid for protein expression. After transformation and overnight growth on LB agar Kanamycin selection plates at either 37°C or 30°C, colonies were picked and transferred to 50mL autoinduction media ^38^ in 250 mL baffled flasks, where they would typically be incubated at 37°C for 16-24 hours. At the end of this incubation period, cells were harvested by centrifuging at 4000xG for 10 minutes at 4°C.

##### Protein purification

Cell pellets were resuspended in 30 mL lysis buffer on ice; this was performed by vortexing and by mixing with serological pipette. This suspension was lysed by sonication (QSonica Sonicators, CT, USA) at 65% power for 3 minutes (15 sec on/15 sec off) with ¾” tips while keeping the cell suspension on ice. Soluble and insoluble lysates were separated by centrifugation at 4°C and 18,000 G for 40 minutes and applied to chromatography columns containing Ni-NTA (Qiagen, MA, USA) resin pre-equilibrated with lysis buffer. The columns were typically washed twice with 10x column volume of lysis buffer, followed by 15mL of elution buffer for protein elution. Concentration in a 10 kDa molecular weight cutoff spin concentrator was performed after this when necessary.

For two-component nanocages, the two components were expressed in separate cell cultures (50mL each) and harvested into the same tube. The mixed cells were then resuspended in 2.66 mL lysis buffer and sonicated at 80% power with ⅛” tips.

In rare cases, such as with the rails of the train track, the insoluble lysate pellet was taken and resuspended in lysis buffer supplemented with 6M guanidine hydrochloride. This suspension can follow the same centrifugation and Ni affinity purification as above, just with the 6M guanidine hydrochloride added to each buffer, which ultimately was dialyzed away overnight into TBS with Slide-A-Lyzer MINI Dialysis Devices (Thermo Fisher Scientific, MA, USA).

Proteins that expressed well were typically further characterized and purified by size exclusion chromatography (SEC). SEC was done on either the Superdex 6 increase 10/300 GL column (Cytiva, MA, USA) or a Superdex 200 increase 10/300 GL column in TBS buffer or high salt TBS buffer, depending on protein/assembly size. Elution profiles were compared amongst each other in batches of designs and to the manufacturer’s provided elution profiles for molecular weight standards.

##### Small Angle X-ray Scattering (SAXS)

Samples were buffer exchanged to TBS+2% glycerol v/v; a blank buffer was obtained by spin concentrator flow-through. SAXS Scattering measurements were performed at the SIBYLS 12.3.1 beamline at the Advanced Light Source. For each sample, data were collected for two different concentrations to test for concentration-dependent effects; ‘low’ concentration samples corresponded to ∼ 1 mg/ml and ‘high’ concentration samples to ∼ 5 mg/ml. Collected data were processed using the SAXS FrameSlice online server and the FoXS software (Sali Lab) was used to compare experimental scattering profiles to design models and assess quality of fit by computing χ2 ^39^.

##### Buffer recipes

Lysis buffer: 25 mM Tris, 25 mM NaCl, 20 mM Imidazole, pH 8.0 at room temperature

Elution buffer: 25 mM Tris, 25 mM NaCl, 300 mM Imidazole, 50mM EDTA, pH 8.0 at room temperature

TBS buffer: 25 mM Tris pH 8.0, 100 mM NaCl

High salt TBS buffer: 25 mM Tris pH 8.0, 300 mM NaCl

##### Negative Stain Electron Microscopy

Samples for nsEM were typically prepared at 0.1 mg/mL concentration for initial screening in a TBS buffer. 5 μL was applied on glow discharged, carbon-coated 400-mesh copper grids (01844-F, TedPella,Inc.), then washed with Milli-Q Water and stained using 0.75% uranyl formate ^40^. Air-dried grids were then imaged on a FEI Talos L120C TEM (FEI Thermo Scientific, Hillsboro, OR) equipped with a 4K × 4K Gatan OneView camera at a magnification of 57,000x and pixel size of 2.47 Å. Micrographs collection was automated using EPU software (FEI Thermo Scientific, Hillsboro, OR) and were imported into cryoSPARC software ^41^. Typically, micrographs were imported with constant CTF and then manual particle picking of 200-400 particles was done to make templates through 2D classing for automated picking. After full scale templated particle picking, additional 2D classing was done, and selected 2D classes were used for C1 (non-symmetrized) ab initio reconstruction followed by either C1 or symmetric homogeneous refinement, depending on what structural features were being analyzed.

##### Cryo-EM sample preparation for sC4, sC4+2, cage_03_10, cage_T3_5, cage_T3_5+2, cage_T3_5+6, and tC3_A

Grids (QUANTIFOIL® R 2/2 on Cu 300 mesh grids + 2 nm C) were frozen using a Vitrobot Mark IV with a chamber maintained at 22°C and 100% humidity. 3.5 μl of protein at 0.5 mg/ml (cage_03_10, cage_T3_5, cage_T3_5+2, cage_T3_5+6), 0.8 mg/ml (sC4, tC3_A), or 1.0 mg/ml (sC4+2) was applied to the surface of a freshly glow-discharged (for 5 s) grid. Grids were then blotted for 3 - 4 s and plunge-frozen in liquid ethane. Due to low particle density in initial cryo-EM screening, cage_03_10 was concentrated ∼4-fold using 10K MWCO Amicon centrifugal filtration devices prior to grid freezing. All grids were screened at the NYU Cryo-EM core facility using a Talos Arctica microscope operated at 200 kV with a Gatan K3 camera.

##### Cryo-EM sample preparation for cage_04_34

3 μL of 1.0 mg/mL of cage_O4_34 in 25mM Tris pH 8.0 300mM NaC was applied to glow-discharged 2/2 Quantifoil carbon grids. Vitrification was performed on a Mark IV Vitrobot with a wait time of 7.5 seconds, with a blot time of 0.5 seconds, and a blot force of either 0 or -1 before being immediately plunged frozen into liquid ethane. The sample grids were clipped following standard protocols before loaded into the microscope for imaging.

##### Cryo-EM sample preparation for strut_C6_21

Cryo-EM grids for strut_C6_21 samples were prepared by diluting protein samples with TBS 1 to 10 times immediately before applying 3.5 μL to glow-discharged 400 mesh, C-flat, 2 micron holes, 2 micron spacing, CF-2/2-4C (CF-224C-100) (Electron Microscopy Sciences, Hatfield, PA) cryo-EM grids. Grids were 10 blotted using a blot force of 0 and 5.5 second blot time at 100% humidity and 4°C and plunge-frozen in liquid ethane using a Vitrobot Mark IV (FEI Thermo Scientific, Hillsboro, OR).

##### Cryo-EM sample preparation for R12B

Four datasets of R12B were collected. For the first two, 3 µL of 10 µM R12B in buffer (300 mM NaCl, 25 mM Tris, pH 8.0) was applied to glow-discharged C-flat 2/2 holey carbon EM grids (Protochips), then blotted and plunge-frozen into liquid ethane using an FEI Vitrobot set to 22 °C with 100% relative humidity. For the third and fourth datasets, 3 µL of 10 µM RK12B in buffer + 0.05% fluorinated octyl maltoside was applied to glow-discharged UltrAuFoil 1.2/1.3 holey gold EM grids (Quantifoil), then blotted and plunge-frozen as above.

##### Cryo-EM data collection, processing and model building of sC4

A total of 3,850 movies were collected with Leginon in super-resolution mode at 0.4124 Å per pixel on a Krios microscope equipped with a K3 camera ^42^. Data collection parameters are provided in Supplemental Table S1 and a data processing workflow is provided in Supplemental Figure S25. Movies were pre-processed (2X binned and motion-corrected with MotionCor2) within Appion, then imported to CryoSPARC v.2 and 3 for further processing ^41, 43, 44^. CTF was estimated in CryoSPARC using CTFFIND ^45^. 3,781,336 particles were picked using templates generated from processing of a subset of micrographs. Iterative 2D classification was performed in CryoSPARC, and a subset of the 2D-curated particles were used for *ab initio* 3D reconstruction. To curate the particle set in 3D, homogeneous refinement was alternated with C1 heterogeneous refinement with 4 classes-one in which the starting model was the best working reconstruction, and three of which were “junk” classes generated by 3D reconstruction of rejected particles. After the final round of homogeneous refinement in CryoSPARC, the particle set was subjected to a final round of 2D classification after which a few particles were excluded, and 1,212,156 particles were imported to Relion v.3 for a final round of 3D classification ^46^. 378,829 particles from the single best class were then refined in Relion using either C4 symmetry or no symmetry (C1). The C4 symmetric map was sharpened using a post-processing job with automatic B-factor assignment. Individual chains of the sC4 design model were docked as rigid bodies into the z-flipped final cryo-EM map using Chimera and imported to Phenix for real-space refinement ^47, 48^. A single round of simulated annealing was used in the first round of refinement, after which rounds of restrained refinement (using secondary structure, non-crystallographic symmetry, rotamer, and Ramachandran restraints) were alternated with manual inspection and adjustments in Coot to generate the final mode_l_^49^. Map-model correlation coefficients were calculated in Phenix and model geometry was analyzed using MolProbity. Sphericity was estimated using independent half-maps from refinement using the 3DFSC server (https://3dfsc.salk.edu/)^50^. Coordinate refinement details are shown in Supplemental Table S3.

##### Cryo-EM data collection and processing of cage_03_10

A total of 4,262 movies were collected with Leginon in super-resolution mode at 0.4124 Å per pixel on a Krios microscope equipped with a K3 camera ^42^. Data collection parameters are provided in Supplemental Table S1 and a data processing workflow is provided in Supplemental Figure S26. Movies were imported into CryoSPARC v2 and 3, motion corrected using Patch motion correction and CTF was estimated using the Patch CTF estimation job ^41^. After visual curation and removal of 87 micrographs, a subset of micrographs (867) was used to pick particles, both manually and using a blob picker. The reference model for homogeneous refinement used was generated using *ab initio* 3D reconstruction and generated with O symmetry from a subset of 24,040 particles picked from 867 micrographs. After one round of 2D classification using images binned 4 times, the best templates were selected and used as inputs of template picker (663,778 particles picked) and topaz picker (1,336,640 particles picked) to re-pick particles on all of the micrographs. After several rounds of 2D classification and removal of duplicate particles, we obtained 86,379 particles. An initial round of 3D homogenous refinement without symmetry (C1) using all images binned 4 times was performed, using the *ab initio* model as a reference. Following homogeneous refinement, particles were subjected to heterogenous refinement with 3 classes, using 2 decoy references for further sorting of heterogeneity. Particles from 2 of the resulting 3 classes were removed, and the 50,540 particles were then refined using unbinned images. Two rounds of non-uniform refinement in both C1 and O symmetries were performed. Overall resolution was estimated using CryoSPARC implementation of the gold standard method, from which the average reported resolutions are 7.4 and 6.0 respectively. The design model was docked into the final map in Chimera to assess fit of the cryo-EM map to the design model, and analyze the design model-to-map fit ^47^. Sphericity was estimated using independent half-maps from refinement using the 3DFSC server (https://3dfsc.salk.edu/).

##### Cryo-EM data collection and processing of cage_T3_5

A total of 5,854 movies were collected with Leginon in super-resolution mode at 0.4124 Å per pixel on a Krios microscope equipped with a K3 camera ^42^. Data collection parameters are provided in Supplemental Table S1 and a data processing workflow is provided in Supplemental Figure S27. Movies were imported into CryoSPARC, motion corrected using Patch motion correction and CTF was estimated using the Patch CTF estimation job ^41^. After visual curation and removal of 118 micrographs, a subset of micrographs (1,236) was used to pick particles, both manually and using a blob picker. After one round of 2D classification using images binned 4 times, the best templates were selected and used as inputs of template picker (2,585,392 particles picked) and topaz picker (1,800,346 particles picked) to re-pick particles on all of the micrographs. After several rounds of 2D classifications and removal of duplicate particles, the resulting 959,145 particles were subject to another round of 2D classification. 676,480 particles from the selected 2D classes were used to generate an *ab initio* model with T symmetry. Homogeneous refinement was performed without symmetry (in C1), using the *ab initio* model as a reference, with 4 times binned images.. Heterogeneous refinement was performed with 3 decoy classes, resulting in 371,057 particles classified into good classes. One round of homogenous refinement and one round of non-uniform homogenous refinement were performed using unbinned images. A final round of heterogeneous refinement was run, resulting in the final particle set. 144,976 particles were refined in C1 symmetry and T symmetry, leading to the final maps, with reported average resolutions of 4.3 Å in C1 symmetry, and 3.6 Å with a T symmetry (class 0 in workflow on Supplemental Figure S27). Class 1 was not an identifiable shape, while classes 2, 3, 4 and 5 appear to be very low resolution reconstructions of the target design, in which part of the intended assembly is missing. Class 3 (116,923 particles) was subjected to another round of heterogeneous refinement leading to two maps, Map 3.0, with a reported average resolution of 6.1 Å, and Map 3.1, with a reported average resolution of 6.5 Å. The design model was docked into the final map in Chimera to assess fit of the cryo-EM map to the design model, and analyze the design model-to-map fit ^47^. Map 3.0 clearly shows that the density for one monomer is absent, while Map 3.1 clearly shows that the density for one trimer is absent. Sphericity was estimated using independent half-maps from refinement using the 3DFSC server (https://3dfsc.salk.edu/).

##### Cryo-EM data collection and processing of cage_T3_5+2

A total of 19,358 movies were collected with Leginon in 3 sessions, in super-resolution mode at 0.4124 Å per pixel on a Krios microscope equipped with a K3 camera ^42^. Data collection parameters are provided in Supplemental Table S1 and a data processing workflow is provided in Supplemental Figure S28. Movies were aligned using Relion v3 ’s implementation of the motion correction algorithm before being imported into CryoSPARC ^41, 46^. After patch CTF estimation and curation, a subset of 436 micrographs were used to pick particles, both manually and using blob-picker. After one round of 2D classification with images binned 4 times, the best classes were selected as templates to feed into a template picker and topaz picker jobs to pick particles on all of the micrographs. After removal of duplicates, the resulting 1,318,959 particles were subjected to several rounds of 2D classification. A subset of 363,381 particles was used to generate *ab initio* models with T symmetry. After an initial round of homogeneous refinement without symmetry (C1), further classification was performed using heterogeneous refinement with decoy references to exclude bad particles. The resulting 906,447 particles were refined using homogenous and non-uniform refinement without symmetry (C1). Another round of heterogeneous refinement into 6 classes led to 4 classes which resembled part of the design target, but with parts of the expected target missing. The remaining 2 classes did not resemble any expected shape, and are likely junk particles. The highest resolution map (38% of the particles) was refined without symmetry (C1) leading to a map with reported average resolution of 6.7 Å. The design model was docked into the final map in Chimera to assess fit of the cryo-EM map to the design model, and analyze the design model-to-map fit ^47^. Sphericity was estimated using independent half-maps from refinement using the 3DFSC server (https://3dfsc.salk.edu/).

##### Cryo-EM data collection and processing of sC4+2, tC3_A, cage_T3_5+6

All movies from these datasets were collected with Leginon as noted in Supplemental Table S1 ^42^, and movies were aligned using Relion’s implementation of the motion correction algorithm before being imported into CryoSPARC ^41, 46^. CTF estimation was performed using the Patch CTF estimation job. Several rounds of 2D classification were performed until a clean stack of particles was obtained.

##### Cryo-EM data collection and processing of cage_O4_34

Data collection was performed automatically using EPU (FEI Thermo Scientific) to control a ThermoFisher Tundra 100 kV TEM equipped with a standalone CETA-F direct electron detector. Data were collected using fractionation mode, with random defocus ranges spanning between - 0.5 and -2.2 μm using image shift and multiple shots per hole. Two sets of movies (3,226 and 1,324) were collected with a pixel size of 1.248 Å.

All data processing was carried out in CryoSPARC ^41^. Alignment of movie frames was performed using Patch Motion with an estimated B-factor of 500 Å2, with a maximum alignment resolution set to 5 Å. Defocus and astigmatism values were estimated using Patch CTF with Amplitude Contrast set to 0.07. 252 particles were initially picked using Manual Picker and extracted with a box size of 380 pixels. An initial round of reference-free 2D classification with 10 classes was performed with a maximum alignment resolution of 6 Å, resulting in classes representing views of all three symmetry axes. These classes were used as input for a round of Template Picking with particle diameter of 342 Å, initially picking 248248 particles (before extraction) which were extracted with 380 pix boxes. A round of 2D classification was performed and the particles were sorted into 100 classes. Classes which clearly resolve the cage particles were selected and 59904 particles were used to perform 3D ab initio reconstruction consisting of 3 classes with a maximum alignment resolution of 12 Å using O symmetry. The largest class was refined using Non-uniform Refinement using all particles and using O symmetry with per-particle defocus optimization to arrive at a 7.5 Å map. Viewing Direction Distribution had particle clusters separated by 90 degrees corresponding to slight preferred orientations for the face of the cube. 3D maps for the half maps, final unsharpened maps, and the final maps sharpened by DeepEMhancer ^51^ (tightTarget model) were deposited in the EMDB under accession number EMD-29915. The processing pipeline for this design is illustrated in Supplement Figure S29, and image processing details are provided in Supplemental Table S4. Figures were generated using UCSF ChimeraX ^52^.

To validate the use of O-symmetry during reconstruction and refinement, we performed an independent round of Ab-initio reconstruction and Heterogeneous Refinement of 12 classes with C1 symmetry. The 3 classes containing fully resolved cages were subjected to another round of reconstruction Heterogeneous Refinement with all particles, followed by Non-Uniform Refinement to arrive at a 8.7 A map which overlaps well with the O4 map.

##### Cryo-EM data collection and processing of strut_C6_21

Strut_C6_21 cryo-EM grids were screened and data was collected on a Glacios transmission electron microscope (FEI Thermo Scientific, Hillsboro, OR) operated at 200 kV and equipped with a Gatan K3 Summit direct detector. Data collection was performed automatically using SerialEM to control a ThermoFisher Glacios 200 kV equipped with a standalone K3 Summit direct electron detector with a nominal 54 magnification of 36,000x (0.883 Å/pixel). Movies were acquired in counting mode fractionated in 50 frames of 200 ms at 8.5 e-/pixel/sec for a total dose of ∼65e-/Å2

All data processing was carried out in CryoSPARC ^41^. Alignment of movie frames was performed using Patch Motion with an estimated B-factor of 500 Å^2^, with a maximum alignment resolution set to 3. Defocus and astigmatism values were estimated using Patch CTF with default parameters. Strut_C6_21 particles were initially picked in a reference-free manner using Blob Picker and extracted with a box size of 450 pixels. This was followed by multiple rounds of 2D classification and subsequent template-picking using the best 2D class averages. The best classes that revealed clearly visible secondary-structural elements, a total of 37.105 particles, were used for 3D ab initio determination using the C1 symmetry operator. This was followed by a 3D non-uniform refinement with C6 symmetry, global CTF refinement and local refinement for a final global resolution estimate of 5.12 Å. Local resolution estimates were determined in CryoSPARC using an FSC threshold of 0.143. 3D maps for the half maps, final unsharpened maps, and the final sharpened maps were deposited in the EMDB under accession number EMD-29893. The processing pipeline for this design is illustrated in Supplemental Figure S30.

For the model shown in Figure 3C, the design model was fit and refined to the C6 experimental map density using the final model after Phenix refinement with Namdinator ^53^.

##### Cryo-EM data collection and processing of R12B

High-throughput data collection was performed with a Gatan K3 Summit direct electron detector on an FEI Glacios Cryo TEM operating at 200 kV accelerating voltage, controlled by SerialEM software ^54^. Datasets 2 and 4 were collected with 40° and 30° stage tilt, respectively.

Movies were collected in super-resolution mode, then aligned and corrected for full-frame motion and sample deformation with the patch motion correction algorithm in CryoSPARC v4, with 2x Fourier binning and dose compensation applied during motion correction. All initial processing was performed in CryoSPARC v4 ^41^. Contrast transfer function (CTF) was estimated with the patch CTF estimation algorithm, and automatic particle picking using Blob Picker was used to generate template 2D classes for template-based autopicking. Picked particles were boxed and extracted with Fourier cropping and 2D classification was performed iteratively to generate sets of quality particles from all four datasets, which were then combined and run through one more round of 2D classification to select the best-resolved classes. These particles were then re-extracted without Fourier cropping for further processing.

Ab-initio reconstruction was used to generate four volumes and associated classes of particles. The best volume and particle class were used for non-uniform refinement with C12 symmetry to generate an initial map. Local CTF refinement was performed and particles were used for another round of non-uniform refinement with C12 symmetry. This particle stack was converted to a .star file for processing in RELION 4.0 using the csparc2star.py program in the UCSF pyem collection ^55^.

Remaining processing was performed in RELION 4.0 ^56^. The particle stack imported with csparc2star was reconstructed using relion_reconstruct, and this volume was used as a reference for a 3D auto-refine with no symmetry imposed. Symmetry expansion (relion_particle_symmetry_expand) was used to generate a particle stack expanded with C4 symmetry, which were run through 3D classification without alignment with the sharpened map from 3D auto-refine as a reference. Best-resolved 3D classes were selected, duplicates from symmetry expansion were removed, and another round of 3D auto-refine with no symmetry imposed was performed with the RELION-generated C1 map and a mask generated from the C1 map as references. 3D auto-refine of this new C1 map with C12 symmetry imposed generated a 5.3 Å map. Higher-order aberrations and anisotropic magnification were estimated, and per-particle defocus values and per-micrograph astigmatism were optimized using CTF refinement, which improved the map to a final resolution of 5.2 Å. The processing pipeline for this design is illustrated in Supplementary Figure S31.

##### Crystallization and Structure Determination

All crystallization experiments were conducted using the sitting drop vapor diffusion method. Crystallization trials were set up in 200 nL drops using the 96-well plate format at 20 ℃.

Crystallization plates were set up using a Mosquito LCP from SPT Labtech, then imaged using UVEX microscopes and UVEX PS-256 from JAN Scientific. Diffraction quality crystals formed in 0.2 M Magnesium chloride hexahydrate, 0.1 M BIS-TRIS pH 5.5, and 25% w/v Polyethylene glycol 3,350 for THR1; in 0.3M Sodium nitrate, 0.3M Sodium phosphate dibasic, 0.3M Ammonium sulfate, 0.1M Sodium HEPES; MOPS(acid) pH 7.5, 30% mixture of 40% v/v Ethylene glycol; 20% w/v PEG 8000 for THR2; in 0.2 M Ammonium acetate, 0.1 M HEPES pH 7.5, 25% w/v Polyethylene glycol 3,350 for THR5 and in 0.1M mixture of 0.2M DL-Glutamic acid monohydrate; 0.2M DL-Alanine; 0.2M Glycine; 0.2M DL-Lysine monohydrochloride; 0.2M DL-Serine, 0.1 M mixture of 1M Imidazole; MES monohydrate (acid) pH 6.5, 30% mixture of 40% v/v Ethylene glycol; 20% w/v PEG 8000 for THR6.

Diffraction data was collected at the Advanced Light Source (ALS) 8.2.1/ 8.2.2/ 5.0.2. X-ray intensities and data reduction were evaluated and integrated using XDS and merged/scaled using Pointless/Aimless in the CCP4 program suite ^57, 58^. Structure determination and refinement starting phases were obtained by molecular replacement using Phaser ^59^ using the designed model for the structures. Following molecular replacement, the models were improved using phenix.autobuild ^60^; efforts were made to reduce model bias by setting rebuild-in-place to false, and using simulated annealing and prime-and-switch phasing. Structures were refined in Phenix ^60^. Model building was performed using COOT ^61^. The final model was evaluated using MolProbity ^62^. Data collection and refinement statistics are recorded in Table S5. Data deposition, atomic coordinates, and structure factors reported in this paper have been deposited in the Protein Data Bank (PDB), http://www.rcsb.org/ with accession code 8G9J (for THR1), 8G9K (for THR2), 8GA7 (for THR5), 8GA6 (for THR6).

#### Visualization and figures

All structural images for figures were generated using PyMOL (The PyMOL Molecular Graphics System, Version 2.0 Schrödinger, LLC) or ChimeraX ^52^. Figures (cartoons and diagrams) were made in InkScape.

#### Supplementary discussion

##### Additional polygonal oligomer design details

For the ***Turn*** module-based oligomer designs, a confounding factor for design successes across the different polygonal shapes is that using a hinge helix to produce a sharp turn ends up creating corners with different amounts of local helical interaction density as a function of the turn angle. This gave the n=3 triangles very rigid corners, but the n=5 and n=6 designs were much more hinge-like at the turn helix. We obtained one n=5 design that showed pentagon shapes in nsEM characterization, although the lower-order n=4 square shape was erroneously prevalent in this design (Supplemental Figure S1F; protein *72_C5_A*). We did not obtain a n=6 hexagon design with this ***Turn*** method, presumably because the hinge region ended up too un-reinforced. So, we made additional hexagon designs that instead were docked to provide increased helical packing density at the corners. We were able to observe 1 hexagon design from an order of 6 that produced the expected hexagonal shape in nsEM as the dominant species (Supplemental Figure S1G; protein *hex_C6*). Thus, we suggest utilizing this angle-encoding ***Turn*** feature with considerations for local helical density at hinge regions. There are some unique THRs with 3 helices in the repeat unit (Supplemental Figure S5C) which can offer unique looping possibilities for different outcomes after using a ***Turn*.**

There are additional cyclic oligomer designs made with Rosetta FastDesign and Rosetta fragment-biased forward folding that are quite different from what we would order today with proteinMPNN and AlphaFold2. Notably, some novel straight-helix heterodimers (SHDs) which were designed to help make cyclic oligomers in Supplemental Figure S1 (proteins *tC3*, *sC4,* and *sC4+6*) were previously much more prevalent in our building block database (details on construction below). These and also many older THR designs ended up being dropped out of our databases due to new structure prediction capabilities suggesting that they were not nearly as straight as designed. Now, we are much more confident in our capability to create databases of structures that are predicted to be the actual correct shape. The experimental experiences that most closely resemble what a current-day (May 2023) researcher would encounter when designing and testing designs from planned straight helix placement would most likely be similar to “Curve THR rings” (both 18 and 30 repeat experiments) as summarized in Supplemental Figure S2.

We used the SHD blocks to construct closed oligomers by choosing cutpoints on the THRs and the heterodimers that generate the required overall rotation around the Z axis (in contrast to the triangular and square oligomers in Figure 2C, the interfaces between chains have geometries distinct from the interfaces within chains and exhibit more variations in shape) (5). Cryo-EM data for 3 designs yielded 2D class averages in close agreement with the design models (Supplemental Figure S1G, proteins *tC3*, *sC4,* and *sC4+6*). The top-down views of the particles show clear densities for the helices, which are very close to the designed placements of helices in the X-Y plane. The distinctive designed interior corner angles for each of these polygon-like structures are evident, 90° for the C4 “square” and 60° for the C3 “hexagon”. The C3 “hexagon” has two unique corners that both form similar angles; one at the N terminal fusion of the THR to one half of the SHD, and the other at the C terminal fusion of the THR to the other half of the SHD. The C3 “hexagon” design *tC3* yielded 2D class averages that showed the helices in expected placement (Fig 2E), and the C4 “square” sC4 yielded a 4 Å resolution 3D reconstruction in C4 symmetry with 1.6 Å backbone RMSD to the design model, maintaining the characteristic straightness and phases of all helices (Fig 5A, S9).

##### Using concentric ring strutting to rescue assembly behavior

When we were planning our designs for ring-strutting (Figure 3) we picked some with imperfections that could hopefully be rescued by reinforcement. For example, we sought to use a correctly sized monodisperse 20-repeat ring to reinforce a 30-repeat ring which had failed to express solubly (fig. S8A). Both rings were cut into ten subunits, the rotation and Z displacement of one ring relative to the other was sampled, and linear THRs were placed to connect the inner and outer rings. The resulting single component C10 strutted assembly formed 10 and 11-mer species in nsEM 2D classes with the outer ring clearly present, and thus rescued to large extent (Fig. 3A, S8A). In a subset of the classes, a segment of the outer ring was missing, suggesting a distal part of one chain may flay out to alleviate strain. We next used a monodisperse 18-repeat ring to support a 30-repeat ring which on its own formed oblong closed rings often much larger than the design (20-25% increase, fig. S8B). Both were cut into six subunits (with 5 and 3 repeats per subunit, respectively), and the two subunits were connected by a THR strut. The resulting 2 component strutted assembly was monodisperse and close to the design model by nsEM (Fig. 3B, S8B). Dominant 2D class averages showed both rings with all chains present, without the irregular oblongness that was present in the original 30-repeat ring. A 5.1 Å cryo-EM reconstruction was obtained for this strutted ring sample, which showed that at the regions closest to the strut, the outside diameter was only 2% increased compared to the ideal design model (19.7 nm vs 20.1 nm, fig. S8C). The helix positioning in the inner ring and the strut were close to the design model (fig. S8C inset boxes). These results highlight the utility of modular interactions for installing reinforcements that can improve the assembly of nanomaterials.

##### Additional single-component expandable nanocage designs

If the repeat protein propagation axis is not perfectly aligned in the plane of the symmetry axes (or if the repeat protein is not truly linear), upon extension two outcomes are possible: the building blocks could either flex to compensate, or if the strain in flexing is too large, the assemblies might only partially form. To explore the flexibility of the building blocks, and to investigate whether symmetry could be broken by introduction of strain, we generated designs with varying amounts of repeat propagation axis deviation by docking ^8^ THRs that were fused to existing cyclic oligomers ^6^ (fig. S1H, S15) into polyhedral architectures. Of the 12 one-component T3 and O3 nanocages that were tested, cryo electron microscopy maps of a T3 and an O3 design at 4Å and 6Å resolution were very close to the design model, where the helices of the building blocks are clearly visible and aligned as designed. For these designs, based on the design model, the deviations of the linear THR from the plane spanned by the symmetry axes are approximately 8° (fig. S16, S17). Following addition of one or two repeat units, higher order assembly was still observed, but SEC followed by cryo-EM analysis showed that most of the particles do not contain all expected oligomeric subunits (fig. S17). The *cage_T3_5* expansion series showed clear particles, where *cage_T3_5+2* and *cage_T3_5+6* are largely incomplete and are missing a trimer unit when viewed from the 3-fold axis. C1 3D reconstructions showed that a significant population of the particles are missing one trimer. Rigid body fitting of the design model into the *cage_T3_5+2* reconstruction suggests that the trimer itself is quite rigid, matching well into the map. The three trimer units in turn, to compensate for the increase in length of the THR, “open up” thus preventing the capping fourth trimer from joining the assembly. These results suggest that the angle deviation of the THR propagation axis to the required plane is quite strict, and the flexibility of the well-packed designed protein to compensate for angular deviations is relatively low. This opens an exciting possible design strategy for generating asymmetric protein assemblies with unique subunits to fill the now-empty slot for additional functionality, analogous to the single phage tail-spikes emanating from otherwise symmetric icosahedral phages ^63^ (fig. S18).

For the C2 “handshake” based designs (Figure 4), variants of the O3, I3, and O4 cages were tested for expandability (no expanded T3 designs were ordered, just because they were deemed less interesting due to being lower order). While the O4 (*cage_O4_34*) worked remarkably well at both the base size and the variant sizes as discussed in main text (Figure 5B), the O3 and I3 cages that were shown in Figure 4 were not as monodisperse in the base size as the O4 cage (Supplemental Figure S13), and upon extension the off-target species became more prevalent. For all of the handshake cages, there is plenty of room left in the sampling space, even with our tight constraints, so it should be very possible to design and filter more C2 handshake angle designs to allow characterization of more designs to select a subset that are similarly robust to *cage_34_O4* to offer more starting places for successful modular expansions.

##### Additional two-component docked nanocage designs for expandability

From a set of two-component docked O43 nanocage designs for the expandability goal, four designs showed monodisperse particles by negative stain EM and yielded 3D reconstructions in octahedral symmetry that show the key features of the assembly including the pores and extended arms (fig. S19-23). For cage_O43_54 and cage_O43_59, parts of the arms formed by the THR are slightly rotated compared to the design model. Deletion of 4 or addition of 4 or 8 repeat helices from the THRs yielded assemblies in some but not all cases. EM 3D reconstruction of cage_O43_54_-4 and cage_O43_59_+4 yielded maps very similar to the original, where the rotations of both the tetramer and trimer components stay consistent and only the THR portion extends, causing both components to displace further along their respective symmetry axes as expected (fig. S19, S20). To avoid alternative start sites, cage_O43_59 also tolerates mutations to remove methionine residues in the THR (fig. S20). Other sizes of these two cages failed to yield cage-sized assemblies as determined by SEC. An additional cage, cage_O43_164 yielded cages that closely resembled the design model by negative stain EM (fig S23A), but extended variants of the C4 component hosting the THR resulted in insoluble protein. In efforts to solubilize the component, “cut” variants were tested where the first five helices were expressed as a separate protein chain (fig S23B). Cut variants cage_O43_164_cut and cage_O43_164_+2_cut yielded uniform 3-component nanocages when purified via IMAC (fig S23A). The cage_O43_164_cut design matches the parent design as expected, while cage_O43_164_+2_cut matches the expected larger size. The cage_O43_164_+2_cut requires assembly in 2M GuHCl to solubilize the extended component, while additional extension variants remained insoluble.

## Supplementary figures

**fig. S1.**
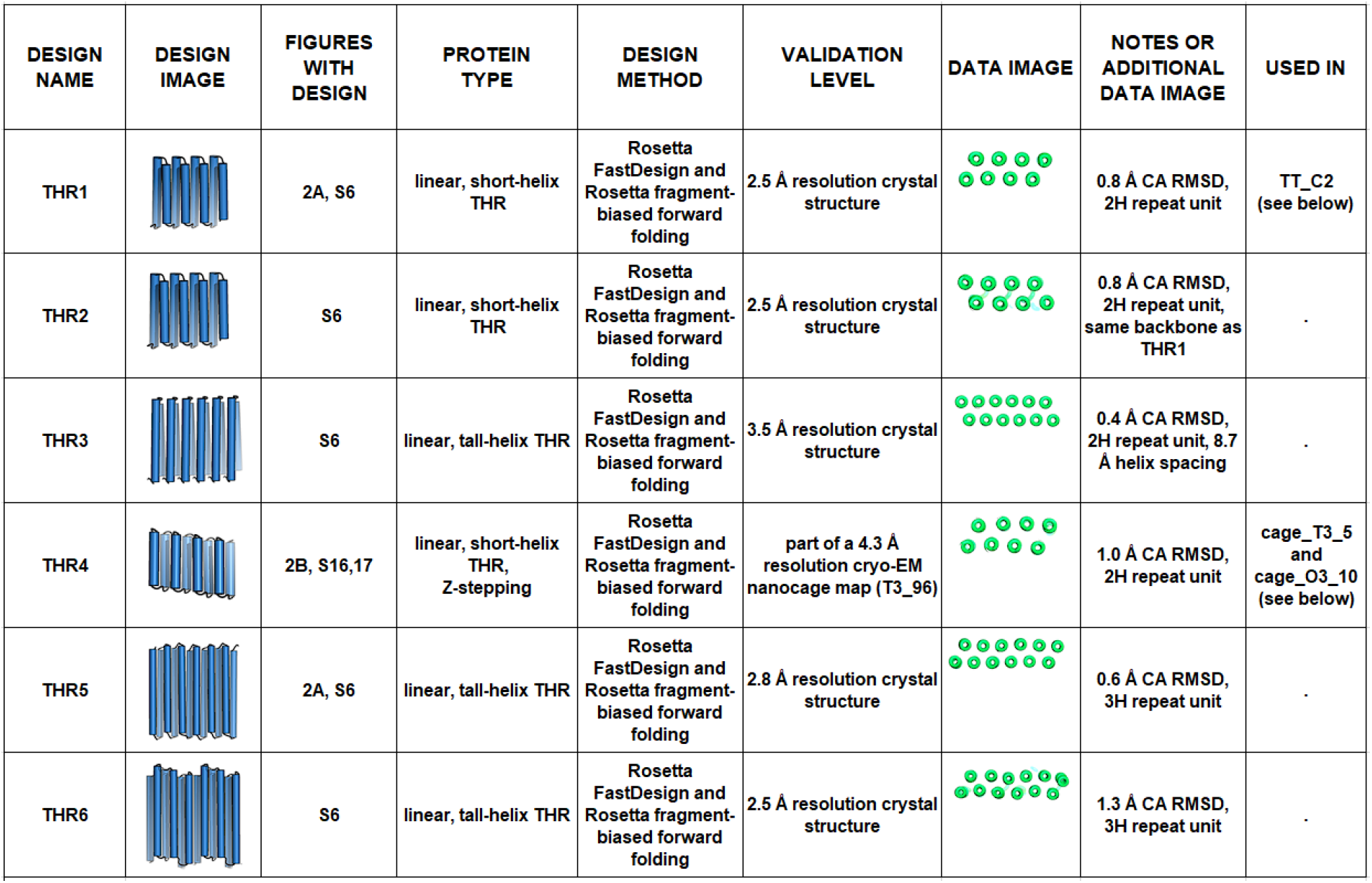
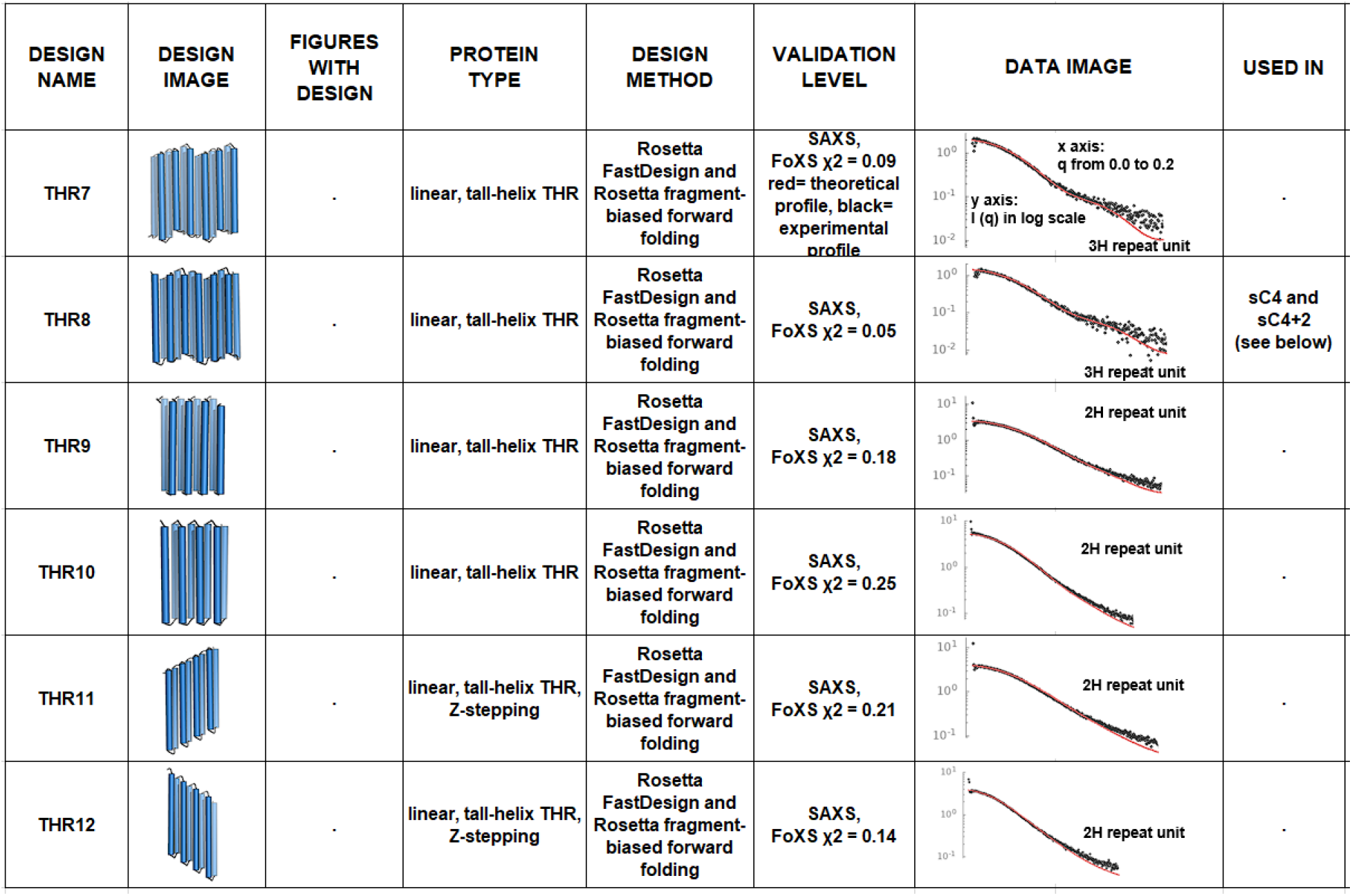
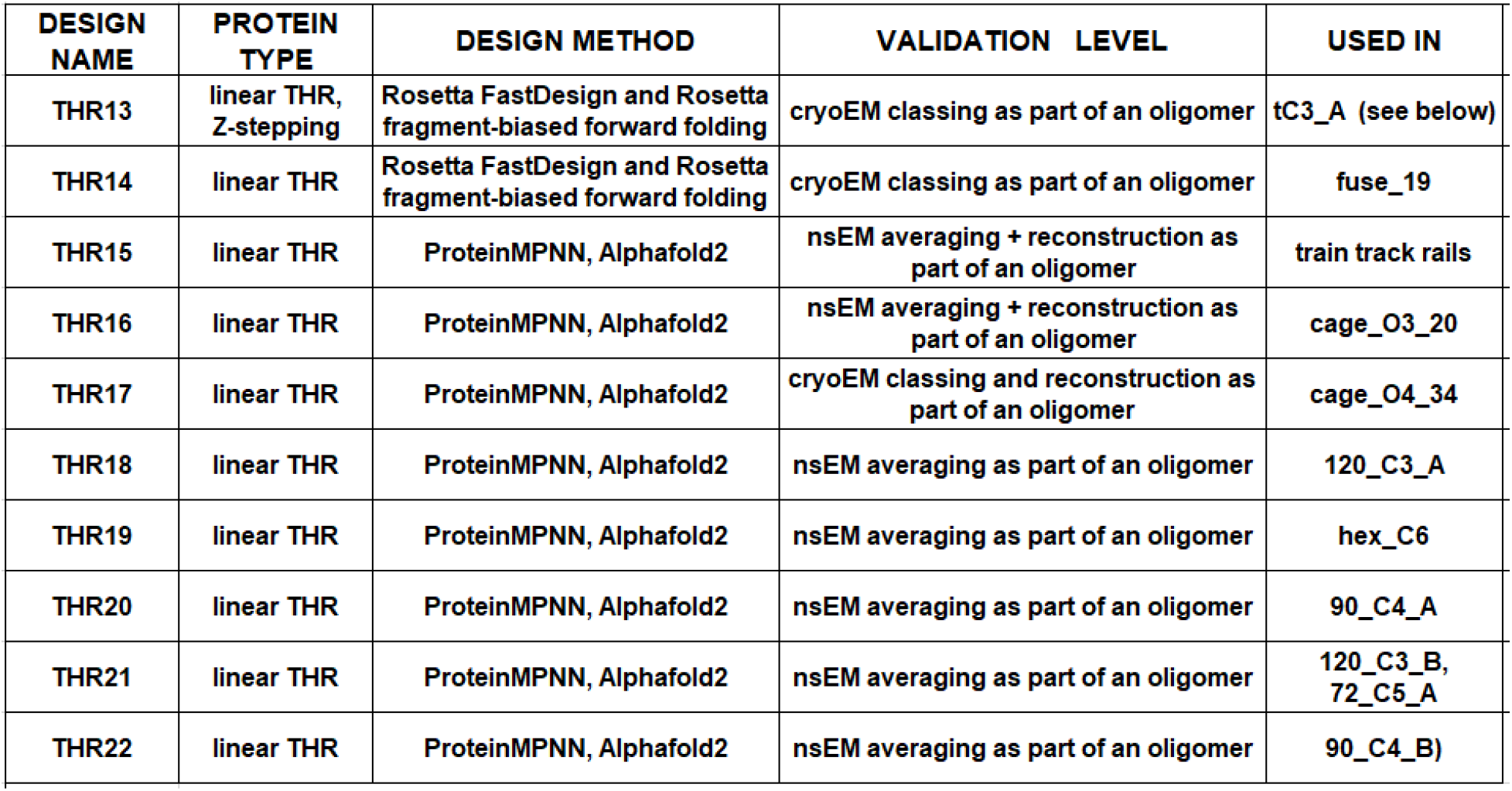
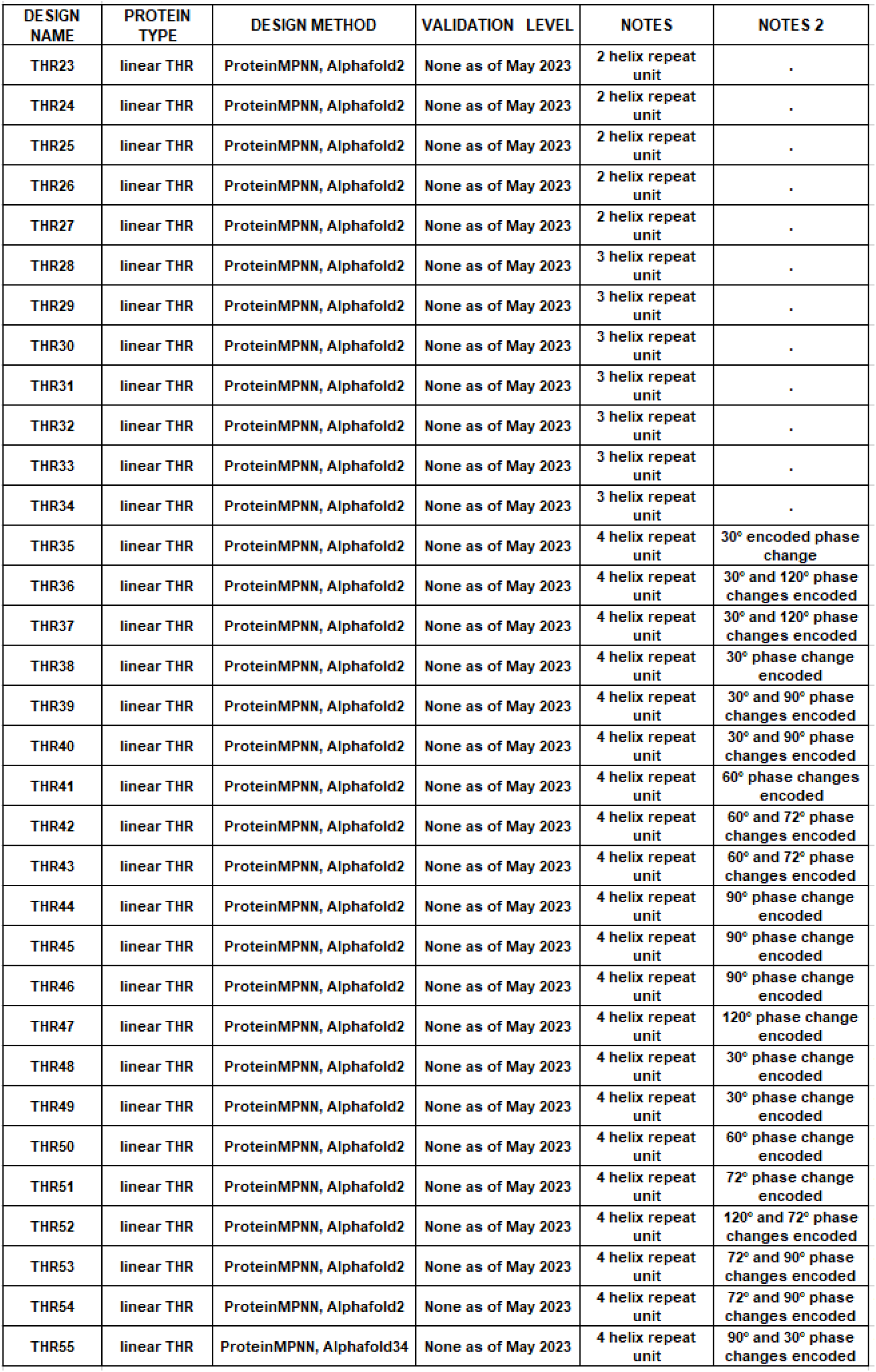
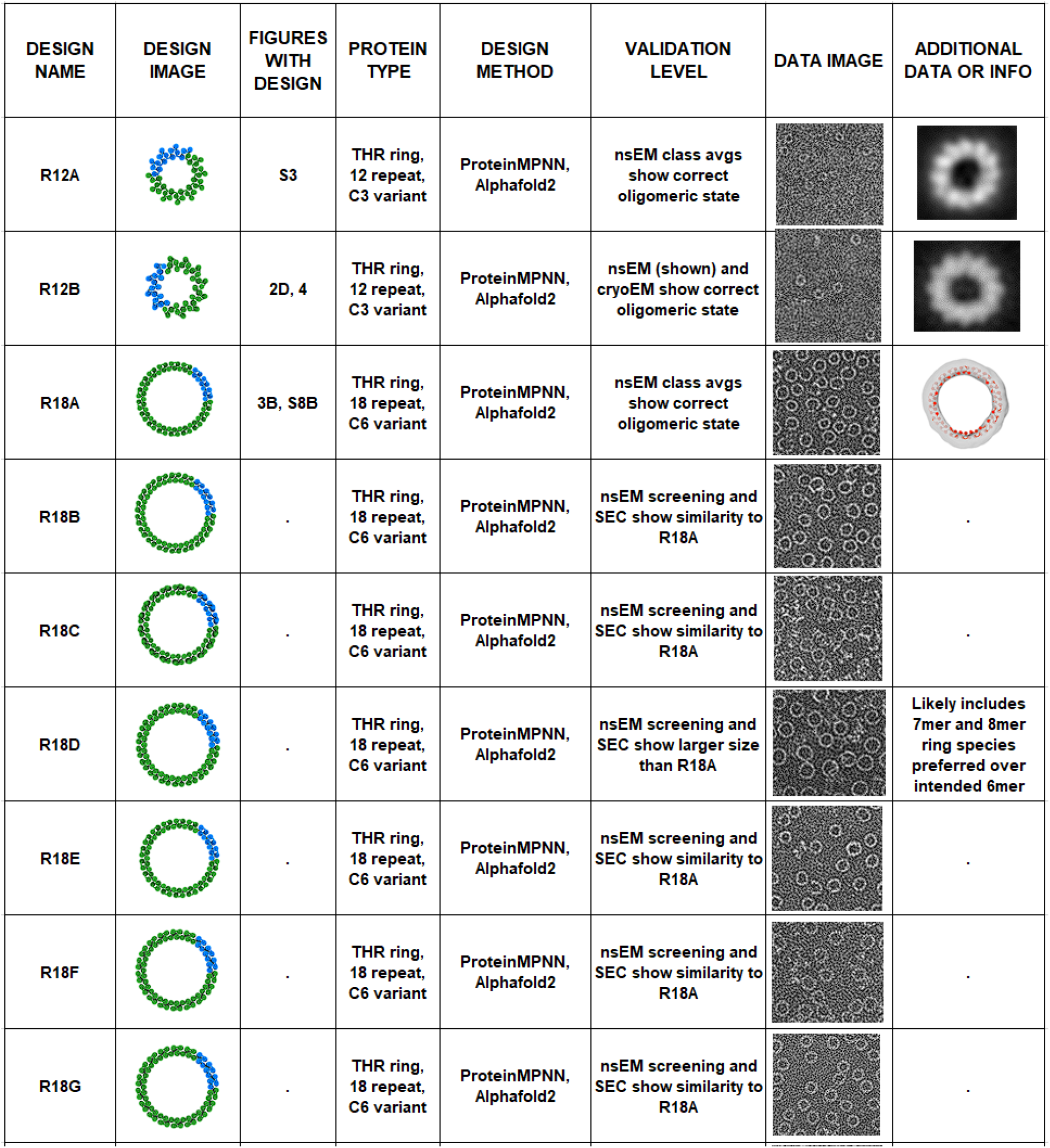
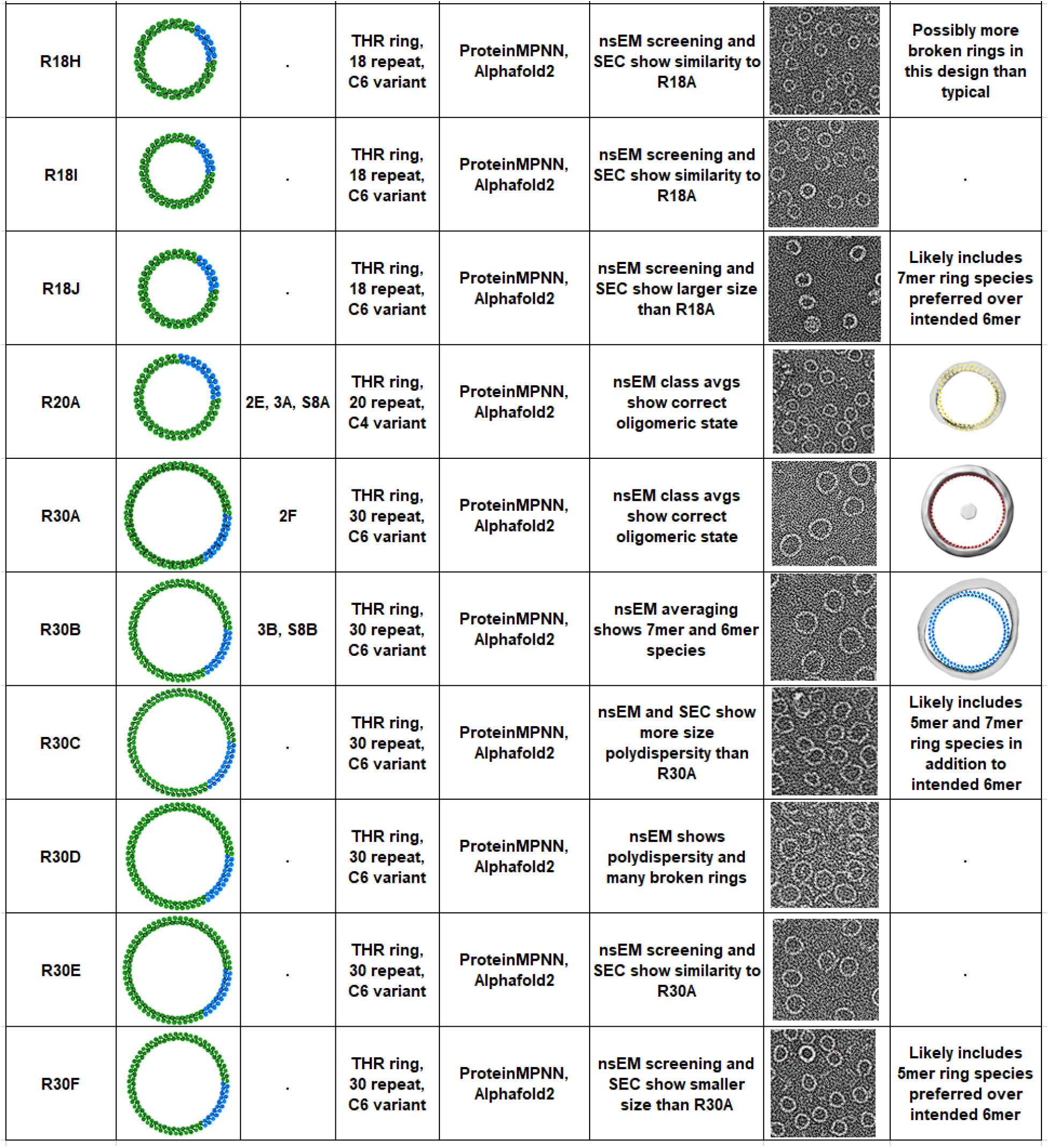
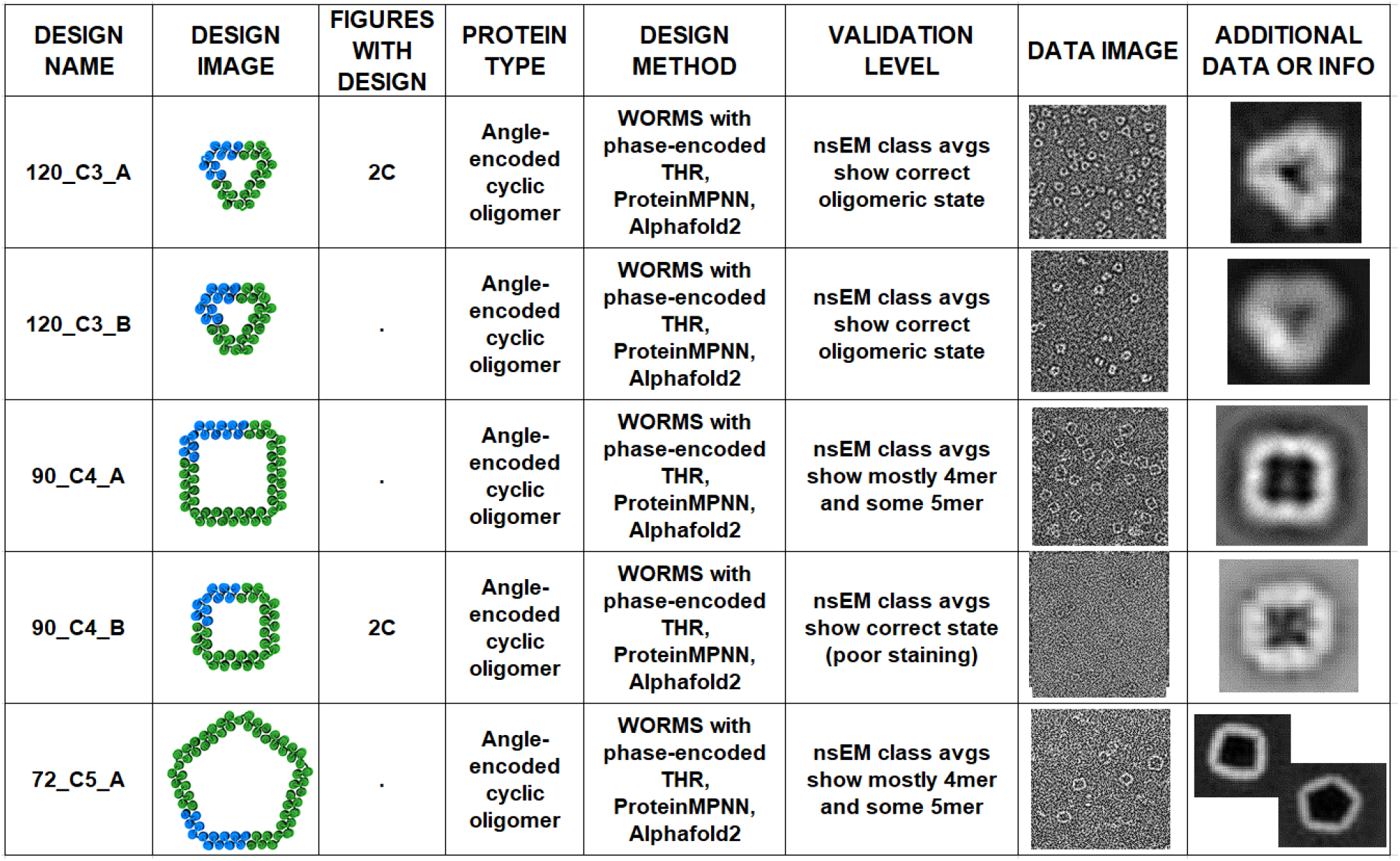
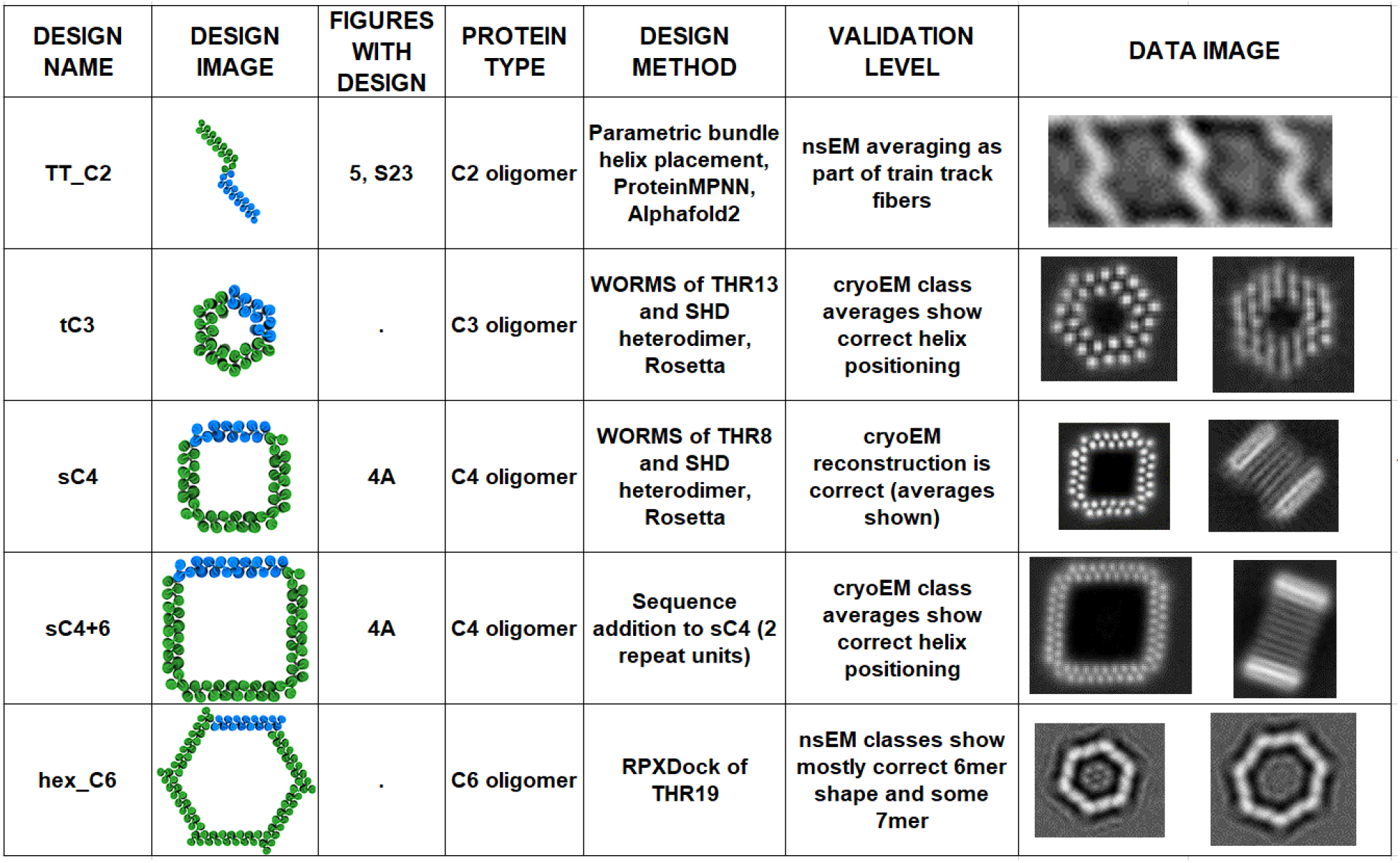
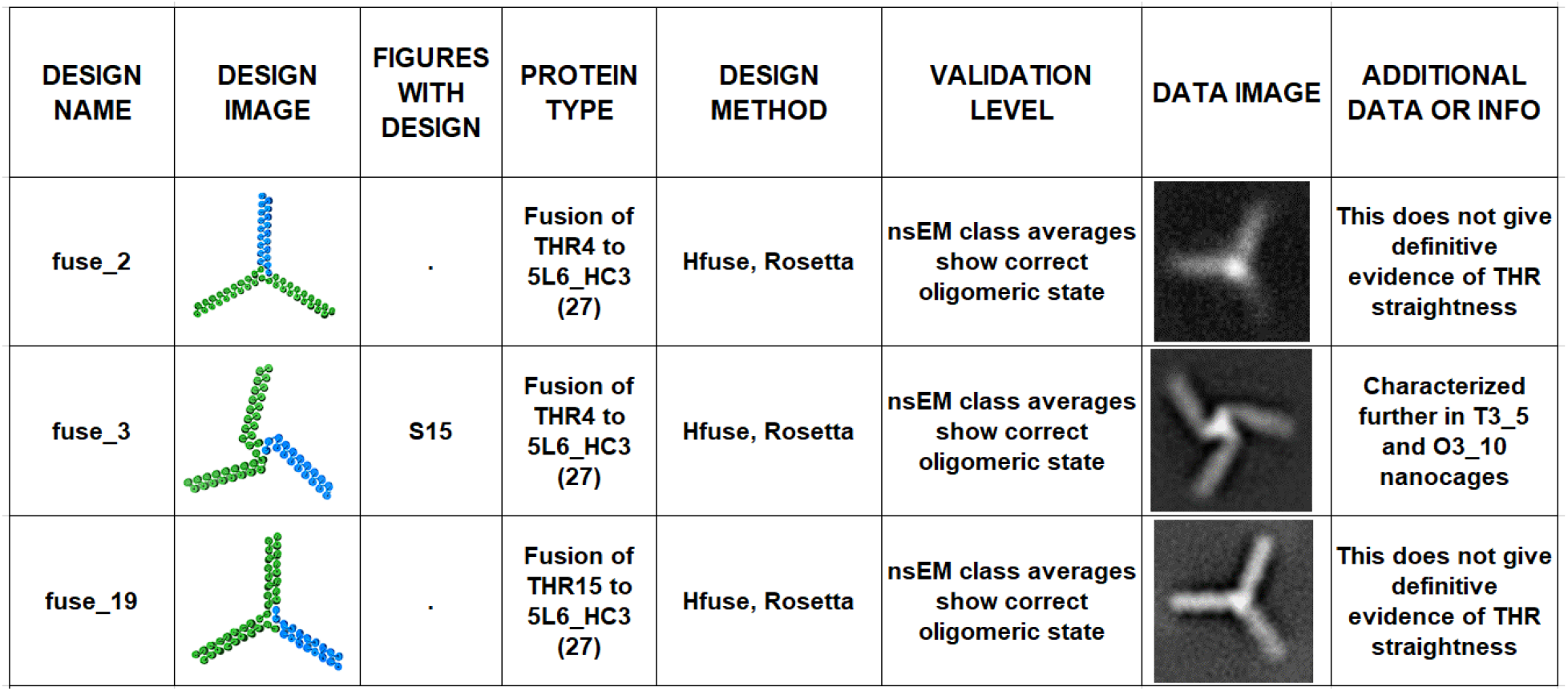
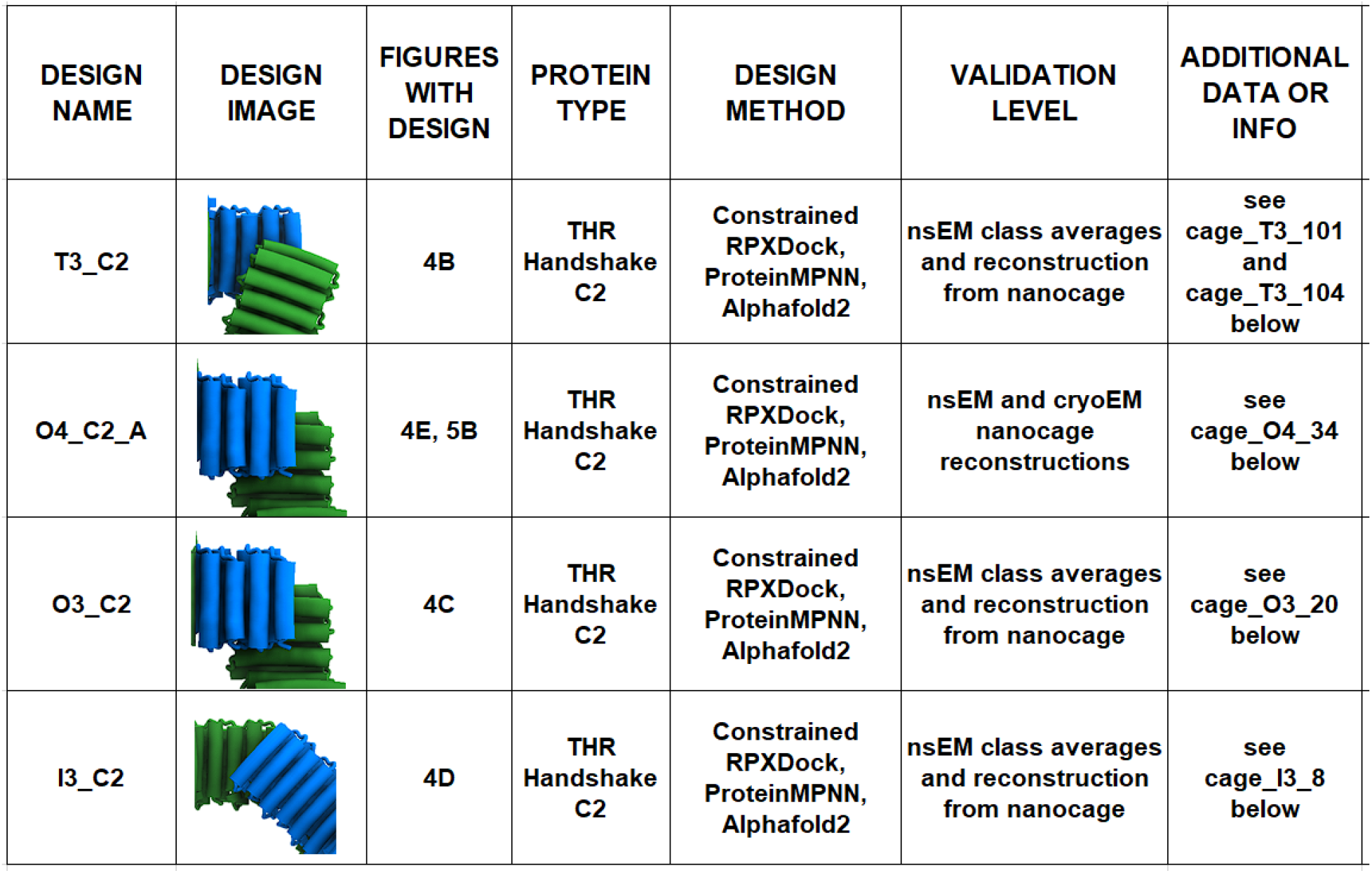
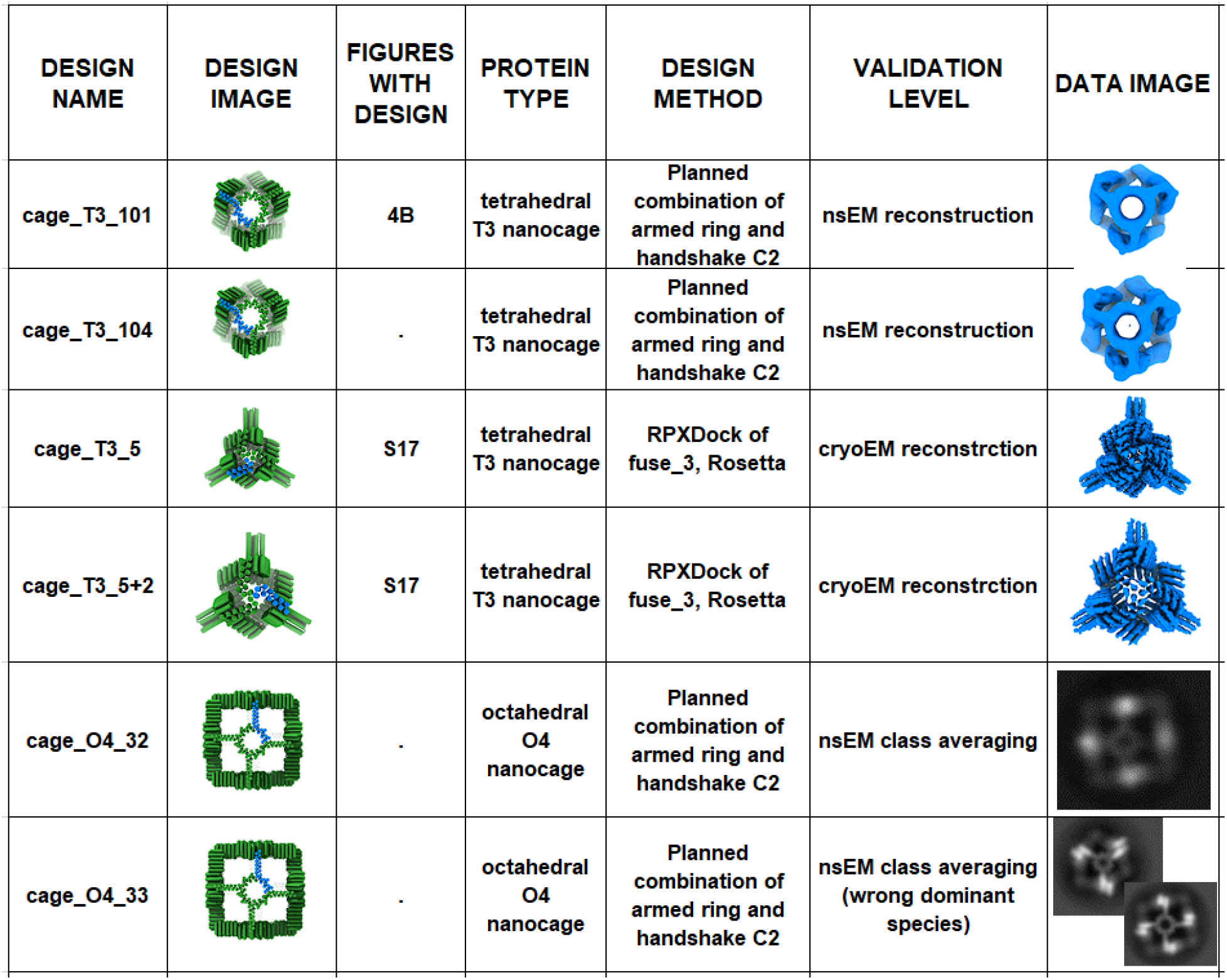
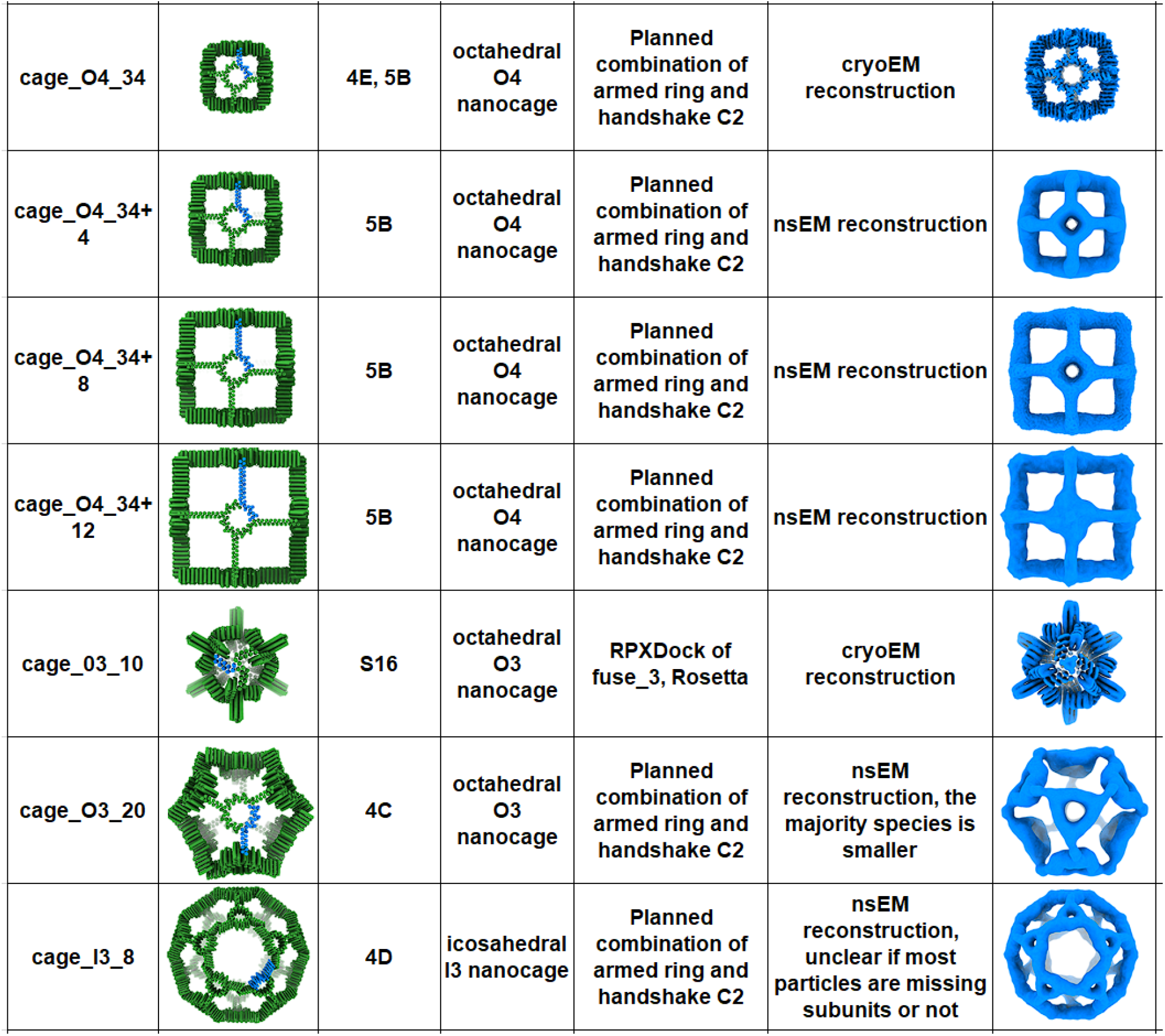
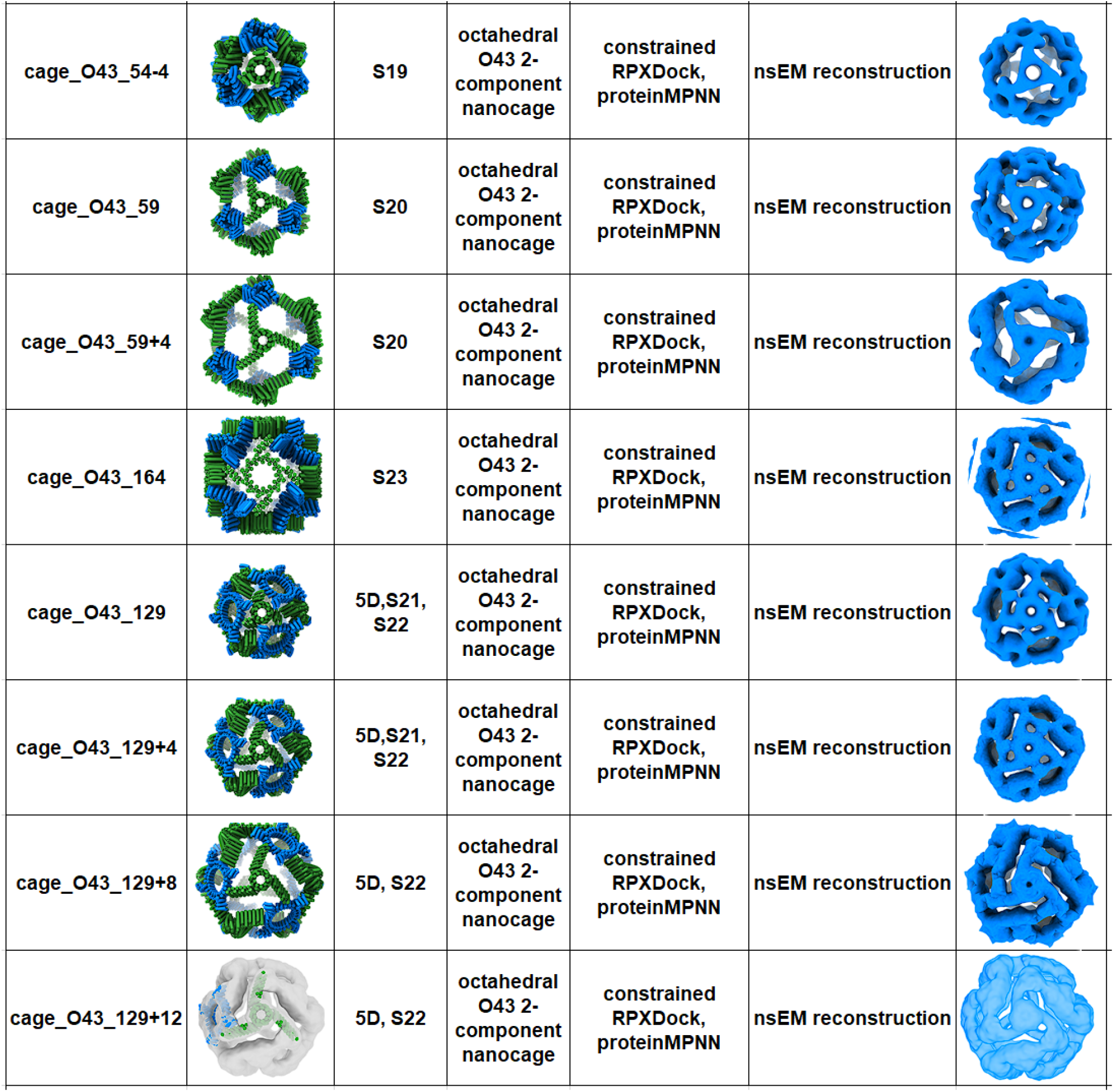
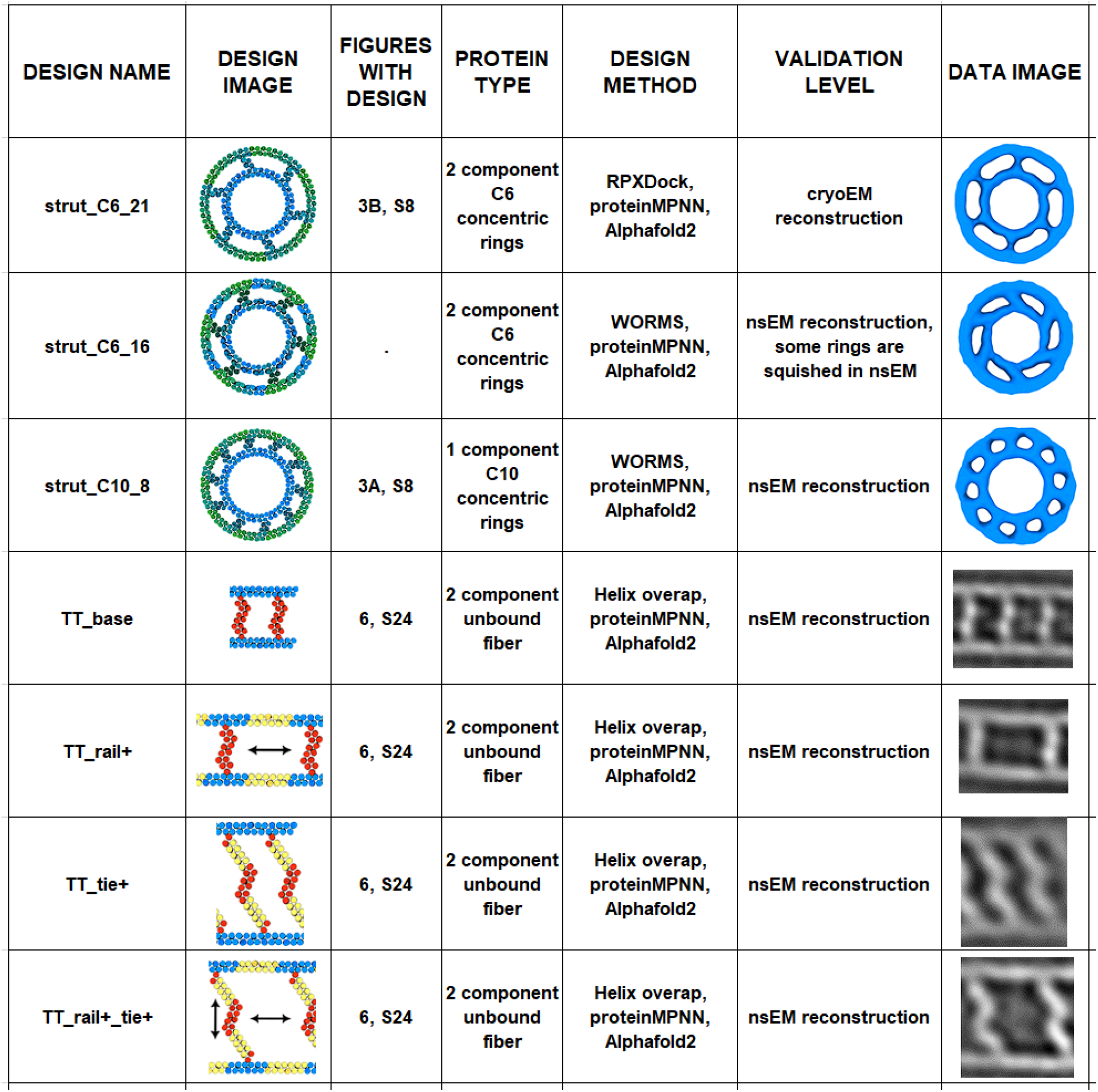
Summary of characterized designs. This is a reference table for all designs that are included with this publication. (**A**) Summary data for linear THRs that had either crystal structure or were solved as part of a cryo-EM structure. (**B**) Summary data for linear THRs that gave reasonable SAXS profiles. (**C**) Summary data for THRs that were validated as parts of other protein assemblies without a fully solved structure. (**D**) List of THRs that achieved high quality AlphaFold2 structure predictions but haven’t been validated yet. (**E**) Summary data for rings made from curved THRs. (**F**) Summary data for polygon oligomers made from angle-encoded THRs. (**G**) Summary data for cyclic oligomers made with other methods and THRs. (**H**) Summary data for THR arms added to a previously characterized design. (**I**) Handshake C2 designs that are pulled out of working nanocage designs. (**J**) Summary data for nanocages made with THR building blocks. (**K**) Summary data for strutted THR designs.

**fig. S2.**
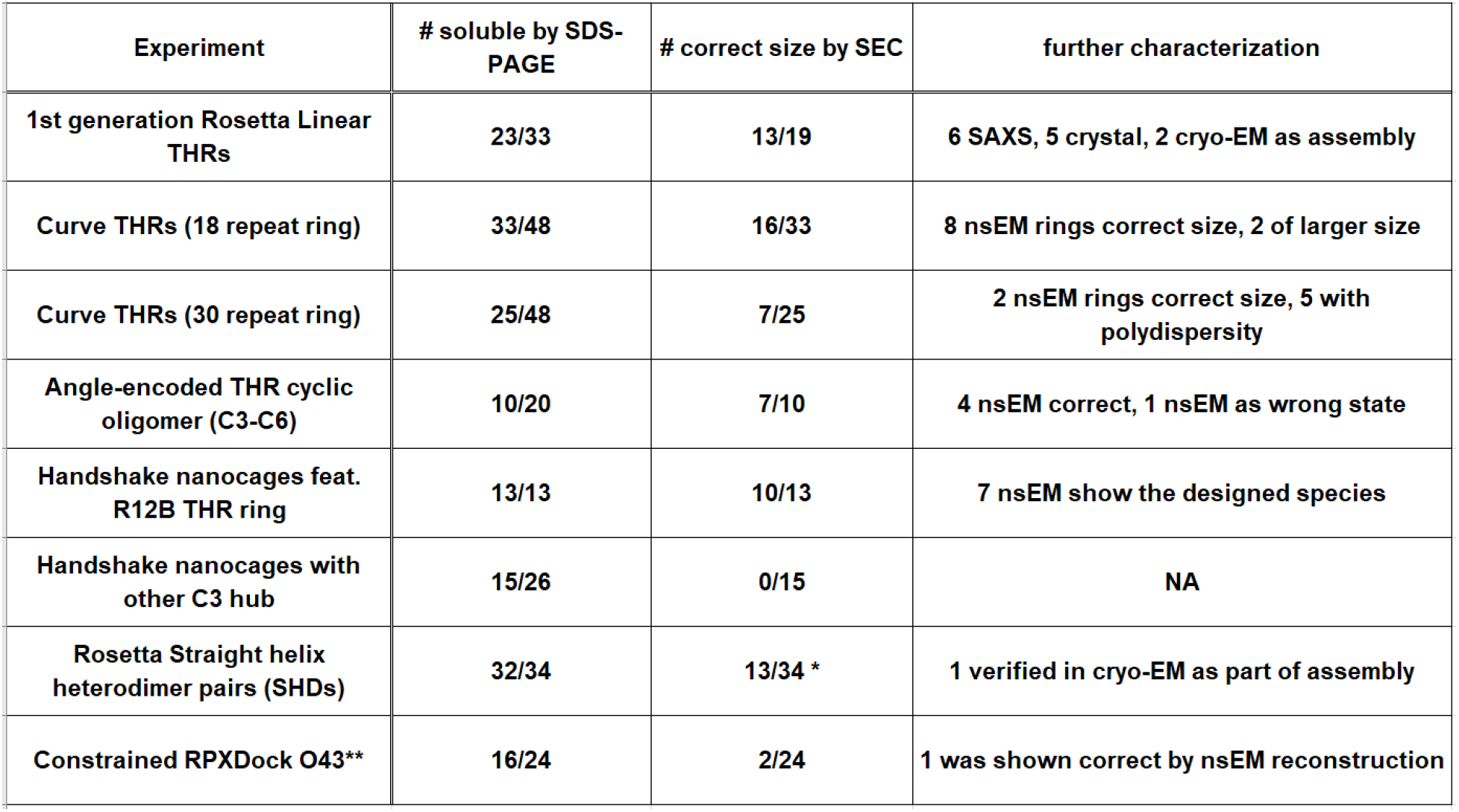
Summary of experimental outcomes. Summary of representative final rounds of design strategies for different architectures tested experimentally, showing a general sense of “success rate”. Number of designs yielding soluble protein via SDS-PAGE represent proteins post immobilized metal affinity chromatography (IMAC). Soluble designs were then passed through size exclusion chromatography (SEC). Designs that show elution peaks in the right size by SEC were then subject to appropriate biochemical characterization such as electron microscopy. (*:characterization was instead performed by Native-PAGE where appearance of a novel band upon combination of components suggested their interaction, **: this is from the final round of testing this architecture; there were additional successes from previous rounds)

**fig. S3.**
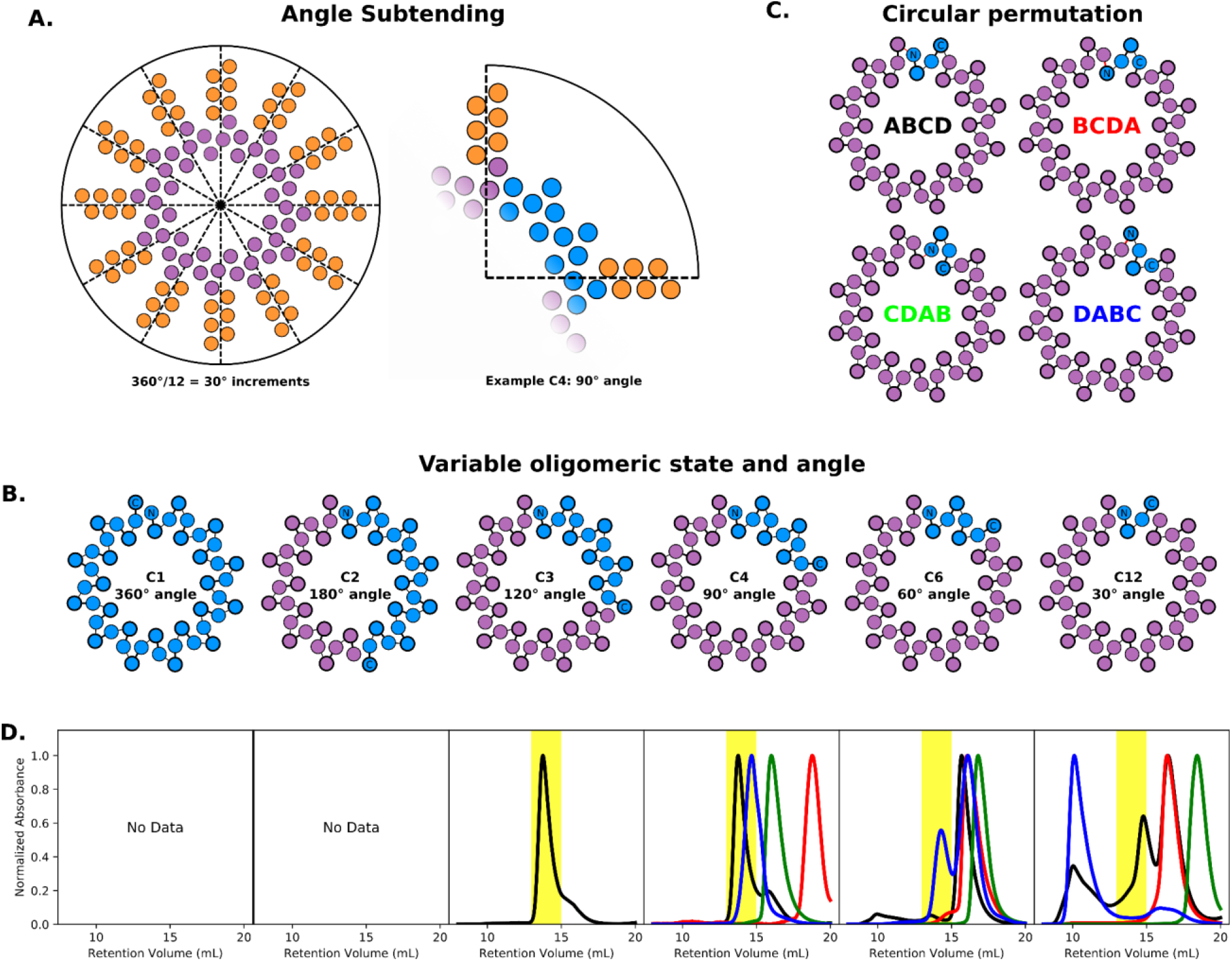
Modular properties of THR ring R12A. (A) An internally C12 curved THR ring structure can be subdivided in 30° increments; for example, a C4 variant of the ring would yield a 90° angle. (B) Depending on desired angle, the protein can be spliced into a variety of cyclic oligomers. (C) The structure is also planned in a way that circular permutations are simple, thus depending where fusion positions are needed, all eight (N- and C-) helical termini are available. (D) Preliminary experimental validation (size exclusion chromatography) confirms that simple splicing can produce proteins of the correct oligomeric state (yellow highlight), although some splices and circular permutations (color of the curve matches the color of the text from C) yield unassembled building blocks. C1 and C2 variants were not experimentally tested.

**fig. S4.**
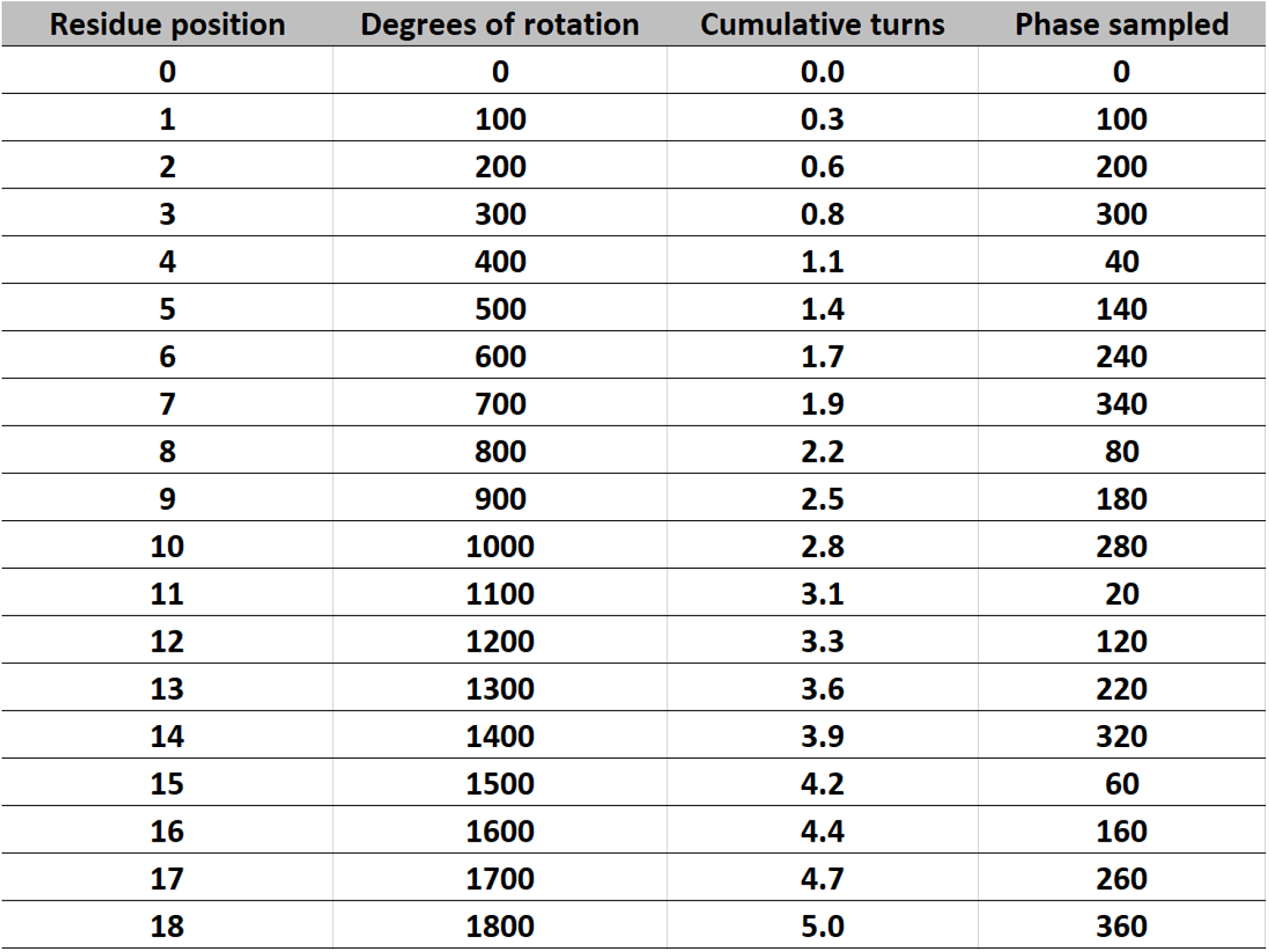
Helix phase repetition after 18 residues. The “alpha_helix_100” helices turn 100 degrees with every increment in sequence/residue position. After 1800 degrees, there have been exactly 5 complete turns (5 multiples of 360°). In the “Phase sampled” column, we see that the “Degrees of rotation” actually samples all the 20° increments in 0-360° if the “Degrees of rotation” are considered for their net rotation in a 0-360° range (if rotation exceeds 360°, subtract integer multiples of 360° until the rotation is in the 0-360° range).

**fig. S5.**
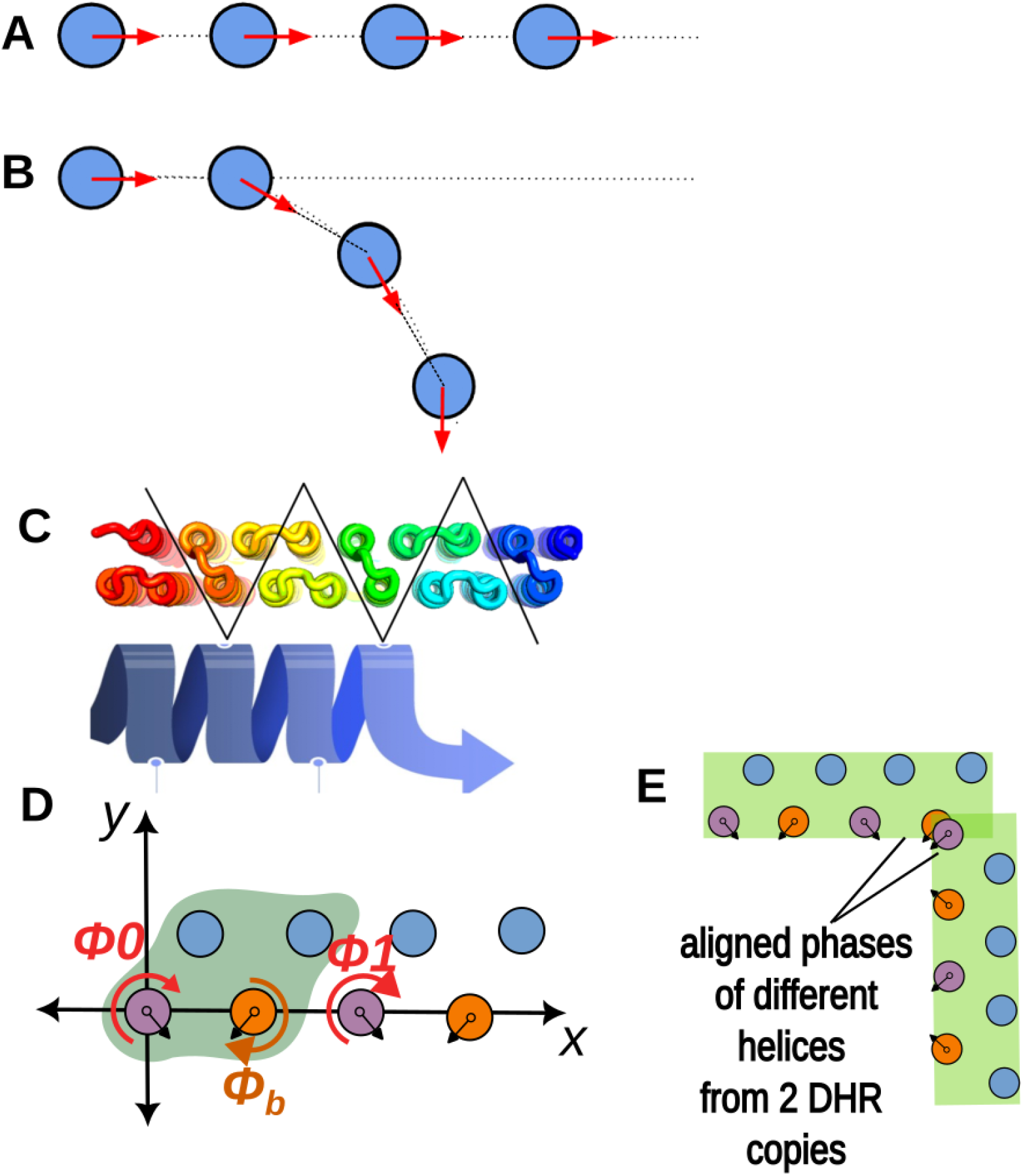
Additional diagrams of THR geometric properties. (A) Here we see that a THR repeat trajectory can be linear if there is no change in helix phase between the first and second copies of the “a” helix. (B) Here we see that a THR repeat trajectory will curve if there is a change in helix phase relative to the dashed line between the centers of the first and second copies of the “a” helix. (C) THRs with a repeat unit of 3 helices can be designed to be perfectly linear (such as THR5 and THR6) if they follow this flipping scheme when the helices are placed. (D) In a 4 helix repeat unit, the orange helix can be lined up with the trajectory-setting helices, but its phase can be freely sampled; this means if a purple helix were fused onto an orange helix, the fusion could make any assigned angle between linear THRs, as illustrated in (D).

**fig. S6.**
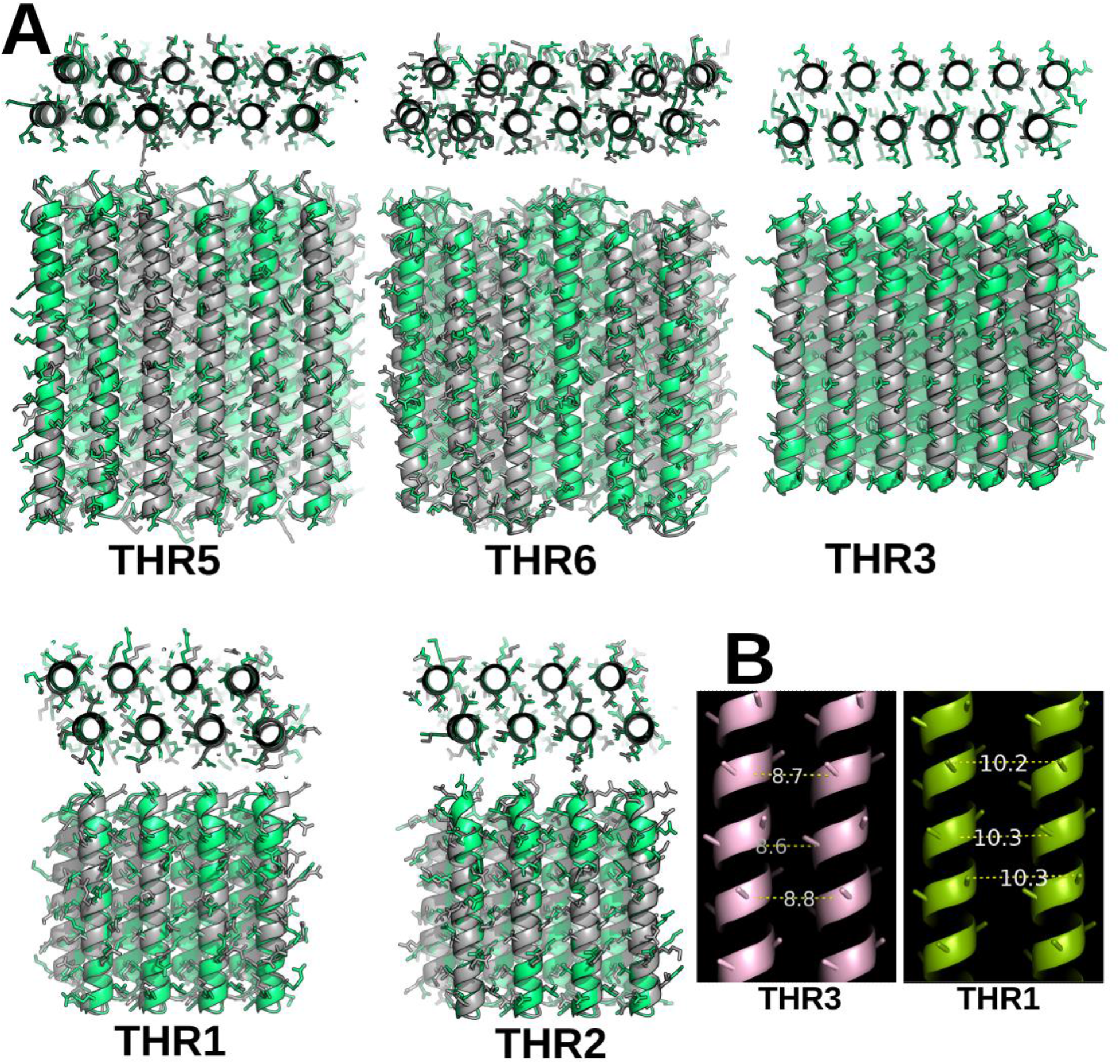
Linear THR crystal structures with side chains shown. (A) Grey crystal structures of linear THRs overlaid with design models in green, with side chains shown on both. THR3 only had enough resolution to show CA-CB atoms of the side chain sticks in the crystal structure model. (B) Measurements from PyMOL taken on representative innermost residue positions of THR crystal structures at matched positions in different repeat units (THR3 design spacing = 8.8 Å, THR1 design spacing = 10.0 Å)

**fig. S7.**
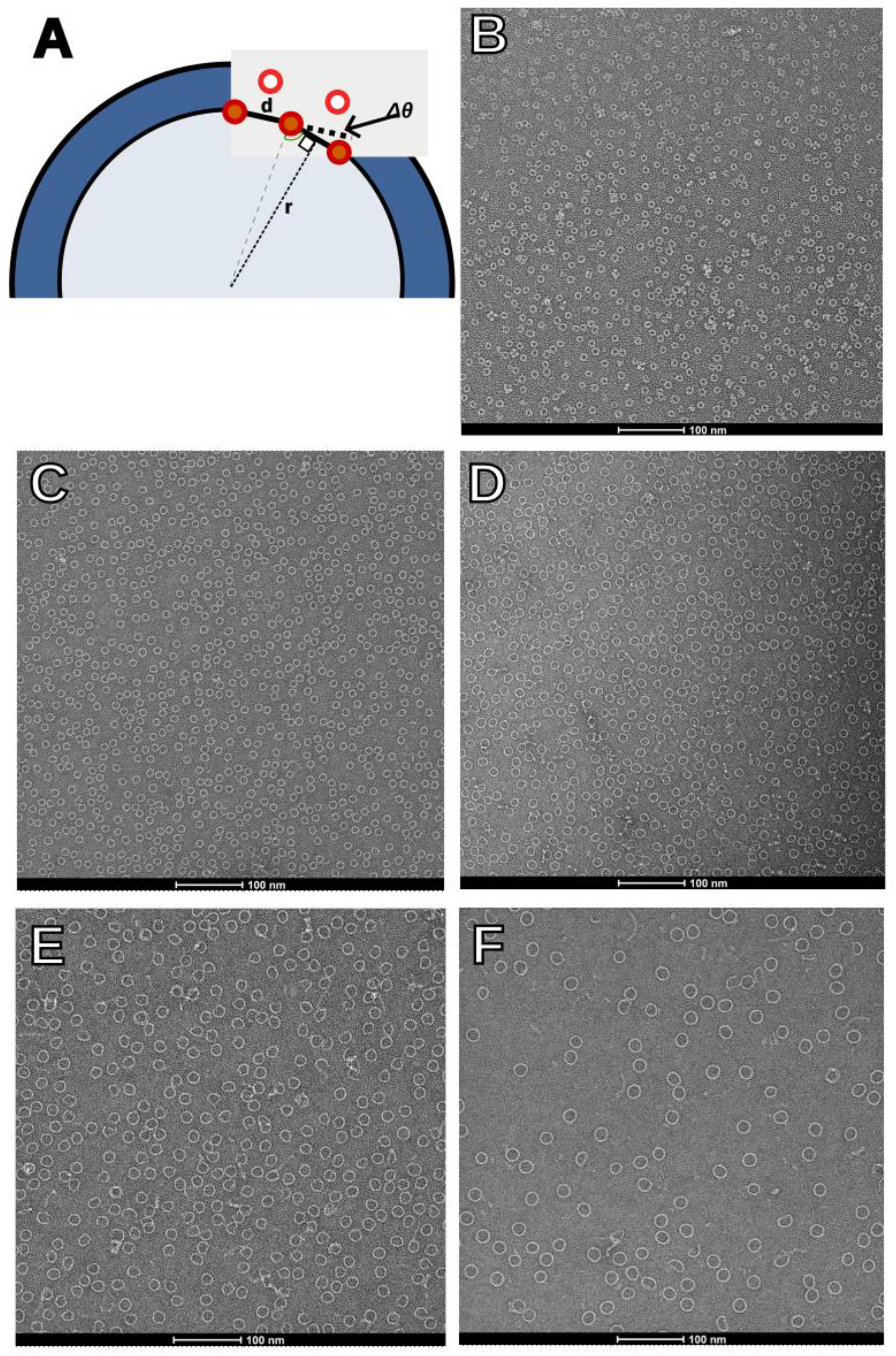
THR Ring diagram and raw nsEM micrographs. (A) This shows how a diameter (of a circle containing the centers of repeat-setting “a” helices) can be described as a function of the repeat parameters ***d*** and ***Δθ***: *radius* = 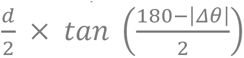; this value is smaller than the outer diameter dimension. (B-F) Raw nsEM micrographs of ***Curve*** THR rings. (B) *R12A* ring with 12 repeats, tested as C4 with expected outer diameter of 9 nm. (C) *R18G* ring with 18 repeats, tested as C6 with expected outer diameter of 12 nm. (D) *R20A* ring with 20 repeats, tested as C4 with expected outer diameter of 14 nm. (E) *R30E* ring with 30 repeats, tested as C6 with expected outer diameter of 21 nm. (F) *R30A* ring with 30 repeats, tested as C6 with expected outer diameter of 22 nm.

**fig. S8.**
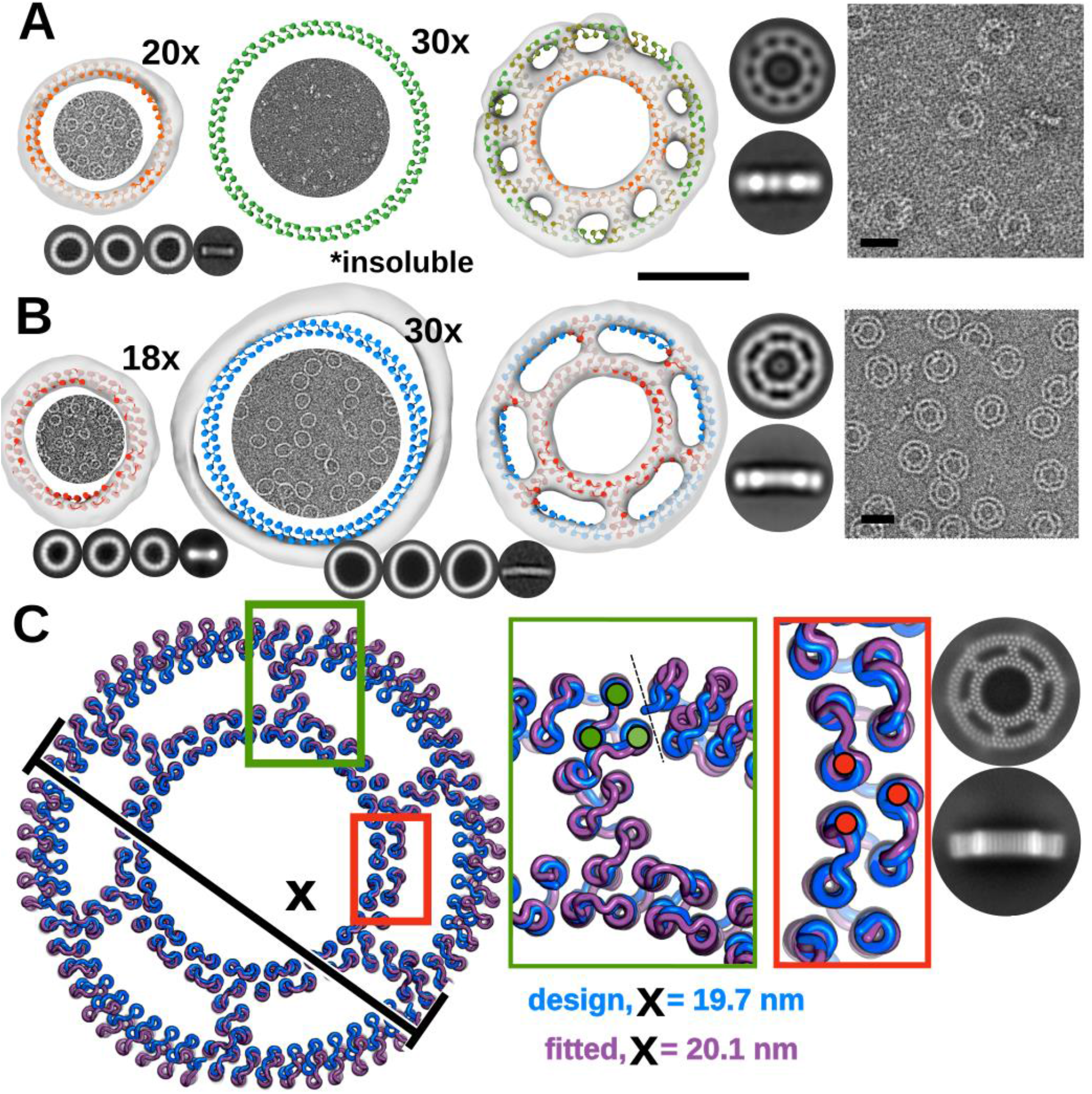
Effects of reinforcement on strutted rings. (A-B) Single ring (first two columns) and double ring (3rd column) design models are shown in cylindrical helix representation and overlaid with C1 nsEM reconstructions. The deviations from perfect circular shape are much larger for the single rings than the reinforced double ring structures. nsEM micrograph samples of the single rings are shown inside of the design models and of the double rings, at the far right. Black scale bars = 10 nm, 2D class averages are not to scale. (A) (Left) A 20-repeat ring *R20A* (orange), tested as a C4 was monodisperse and close to the intended size. (Middle) A 30-repeat ring (green) tested as a C6 expressed insolubly. (Right) A strutted combination of these rings as a single component C10 design *strut_C10_8* rescues the outer ring, as illustrated in the reconstruction, the 2D class averages, and in the raw micrograph. (B) (Left) An 18-repeat ring *R18A* (red), tested as a C6, was monodisperse and close to the intended size. (Middle) a 30-repeat ring *R30B* (blue) tested as an intended C6, was seen more often as a heptamer than a hexamer and was often oblong. (Right) A strutted combination of these rings as a two-component C6 design *strut_C6_21* with the struts attached to the inner ring. (C) Design model of the two-component C6 design *strut_C6_21* from B (blue) with cryo-EM fitted model (purple) shows good alignment in the inner ring (red inset) while deviation is observed in the interface (dashed line) between strut and outer ring (green inset). Structures in inset boxes are locally aligned to 3 helices, indicated by filled circles on top of the helices.

**fig. S9.**
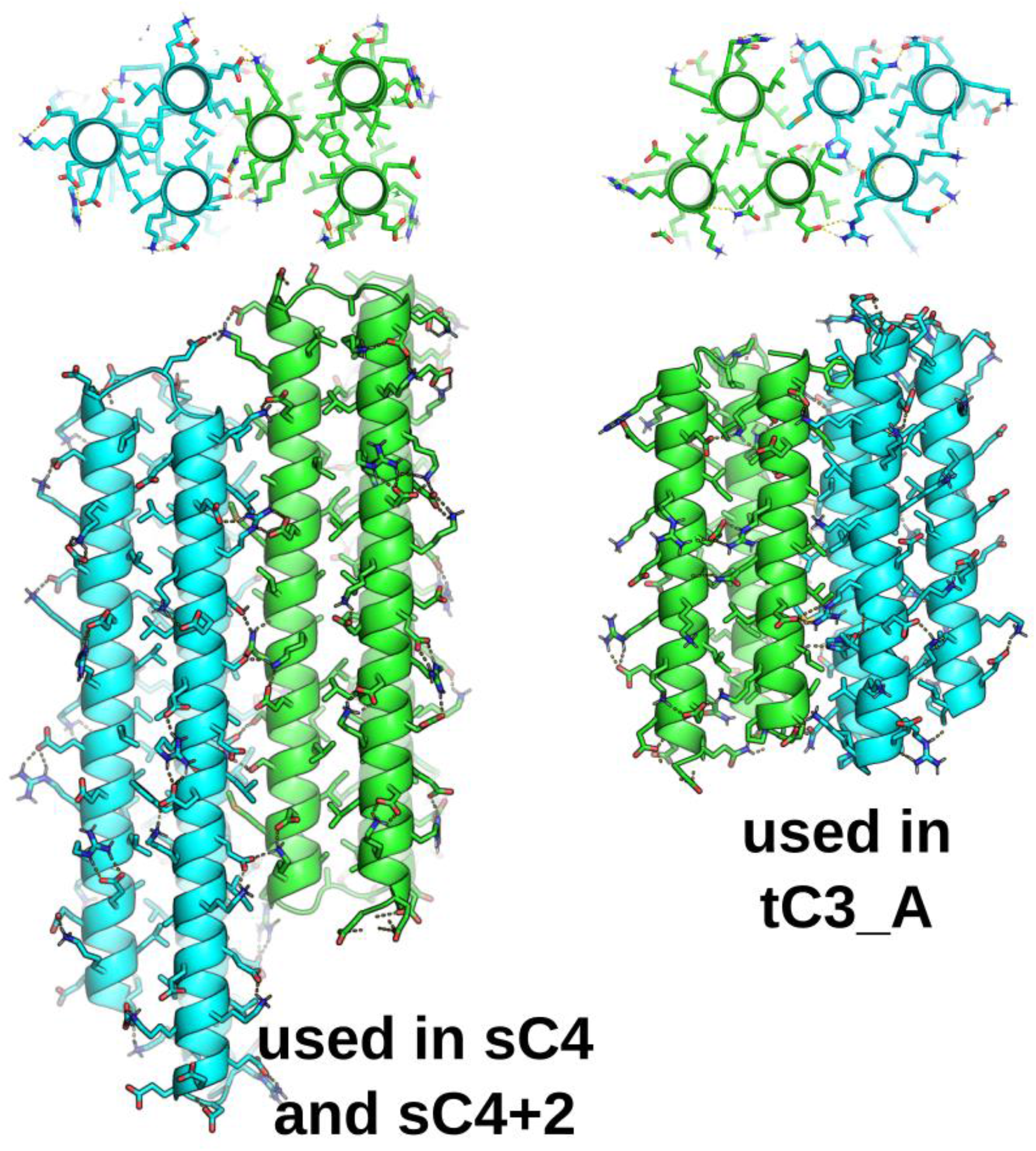
Images of SHD models. Examples of design models of straight hetero-dimers (SHDs) utilizing buried hydrogen bond networks to confer interaction specificity.

**fig. S10.**
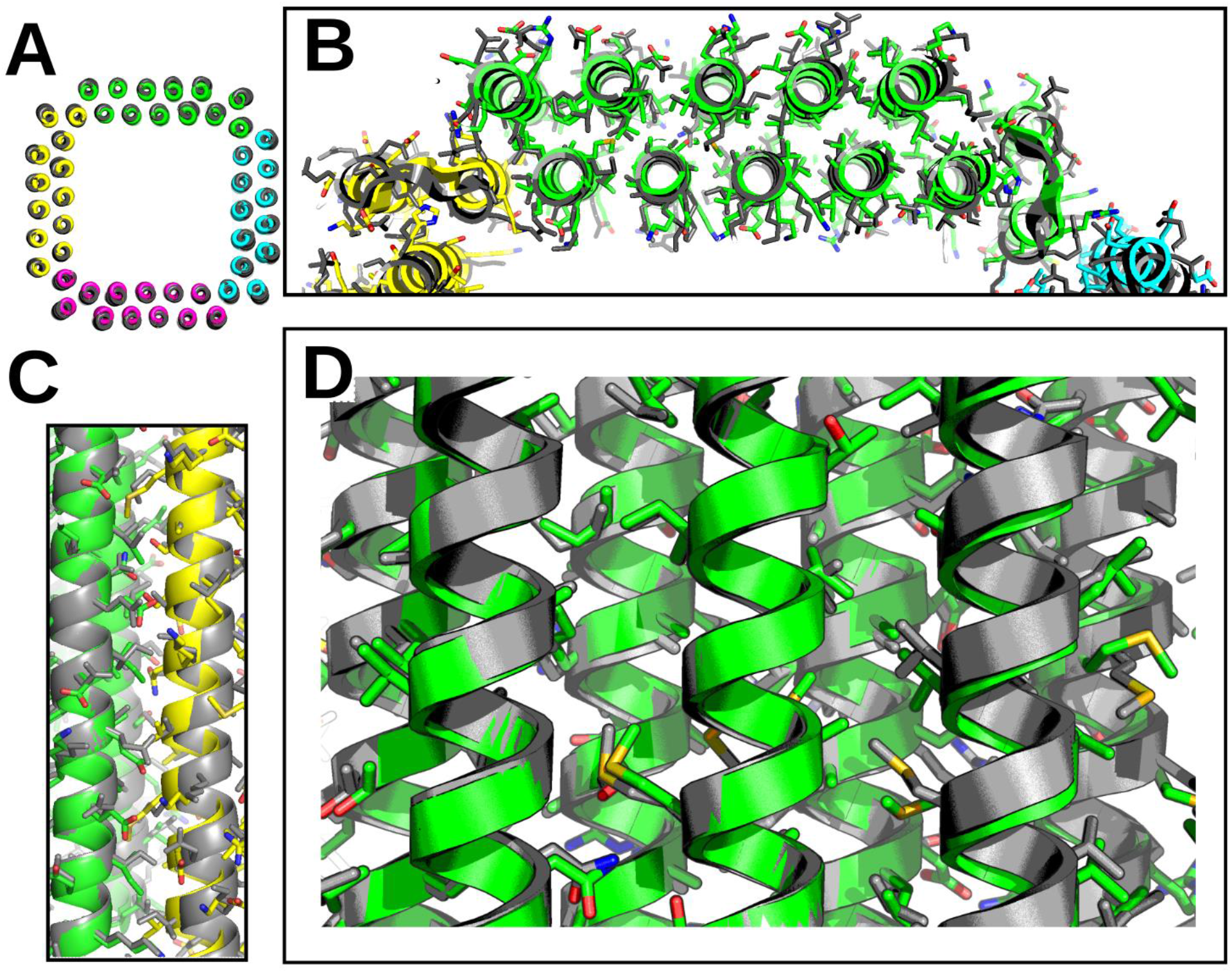
Cryo-EM experimental model of sC4 compared to design. Design model (colored) overlaid with cryo-EM structure (gray). (A) View from down the z-axis (1.6 Å backbone RMSD). Zoom in on the top (B) and side (C) views showing overall agreement of helices. (D) Zoomed in view showing overall agreement of hydrophobic side chain residues.

**fig. S11.**
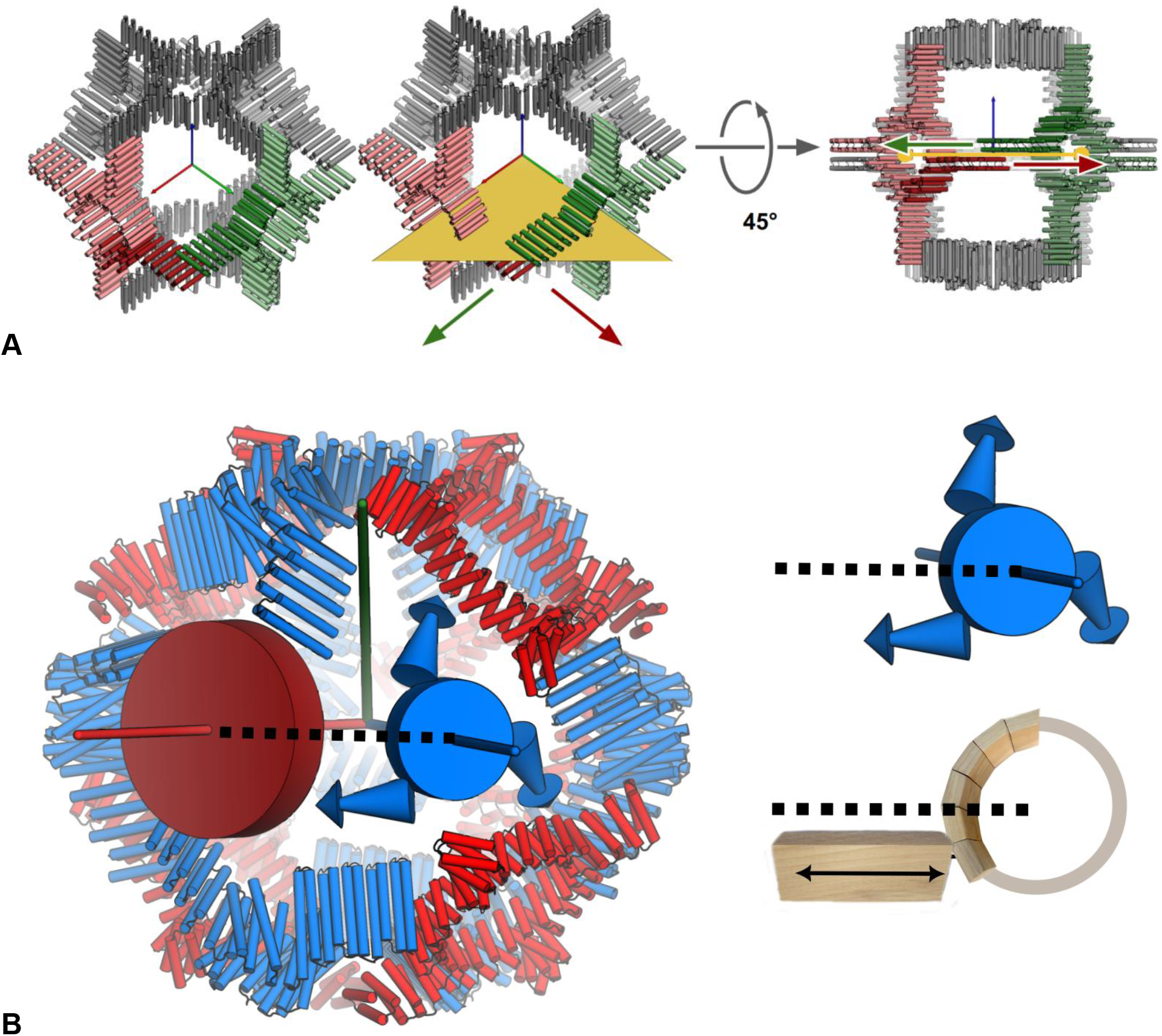
Geometric constraints for expandable polyhedral symmetrical assemblies. (A) An O4 octahedral nanocage showing the 4-fold symmetry axes aligned to the coordinate axes x, y, and z (red, green, and blue, respectively). The THR propagation vector (large red and green arrows) must be parallel to the plane (in yellow) formed by the two encompassing symmetrical axes, x and y. Deviations off this plane will cause contractions and extensions of the THR to clash and/or not connect to its symmetrical partners using the same contacts. (B) Similarly for a two-component architecture (O43 in this case), the component with the THR propagation vector (3D blue arrow) must remain parallel to the plane created by the 3-fold axis (blue) and 4-fold axis (red).

**fig. S12.**
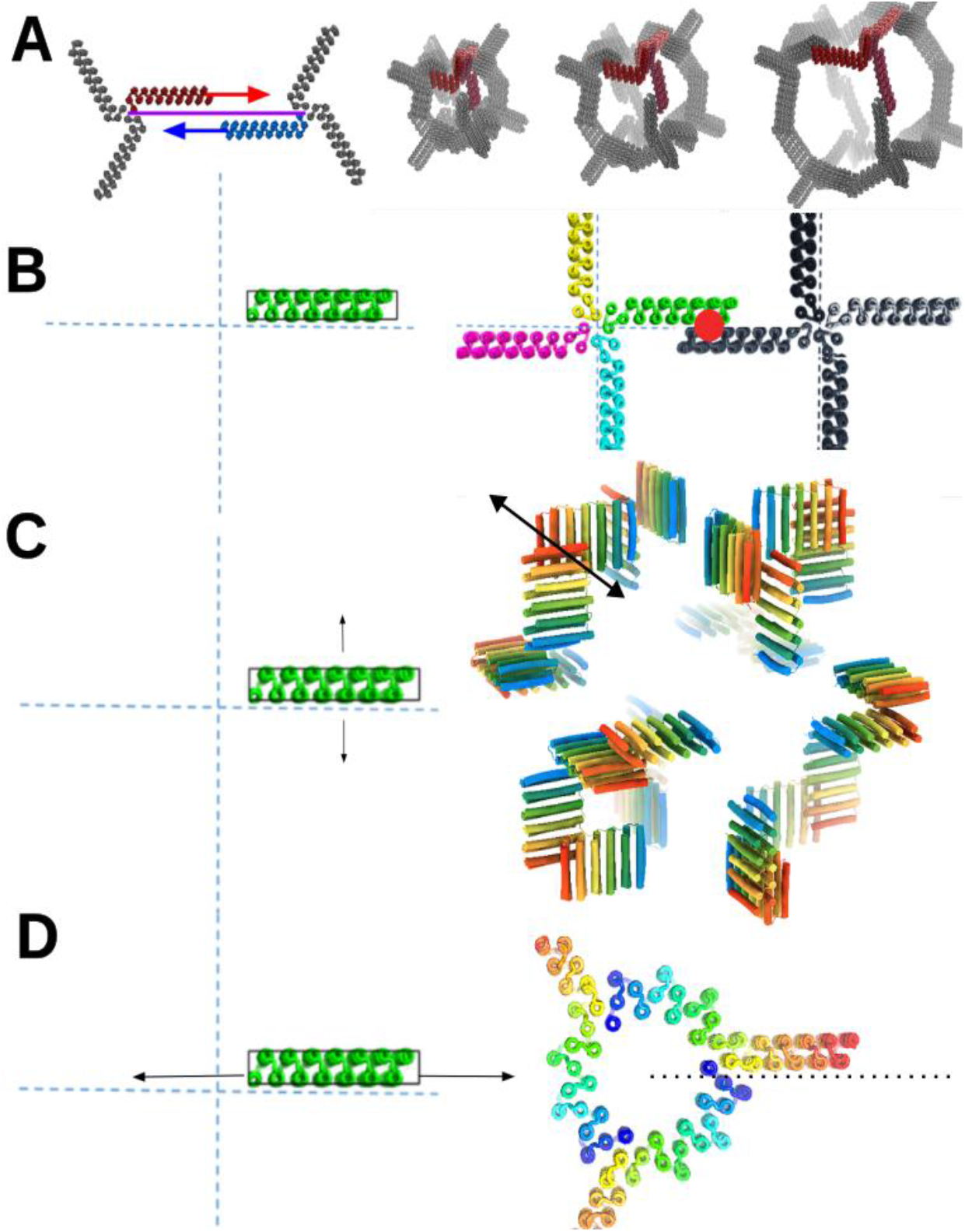
Design strategy for THR Handshake nanocages. **(A)** To maintain the expandability of the cages, the THR propagation vector (red and blue) must be maintained parallel to the plane generated by the symmetry axes (purple). **(B)** the interface formed between the two THRs is the “handshake” interface (red dot). **(C)** Handshake C2 interfaces with angles corresponding to the desired architecture are designed by sampling the THR in the “y-direction” (sideways, but parallel to the required plane, and still within handshake contact range if C2 were applied), then treating that positioned THR as a cyclic oligomer for running RPXDock in nanocage symmetries, such as the O4 output shown to the right. The sampling allows Z movement of the THR, which manifests as sliding (arrow) along the handshake interface to find the best rigid body position to design. **(D)** The same THR is sampled in the “x-direction” (forwards and backwards) to find good fusion positions to the cyclic oligomer using RPXDock Axle protocol.

**fig. S13.**
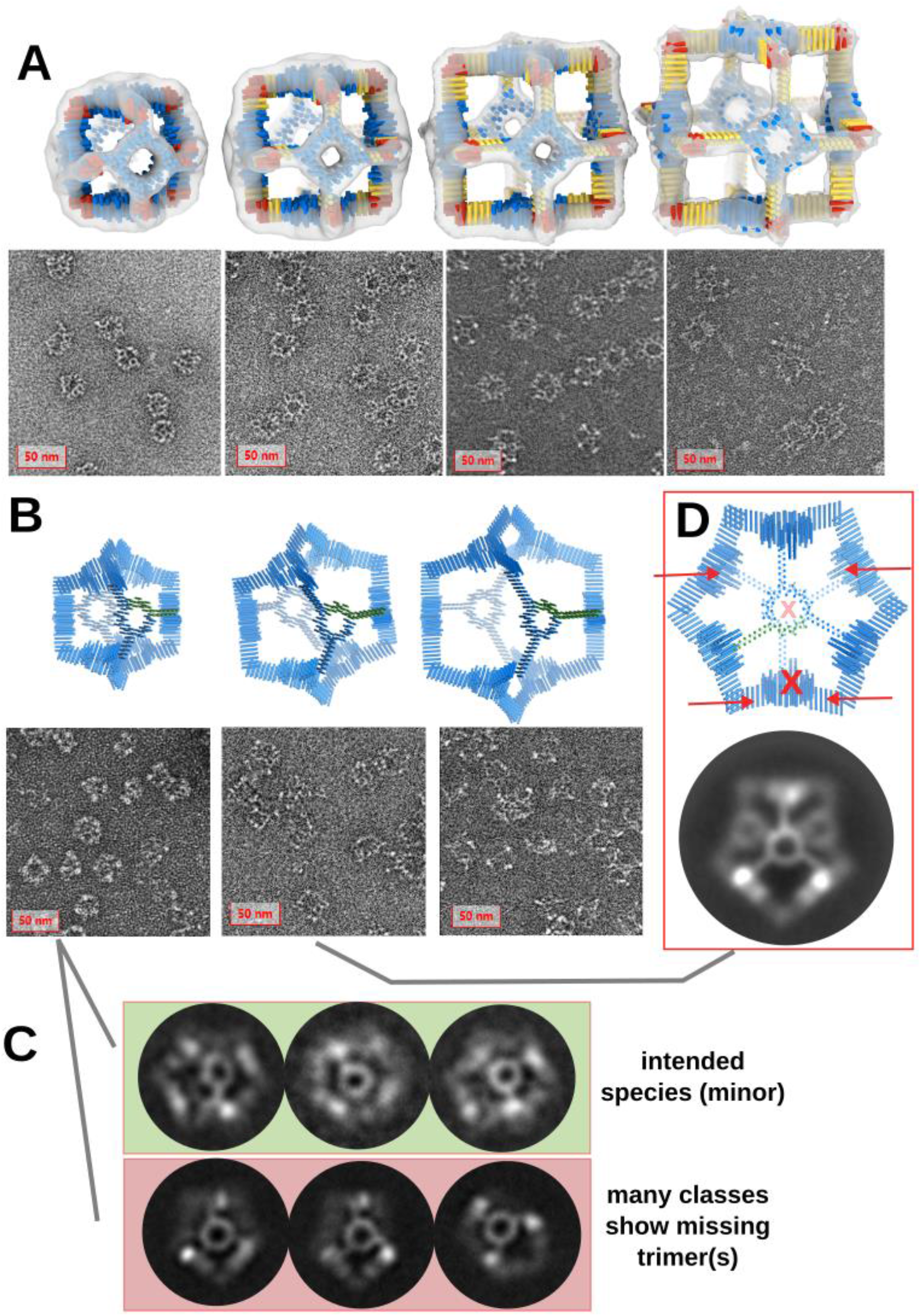
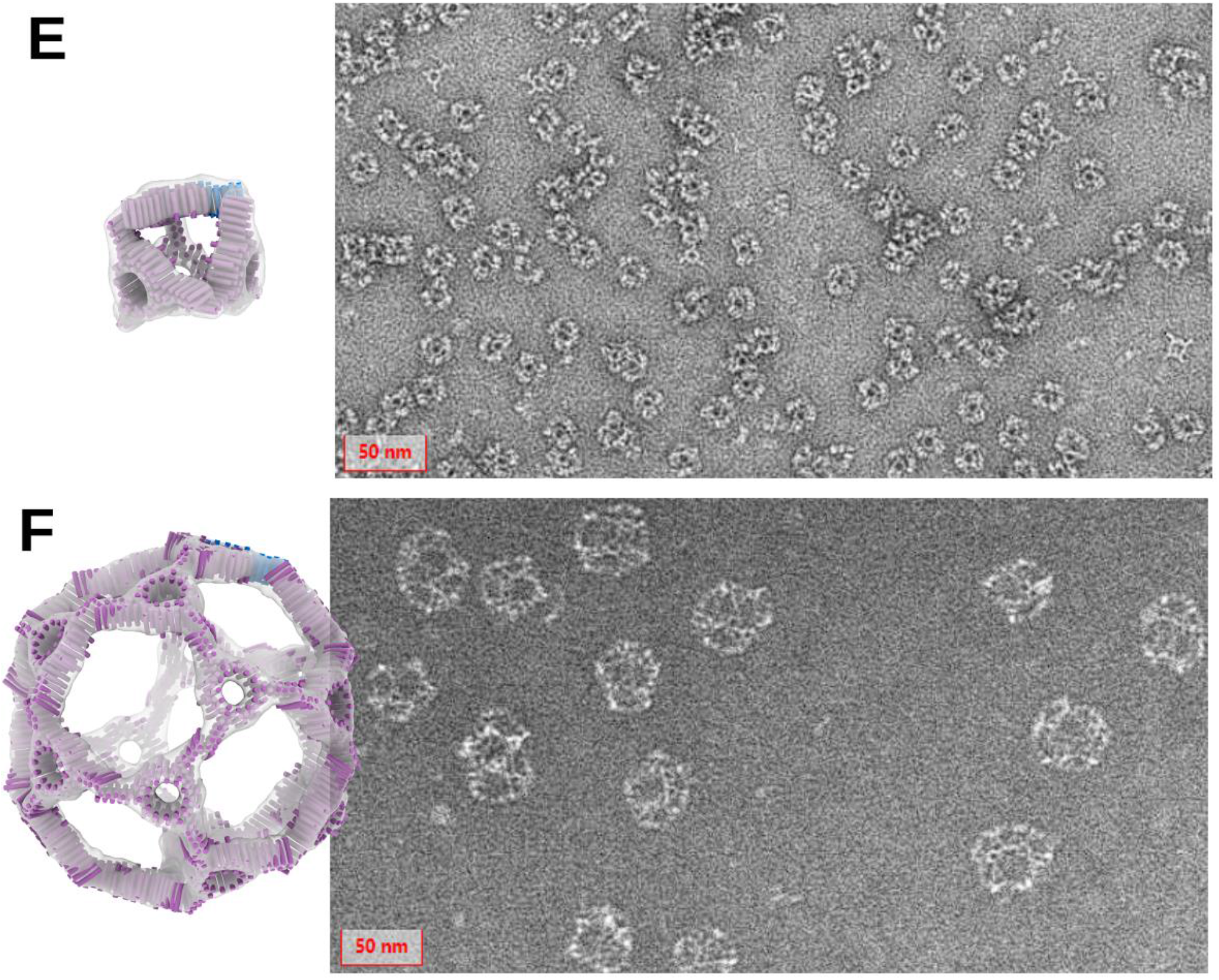
Additional nsEM data for THR Handshake nanocages. **(A)** Representative negative stain EM micrographs of each of the *cage_O4_34* sizes; +0, +4, +8, and +12 respectively. **(B)** Representative negative stain EM micrographs of the *cage_O3_2*0 sizes; +0, +4, and +8. **(C)** Class averaging of the base size shows populations of both the intended octahedral species and off-target populations with missing trimers. **(D)** Class averaging of the two larger sizes shows a monodisperse off-target population where two of the trimeric units are missing from the octahedral structure. **(E)** Representative negative stain EM micrograph of cage_T3_101 showing clear particles. **(F)** Representative negative stain EM micrograph of cage_I3_8 show clear particles, but many particles seem to be missing trimeric subunits.

**fig. S14.**
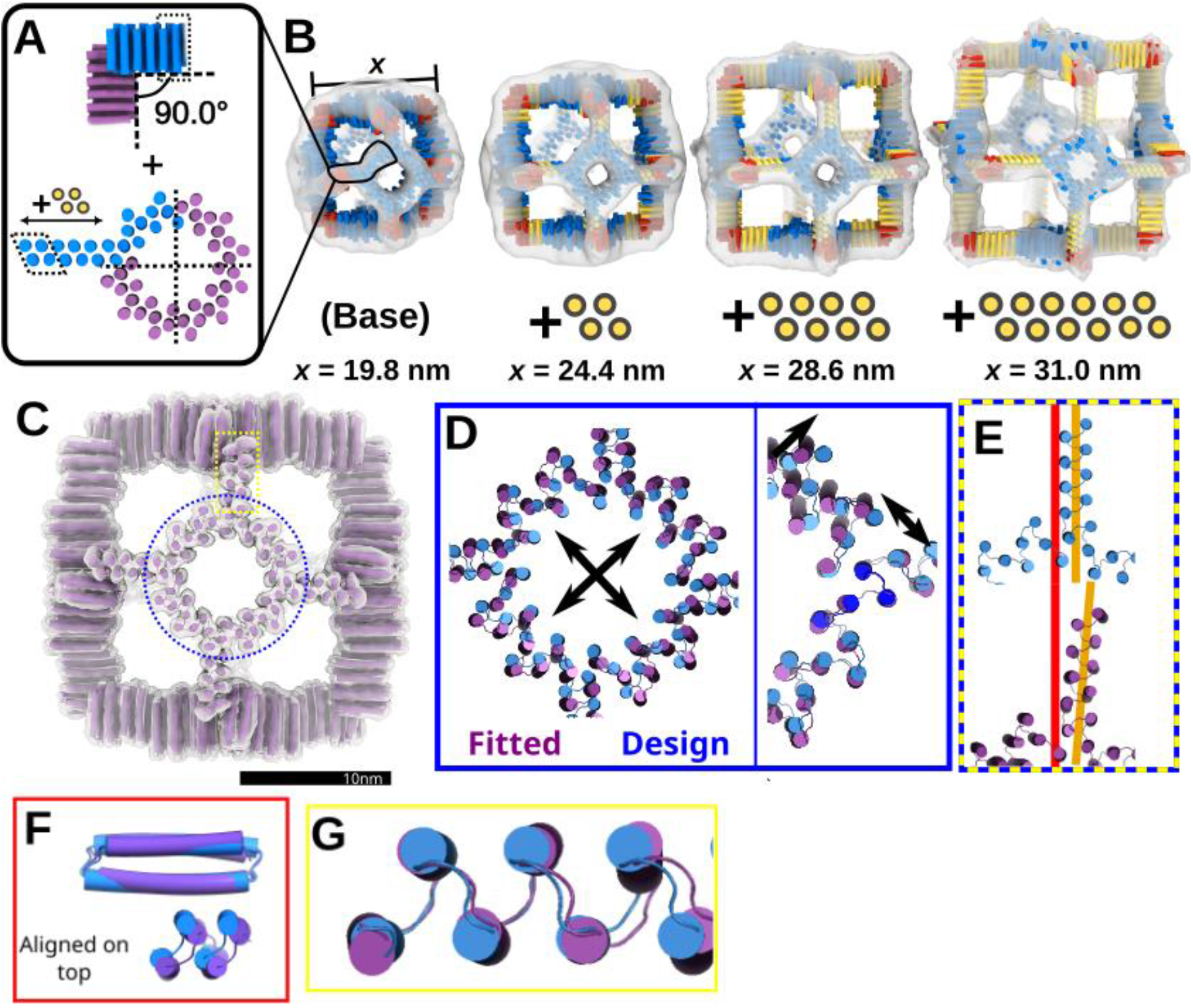
Cryo-EM analysis of a modular O4 nanocage. **(A)** Previously described (Fig. 4) construction method of combining two modules to make the nanocage. **(B)** The O4 octahedral handshake nanocage was characterized at 4 sizes with nsEM reconstruction map overlaid with design models. Yellow represents regions that were extended by locally repeating the linear THR structure/sequence. A distance “x” is measured across the 3D reconstructions of each size. **(C)** Comparison of (left) the full O4 design model in blue and (right) the fitted model in purple in the cryo-EM map. Dashed regions indicate areas of focus in panels D-G. **(D)** Overlay of Handshake models aligned on one Hand to exaggerate the shift observed in the other Hand. **(E)** Overlay of Arm models aligned on the top 4 helices. **(F)** (left) Overlay of the Ring models to show general expansion of the ring. (middle) zoom-in on the Ring repeat element of the design model (Blue) fused to the Arm and making an interface with another Ring repeat on a different chain. Models are aligned on the blue repeat. (right) Overlay of the three Ring repeats in the fit (shades of purple) with the structural Ring repeat of the design model (blue). **(G)** Visualization of the Arm repeat axis (orange) for the Design and Fit models aligned along the C2 symmetry axis (red).

**fig. S15.**
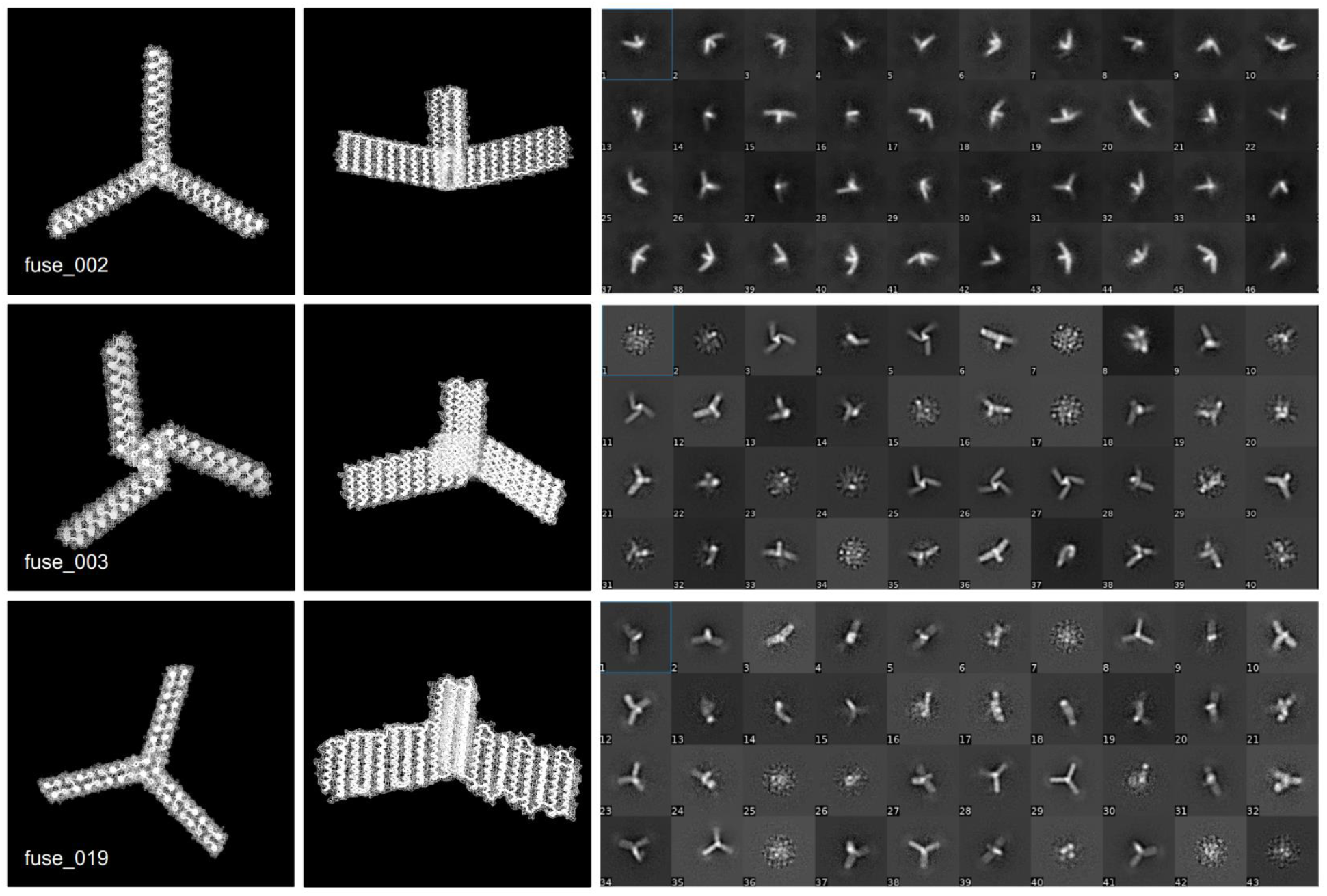
Examples of HelixFuse *de novo* helical bundles to THR proteins. (left) Design models showing 3-fold axis and side view. (right) Raw 2D class averages from negative stain electron microscopy showing many classes resembling the design model

**fig. S16.**
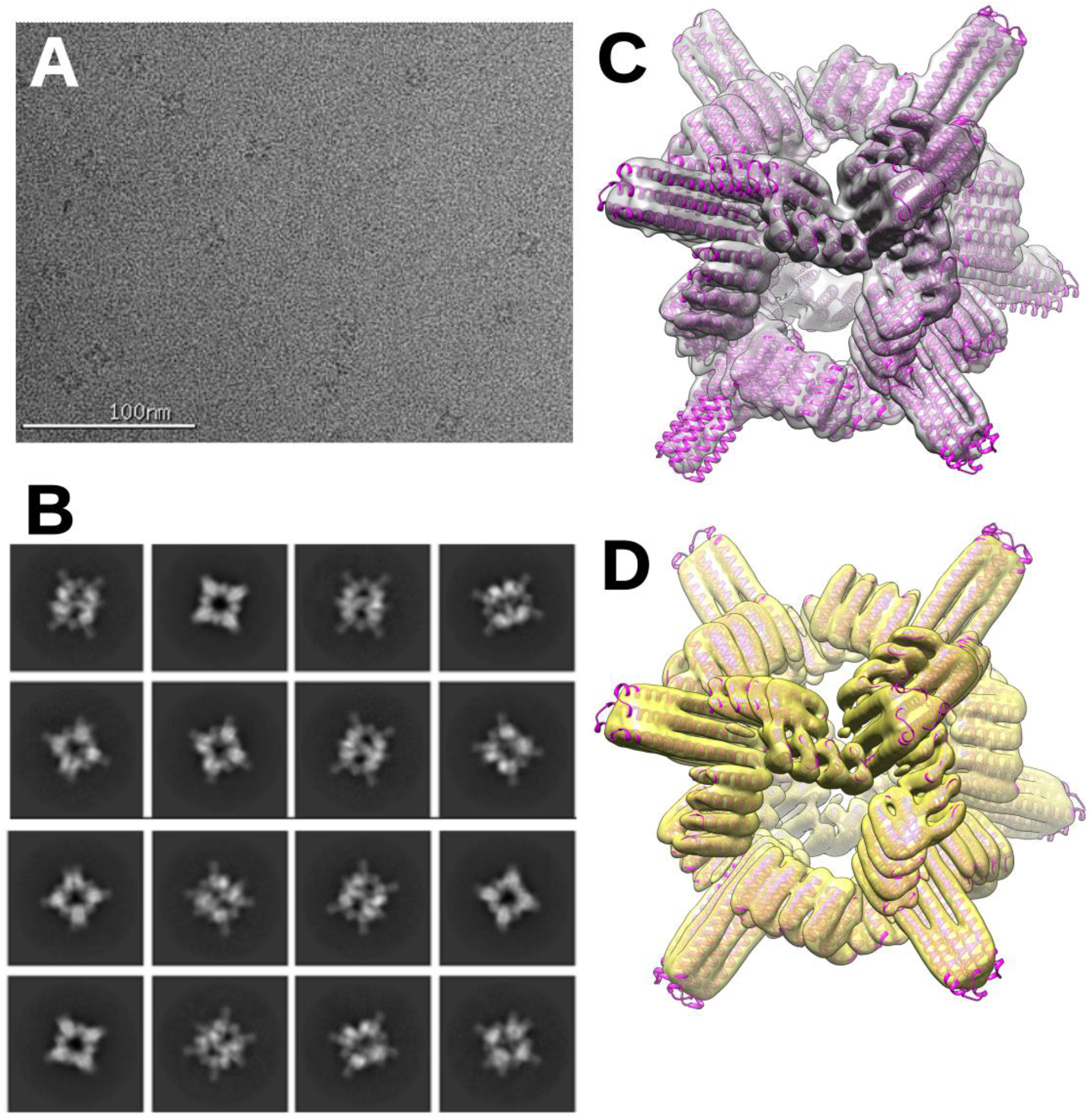
Cryo-EM of *cage_O3_10*. **(A)** Representative Cryo EM micrograph of *cage_O3_10*. (**B**) Representative 2D class averages. **(C)** Cryo-EM map (gray) without symmetry imposed (C1) with the design model (magenta) fit in as a rigid body **(D)** Cryo-EM map (yellow) with O symmetry with the design model (magenta) fit in as a rigid body.

**fig. S17.**
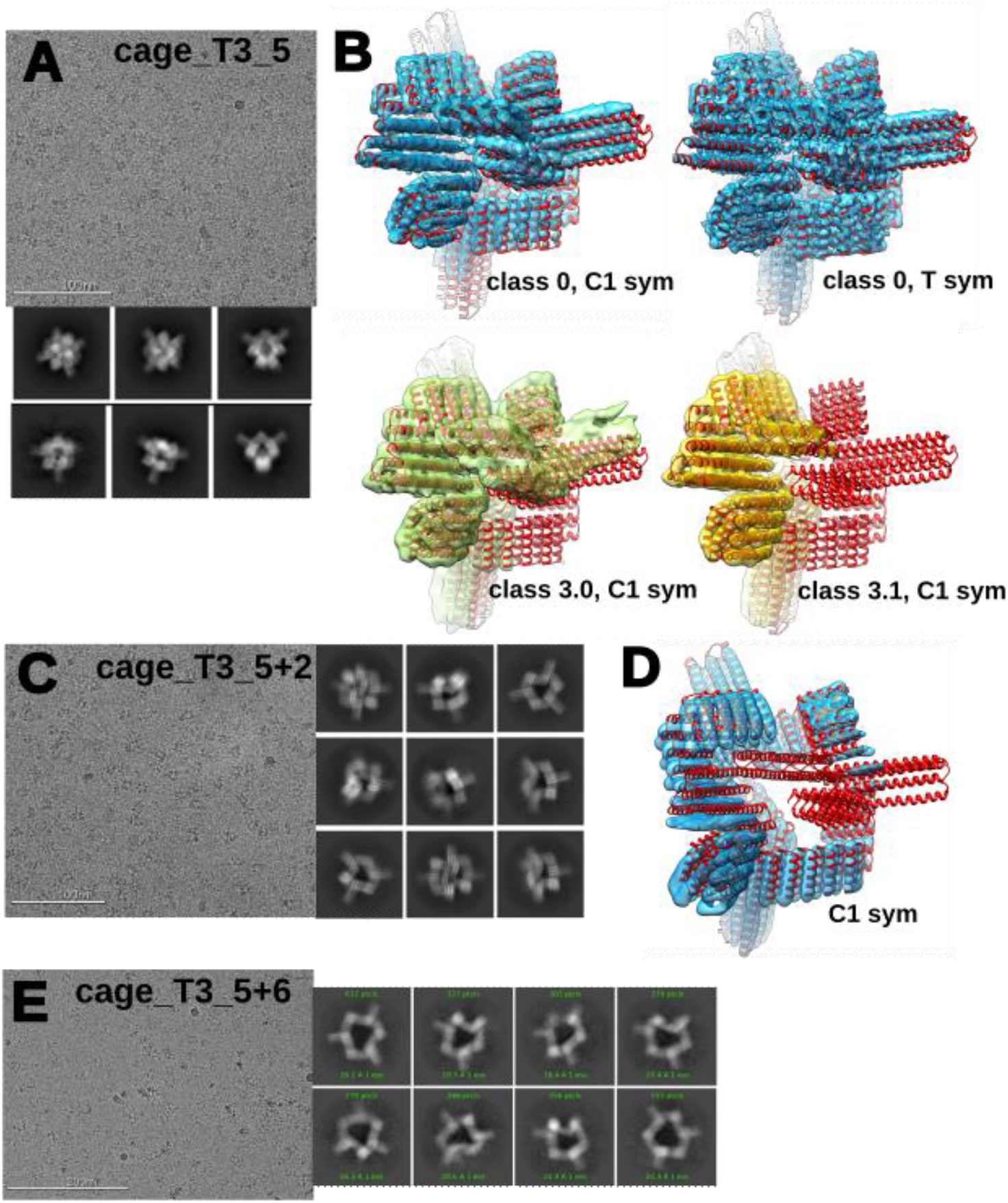
Cryo-EM of cage_T3_5 and its expansions. **(A)** Representative cryo-EM micrograph of *cage_T3_5*, and representative 2D class averages (scale bar = 100 nm). **(B)** Cryo-Em maps of *cage_T3_5* from different heterogeneous refinements as described in Figure S25, with design model (red) fit in as a rigid body. **(C)** Representative cryo-EM micrograph of expanded *cage_T3_5+2*, and representative 2D class averages (scale bar = 100 nm). (**D**) Cryo-EM map of *cage_T3_5+2* without symmetry applied (C1), with the design model fit in as a rigid body. Density corresponding to one trimer is absent, suggesting that this trimer is not present in the cage at this size. **(E)** Representative cryo-EM micrograph of expanded *cage_T3_5_+6*, and representative 2D class averages (scale bar = 200 nm).

**fig. S18.**
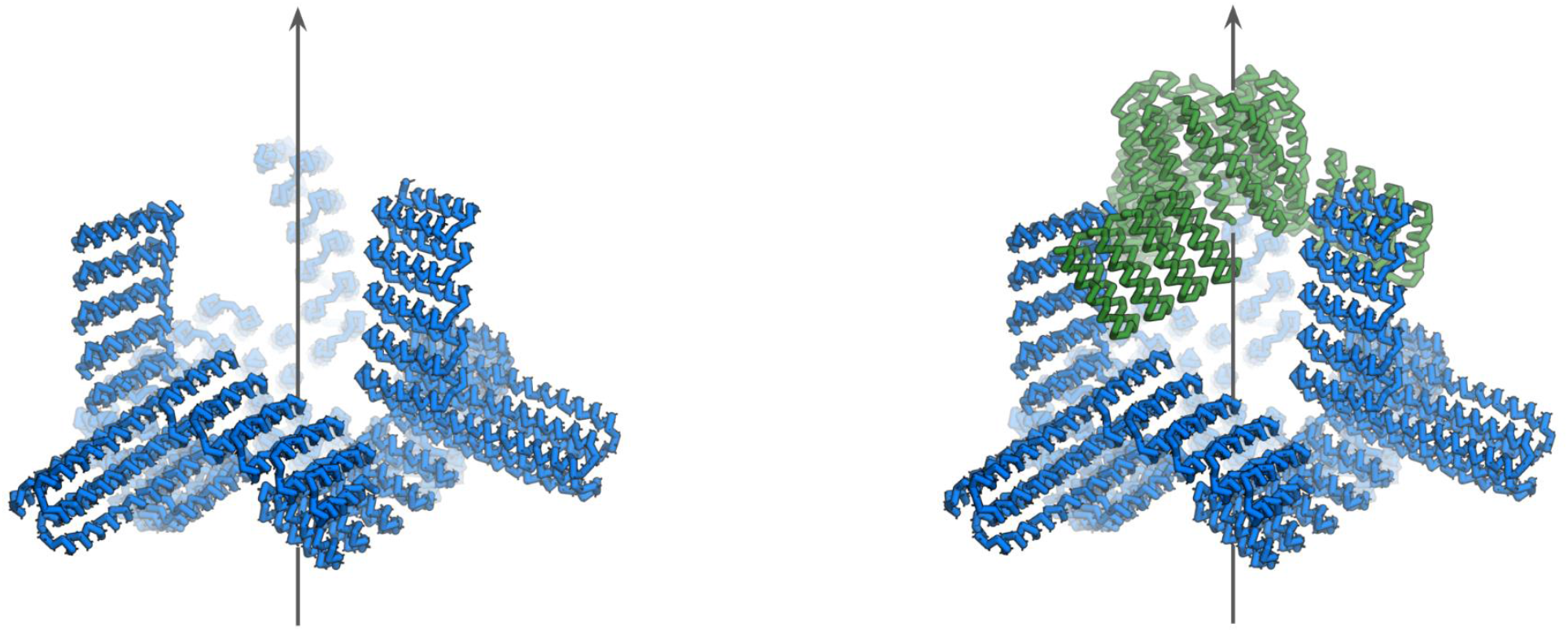
Cage polarization scheme using cage_T3_5_+2. Conceptual illustration to show that if a model of an incomplete cage were to be approximated from cryo-EM data into a simplified symmetry such as C3 (blue) aligned to the z-axis (left), it can be possible to design a new trimeric binding partner (green) that better satisfies the new opened configuration, resulting in breaking true tetrahedral symmetry into C3 cyclic symmetry.

**fig. S19.**
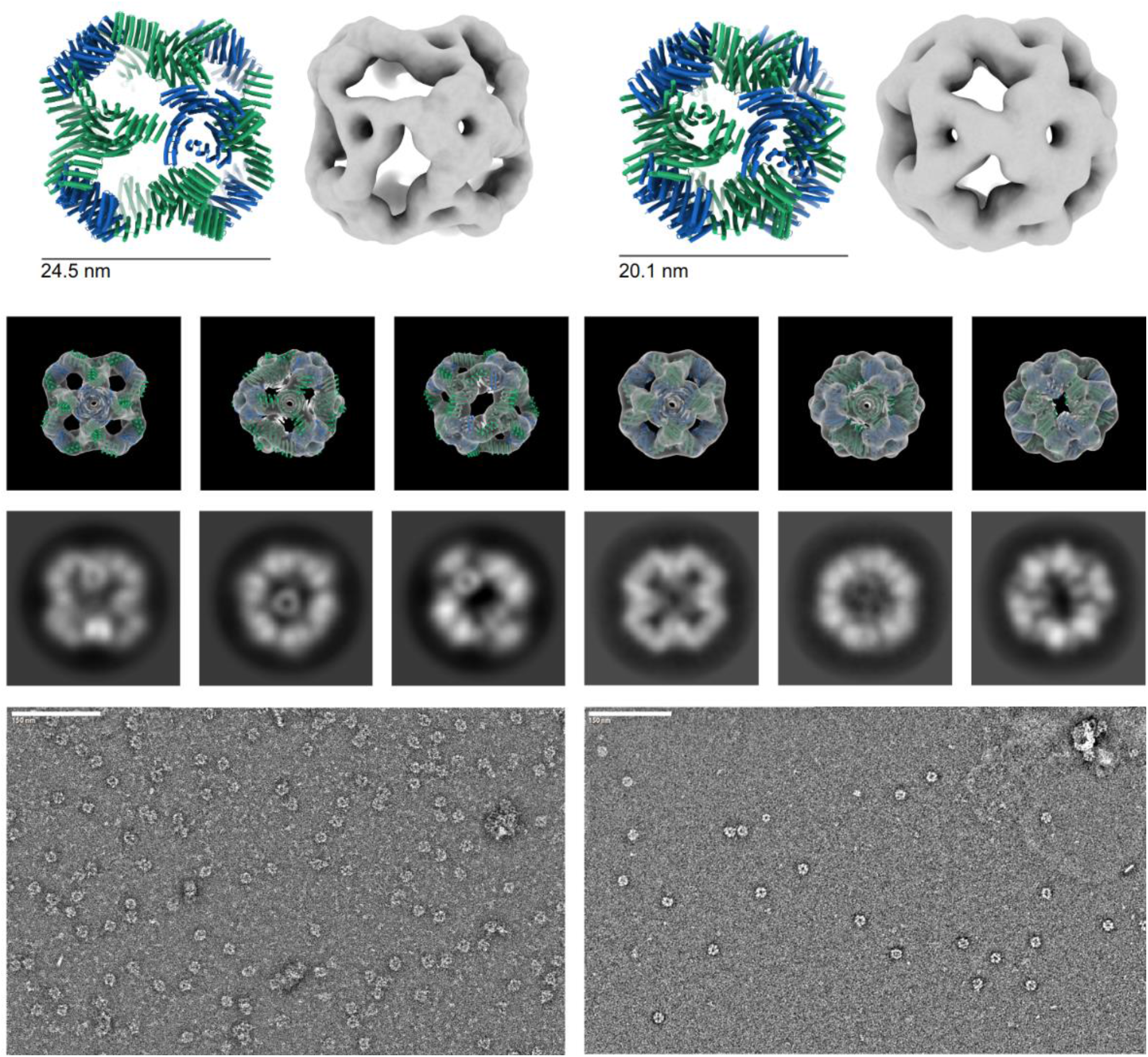
Negative stain EM of cage_O43_54 series. Negative stain electron microscopy of cage_O43_54 (left) and cage_O43_54_-4 (right). Design model with 3D reconstruction (top), design model with reconstruction map overlaid compared to selected class averages from each of the axes views (4-fold, 3-fold, and 2-fold) (middle), and a representative micrograph at 57,000x magnification (bottom); scale bar = 150 nm.

**fig. S20.**
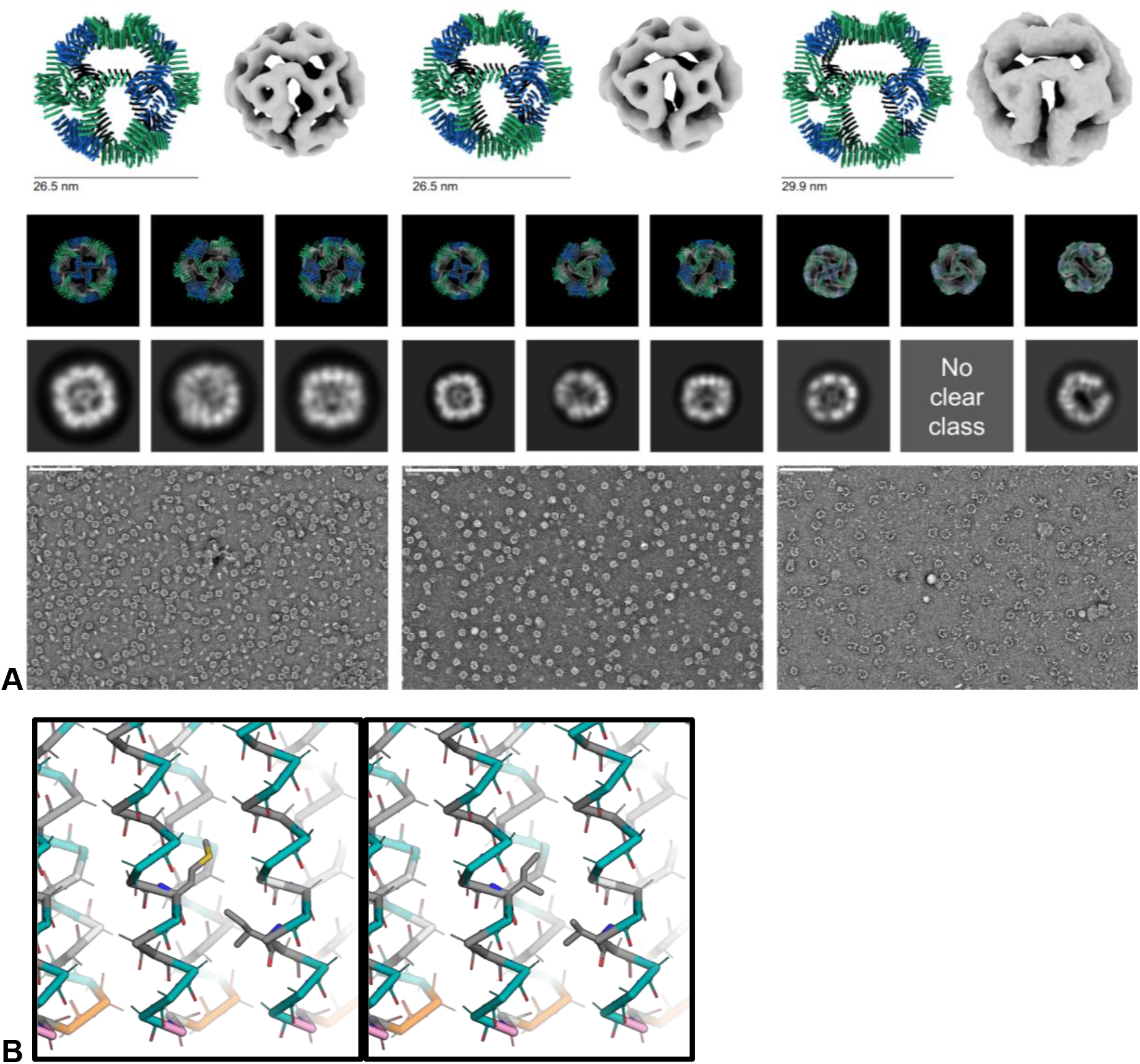
Negative stain EM of cage_O43_59 series. **(A)** Negative stain electron microscopy of cage_O43_59 (left), cage_O43_59_noMet (middle), and cage_O43_59_+4 (right). Design model with 3D reconstruction (top), design model with reconstruction map overlaid compared to selected class averages from each of the axes views (4-fold, 3-fold, and 2-fold) (middle), and a representative micrograph at 57,000x magnification (bottom); scale bar = 150 nm. The cage_O43_59_noMet construct is identical to cage_O43_59 except for methionine and related mutations. **(B)** Original backbone of THR4 (left) and noMet mutant (right). Backbones of polar residues shown in teal, proline in orange, glycine in pink, alanine in white, and hydrophobic residues in gray. Each methionine (one per repeat, first instance M34) can be mutated to isoleucine and the nearby isoleucine (first instance I78) mutated to valine to compensate without disrupting the structure.

**fig. S21.**
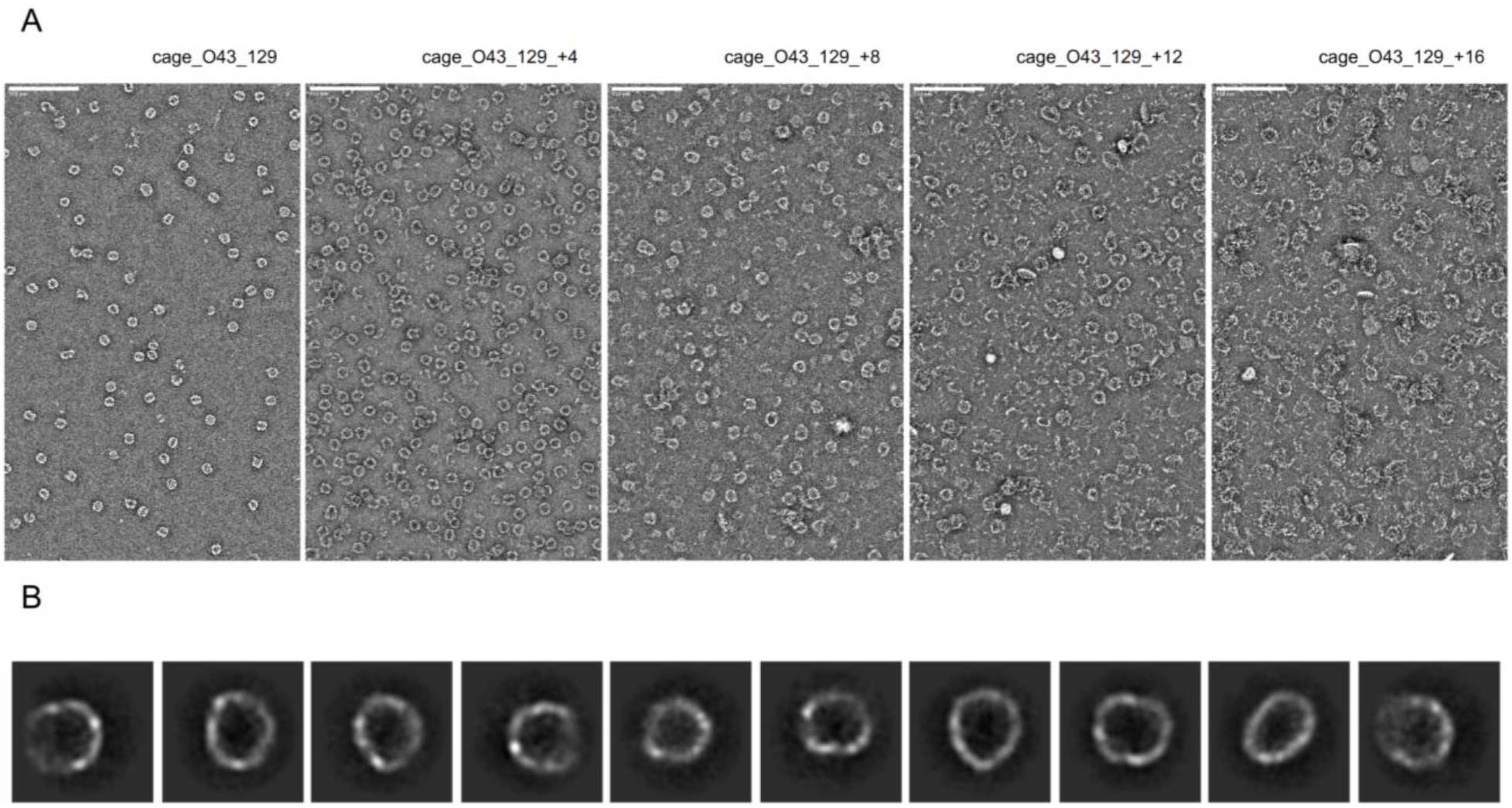
Negative stain EM of cage_O43_129 series. (A) Negative stain electron microscopy micrographs of cage_O43_129 (left) to cage_O43_129+16 (right). Increased amounts of unassembled and/or misassembled protein in the background increases with extended THR arm length. (B) Representative class averages of cage_O43_129+16. Although the design yielded distinct pickable particles, the resulting class averages were heterogeneous and often oblong, suggesting misassembled particles.

**fig. S22.**
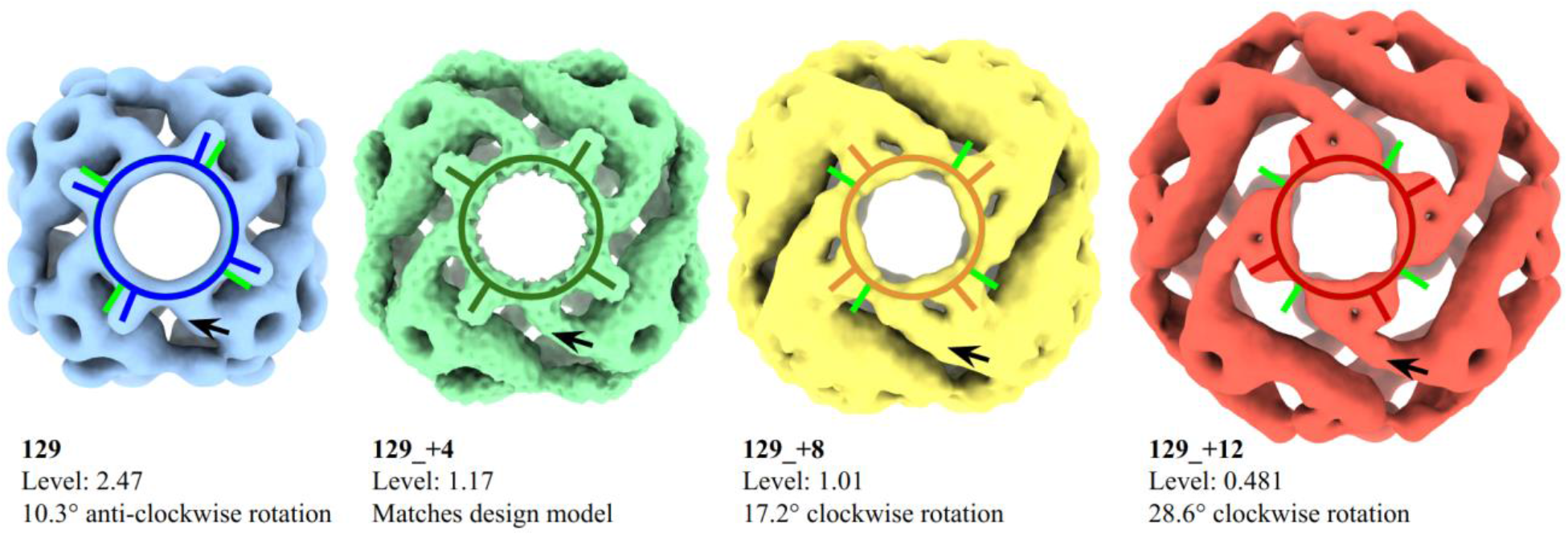
Rotational deviation of cage_O43_129 series. Left to right: Negative stain EM 3D reconstruction maps of cage_O43_129 (blue), cage_O43_129_+4 (green), cage_O43_129_+8 (yellow), and cage_O43_129_+12 (red) visualized along the 4-fold symmetrical axis. The C4 component that matches the design model is shown as a dark green circle (cage_O43_129_+4). In cage_O43_129, the C4 component rotates 10.3° anti-clockwise (blue circle), while in cage_O43_129_+8 and cage_O43_129_+12 it rotates 17.2° and 28.6° clockwise respectively (yellow and red circles) around the 4-fold axis, relative to the designed position (bright green). A black arrow depicts the THR component that is extended in each assembly; the C3 component remains unperturbed rotationally as designed and interacts with the same interface area even though the interaction angle shifts.

**fig. S23.**
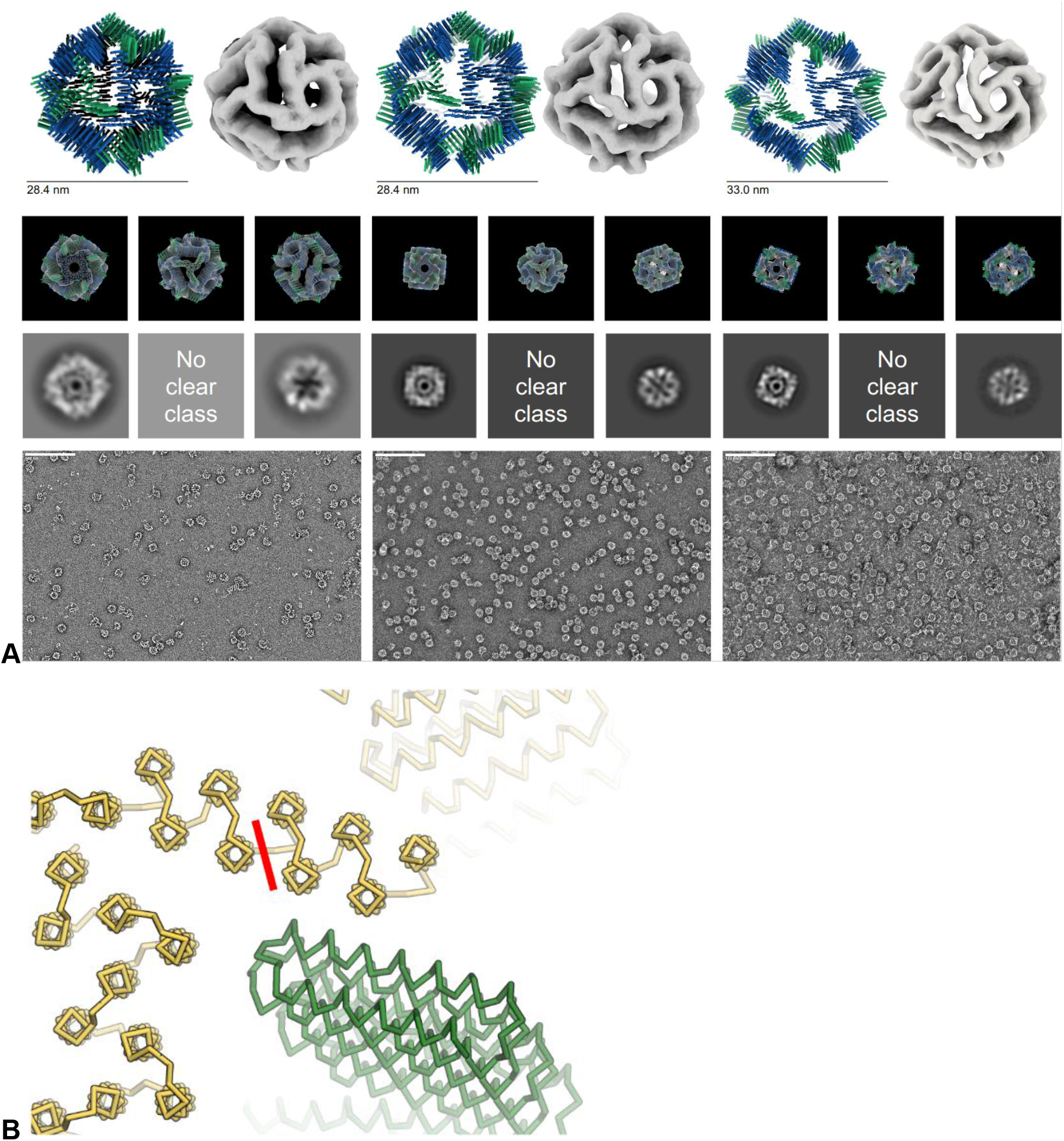
Negative stain EM of cage_O43_164 series. (A) Negative stain electron microscopy of cage_O43_164 (left), cage_O43_164_cut (middle), and cage_O43_164_+2_cut (right). Design model with 3D reconstruction (top), design model with reconstruction map overlaid compared to selected class averages from each of the axes views (4-fold, 3-fold, and 2-fold) (middle), and a representative micrograph at 57,000x magnification (bottom); scale bar = 150 nm. ￼(B) ”Cut” variant expresses the first five helices of the C4 subunit as a separate chain; cut location depicted with the red line.

**fig. S24.**
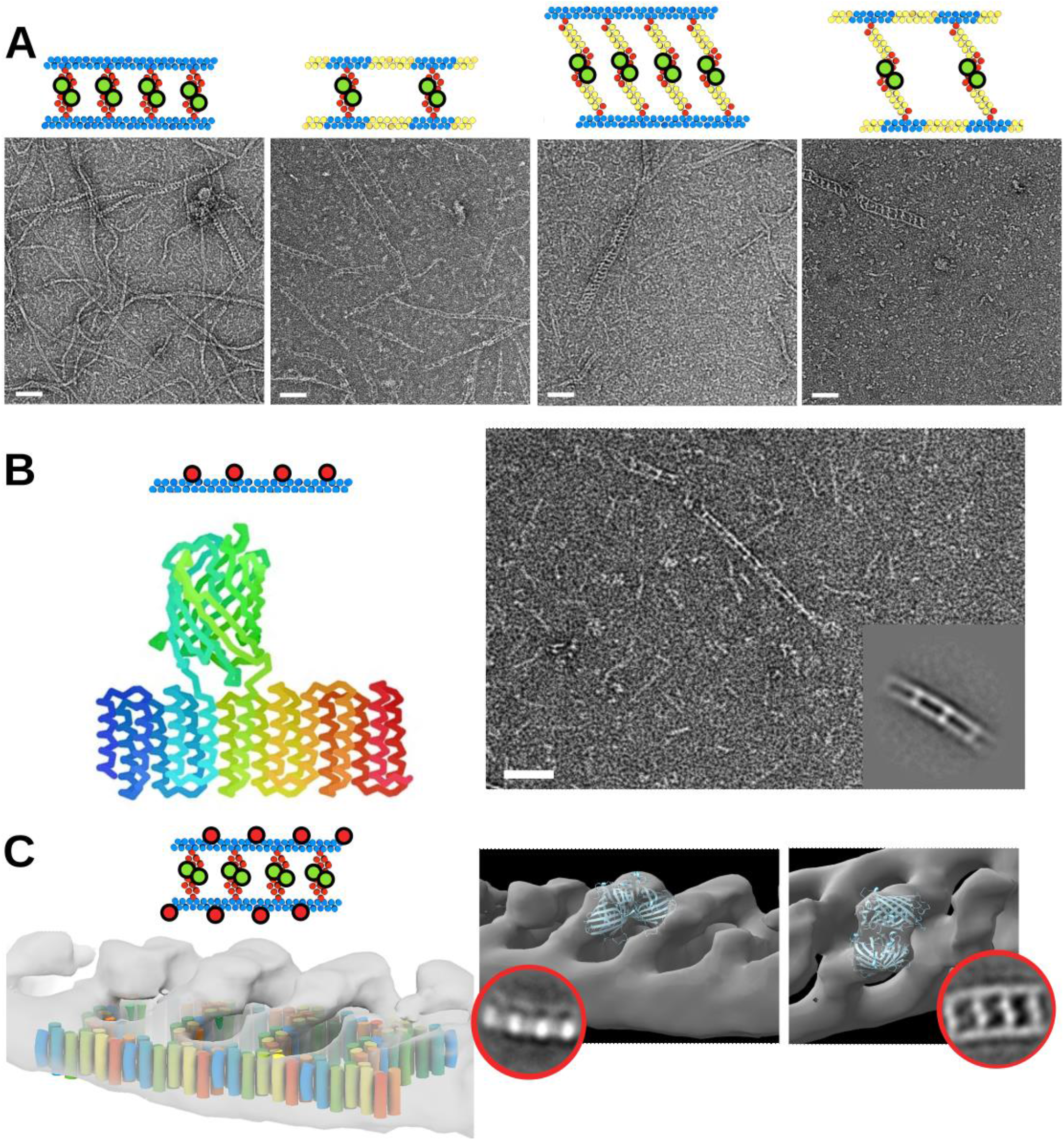
Additional nsEM data for train track designs. (A) Comparison of assembly for the 4 sizes of train track with GFP fused to the C2 module, all prepared under the same conditions of guanidine HCl denaturation of both components and then dialysis into TBS buffer when both components are mixed in approximate equimolar ratio before dialysis. 50 nm white scale bars. (B) A version of the rail/ Split module THR where the interface on the side for the rail is knocked out, and instead a mScarlet sequence (*24)* is inserted into one of the middle loops (this protein is still uncapped so it can assemble long-ways). An Alphafold2 prediction of one such design monomer is shown in rainbow. A nsEM micrograph (50 nm white scale bar) with class average inset shows that single fibrils rarely assemble end-to-end with more than 5 subunits, but perhaps weak dimerization of the mScarlet protein combined with the periodic templating of the rail leads to more stable double-fibrils. (C) An example of the base size train track where density of the GFP was visible in the nsEM reconstruction. An Alphfold2 predicted sfGFP (*23*) dimer was fit into the density above the obvious train track pattern and matched well.

**fig. S25.**
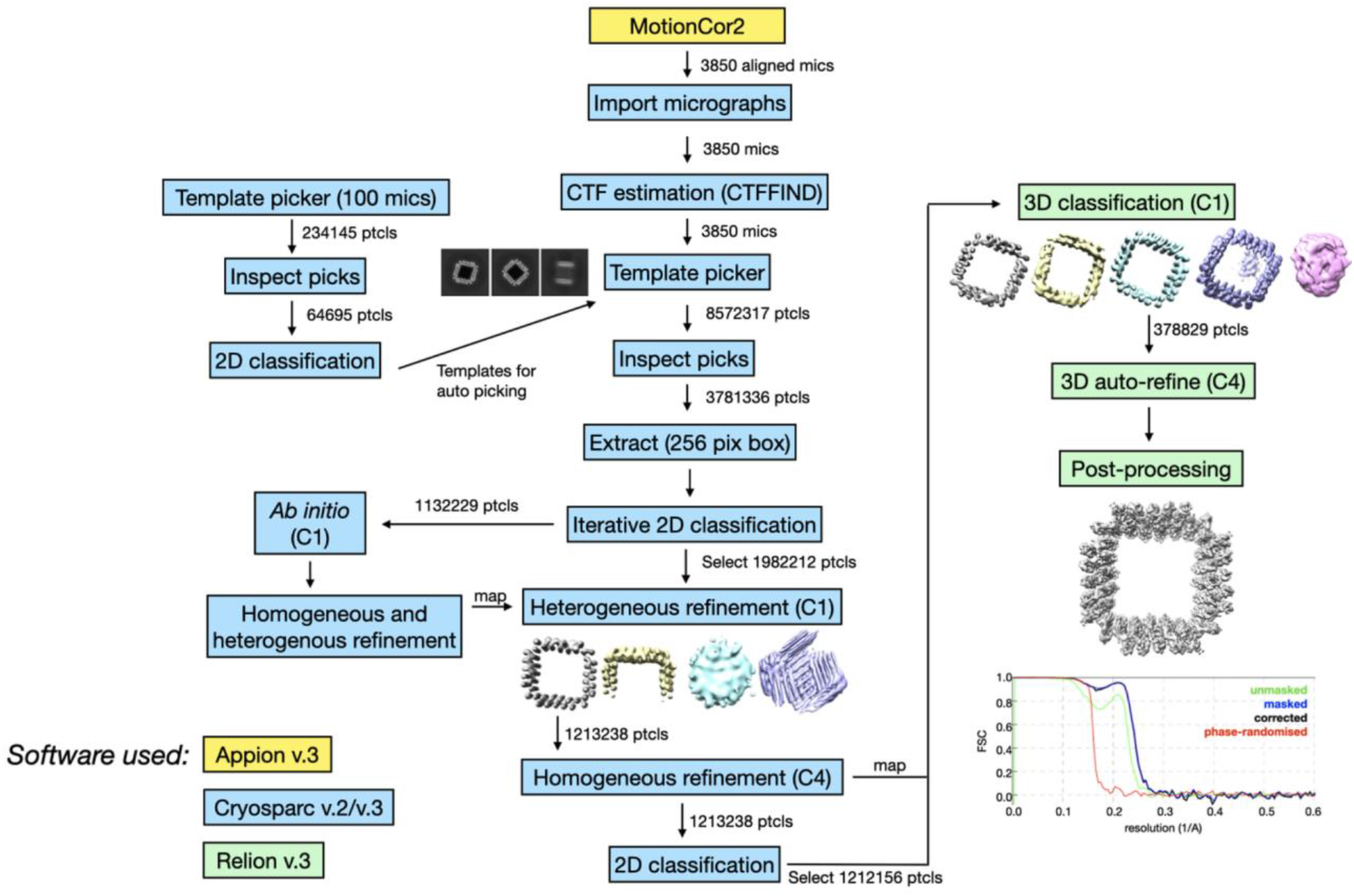
cryo-EM data processing pipeline used for *sC4*.

**fig. S26.**
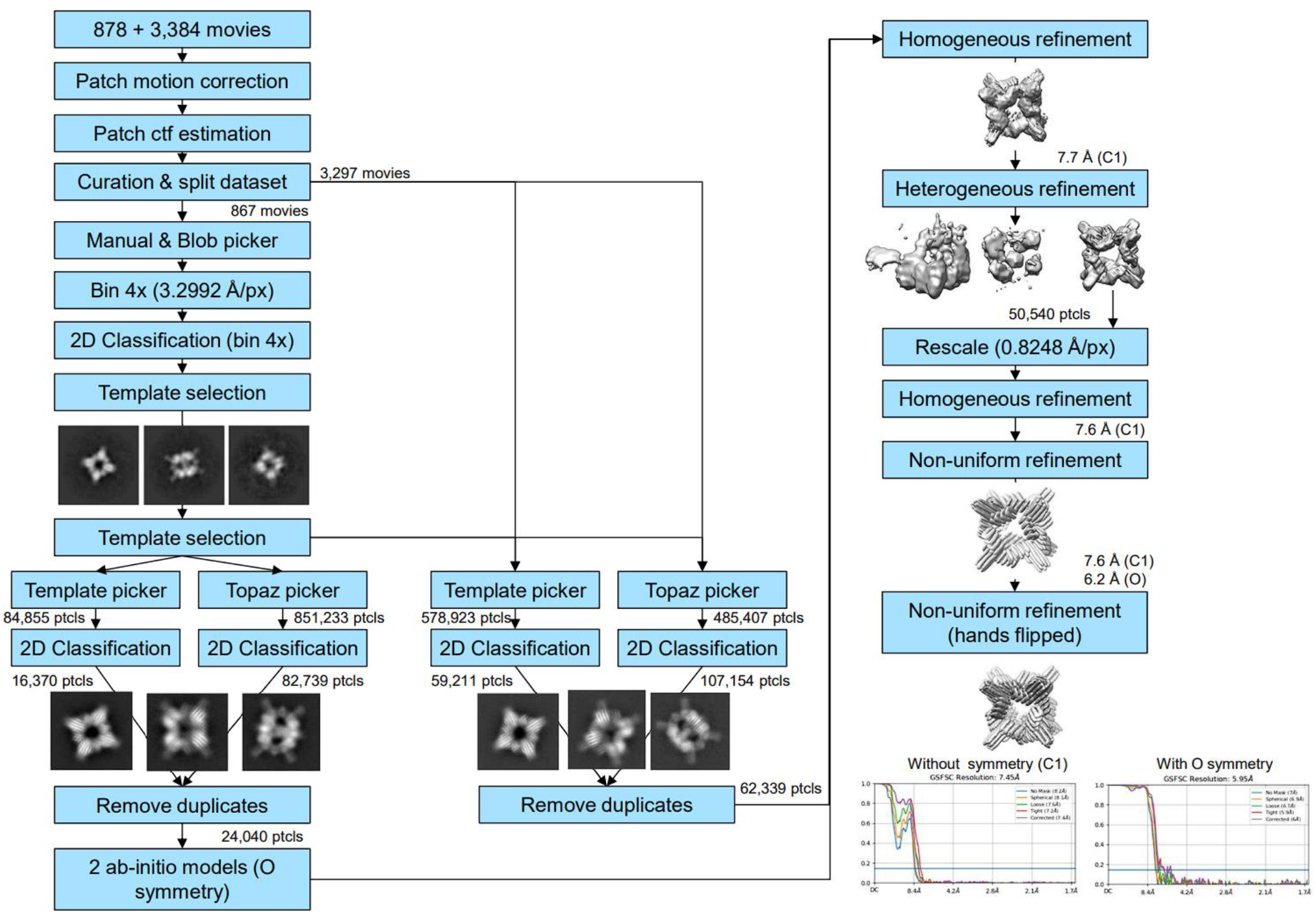
cryo-EM data processing pipeline used for *cage_03_10*.

**fig. S27.**
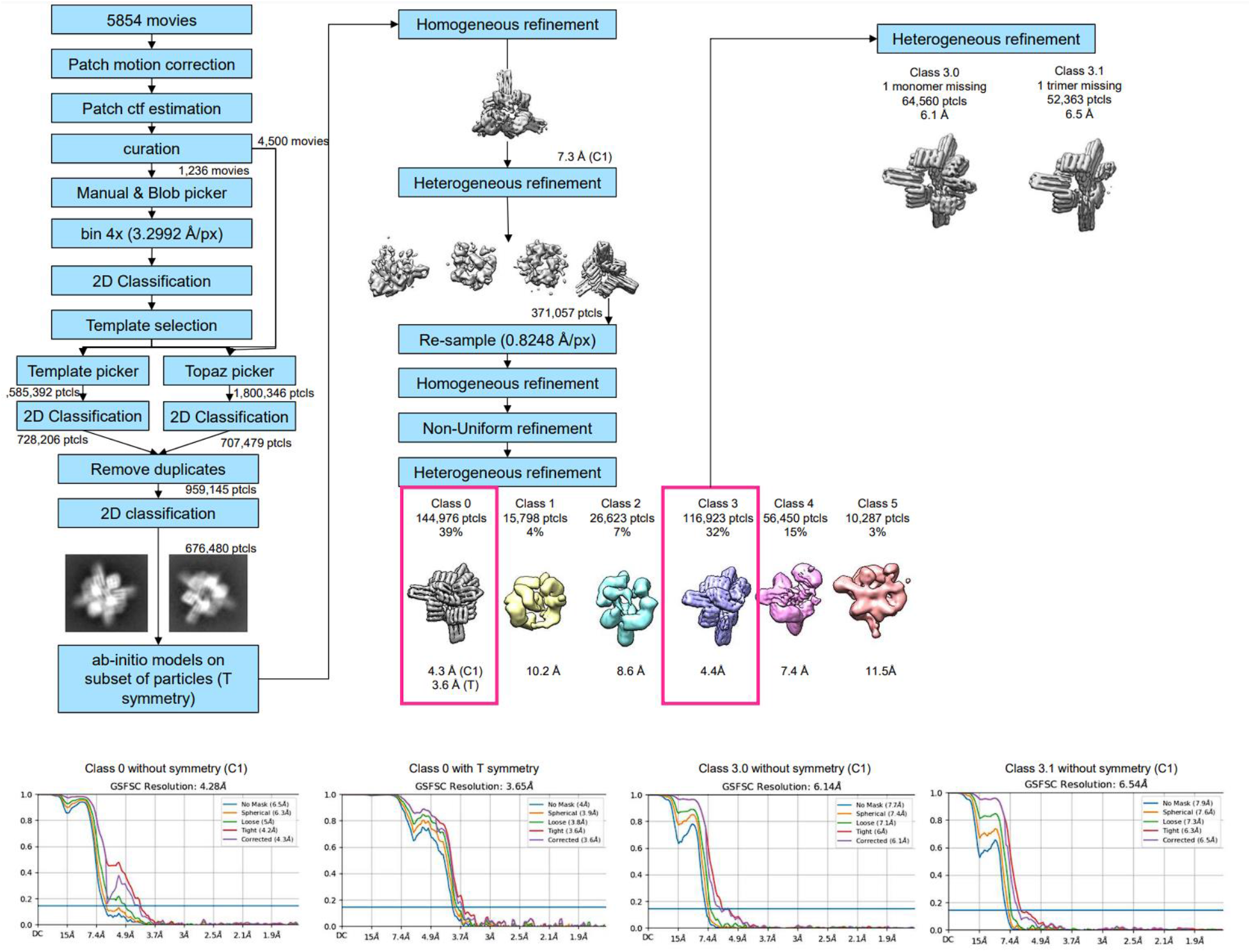
cryo-EM data processing pipeline used for *cage_T3_5*.

**fig. S28.**
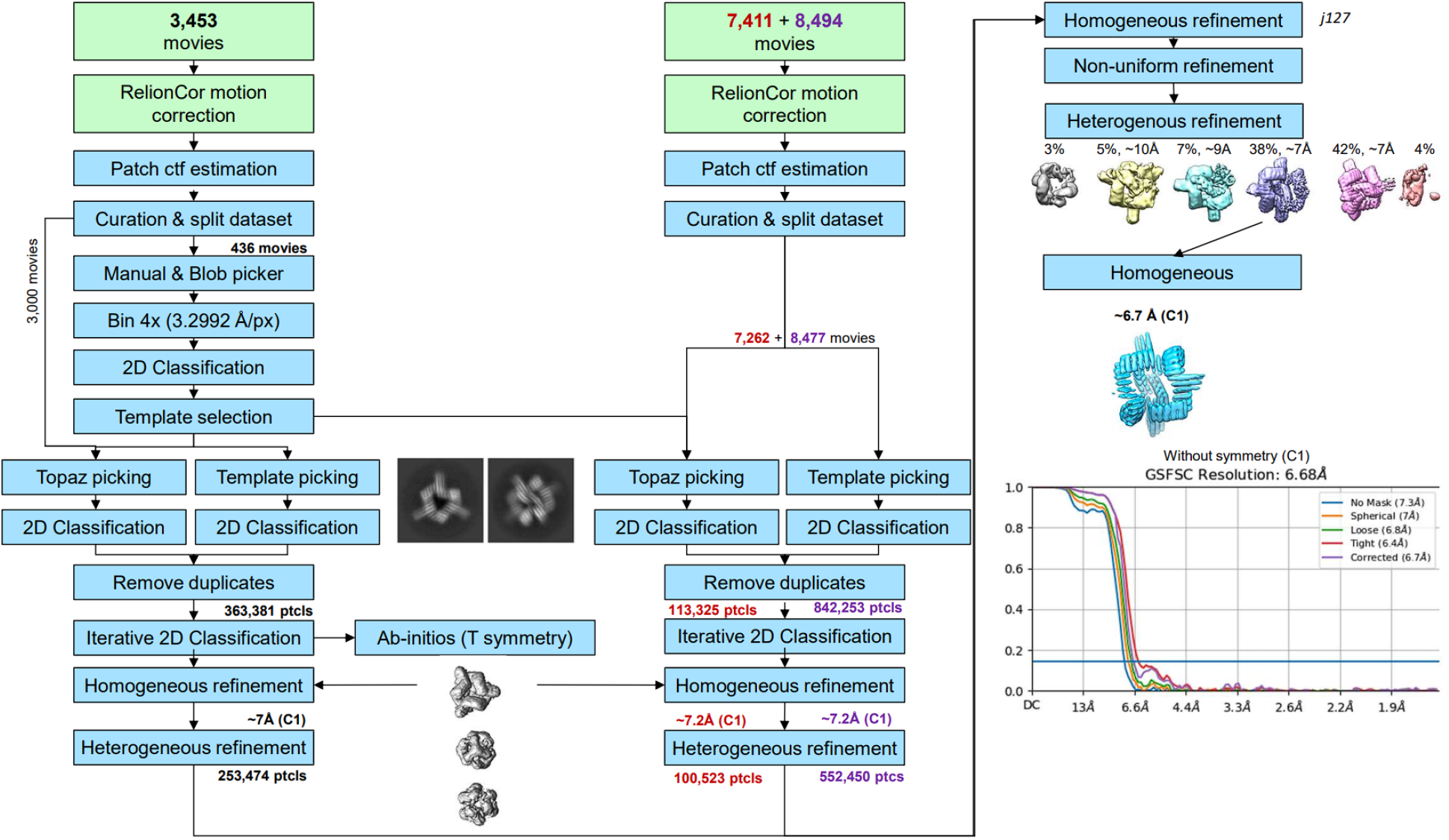
cryo-EM data processing pipeline used for *cage_T3_5+2*.

**fig. S29.**
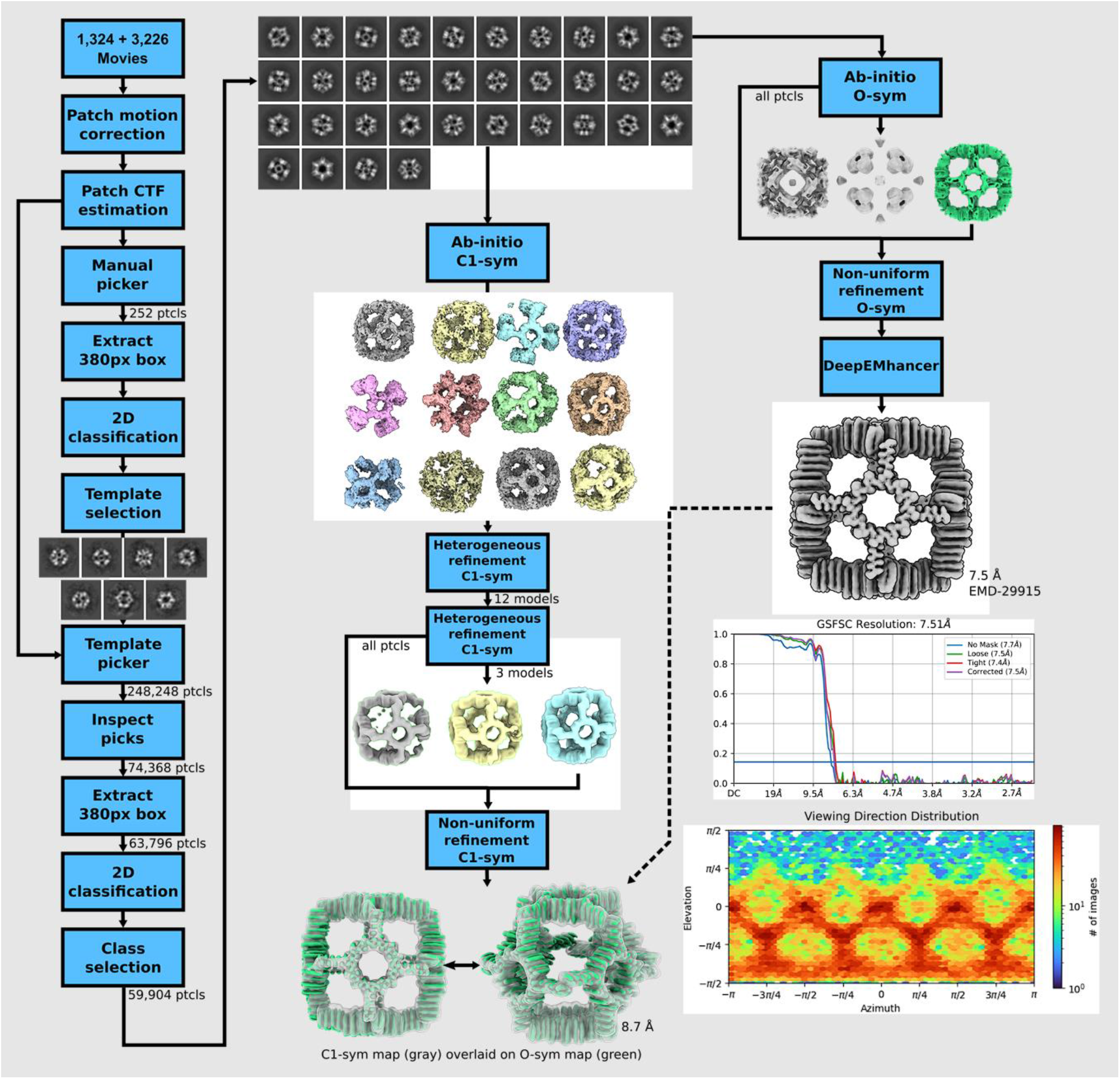
cryo-EM data processing pipeline used for *cage_O4_34*.

**fig. S30.**
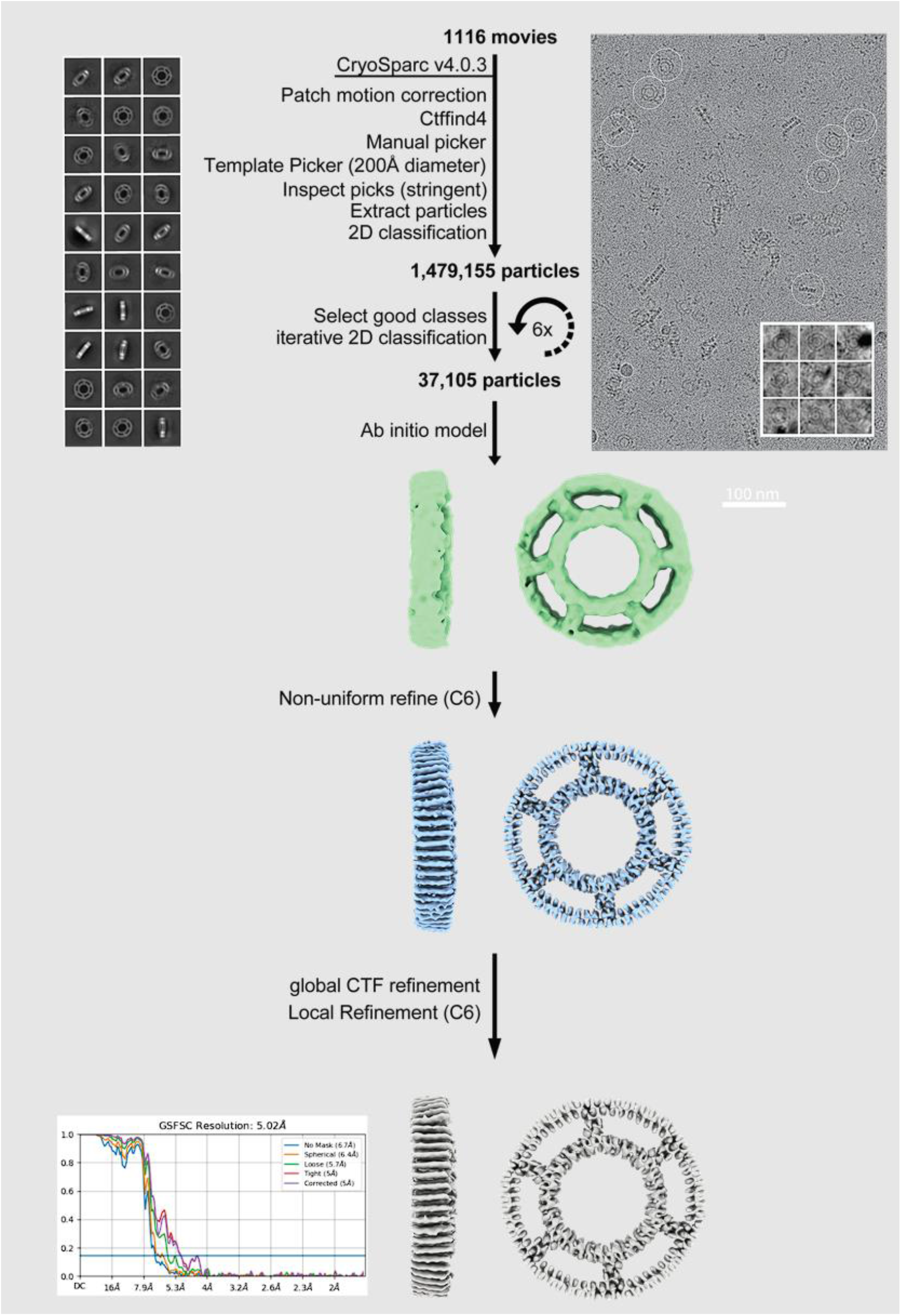
cryo-EM data processing pipeline used for *strut_C6_21*.

**fig. S31.**
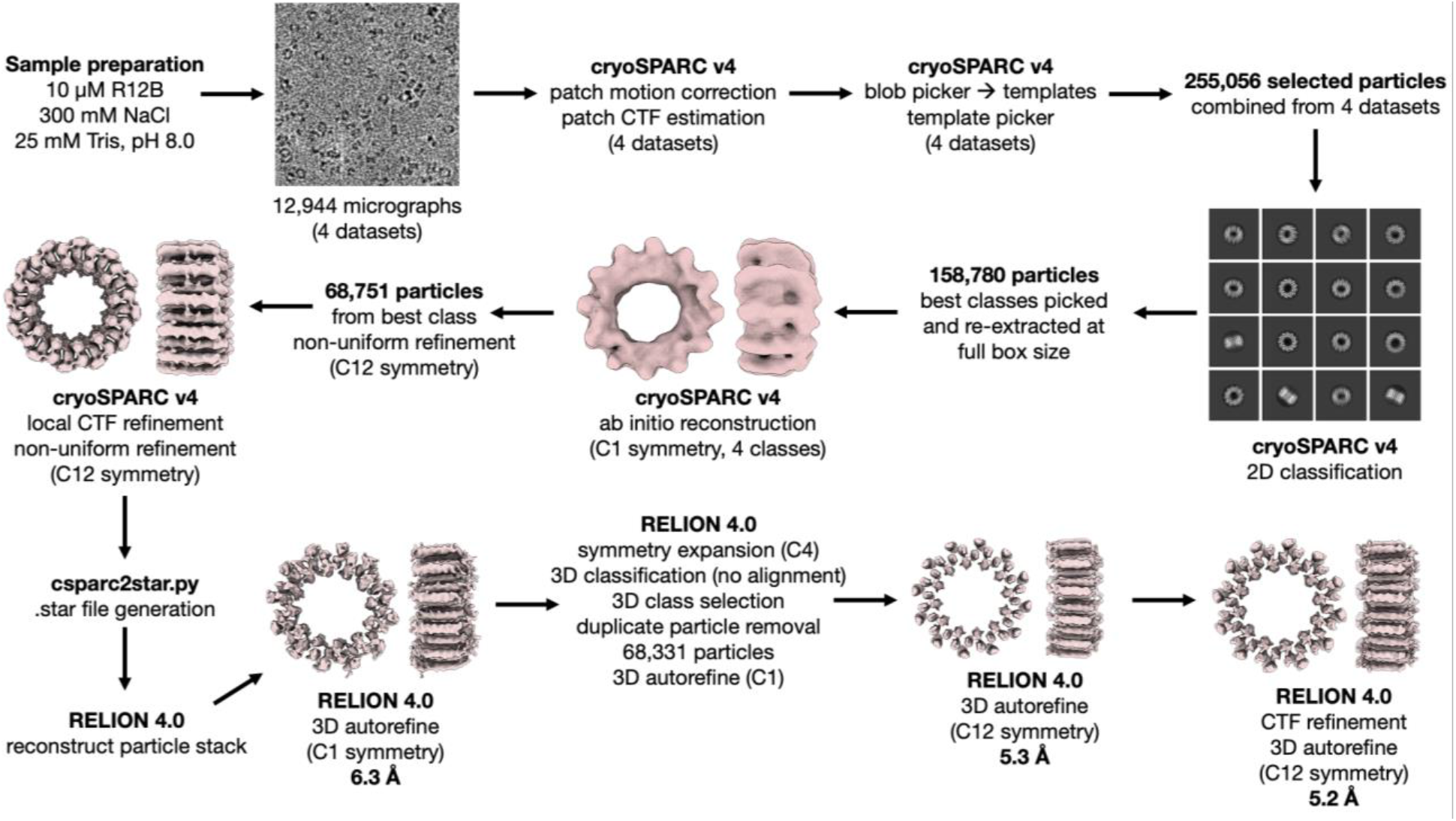
cryo-EM data processing pipeline used for *R12B*.

**Table S1.** Example cryo-EM data acquisition parameters for a subset of designs.

| sample | sC4 | sC4+2 | cage_03_10 | cage_T3_5 | cage_T3_5+2 | cage_T3_5+6 | tC3_A |
| --- | --- | --- | --- | --- | --- | --- | --- |
| Microscope | Krios | Arctica | Krios | Krios | Krios | Arctica | Krios |
| Voltage (kV) | 300 | 200 | 300 | 300 | 300 | 200 | 300 |
| Exposure navigation | Image shift, Leginon | Image shift, Leginon | Image shift, Leginon | Image shift, Leginon | Image shift, Leginon | Image shift, Leginon | Image shift, Leginon |
| Nominal magnification | 105,000x | 36,000x | 105,000x | 105,000x | 105,000x | 36,000x | 105,000x |
| Detector | Gatan K3 | Gatan K3 | Gatan K3 | Gatan K3 | Gatan K3 | Gatan K3 | Gatan K3 |
| Pixel size of movies in super-resolution (Å/pix) | 0.4124 | 0.548 | 0.4124 | 0.4124 | 0.4124 | 0.548 | 0.4124 |
| No. of frames | 40 | 40 | 50 | 50 | 50 | 40 | 40 |
| Dose rate (e-/Å<sup>2</sup>/s) | 29.4 | 20.3 | 29.4 | 29.4 | 29.4 | 20.3 | 29.2 |
| Exposure per frame (e-/Å<sup>2</sup>) | 1.18 | 1.42 | 1.18 | 1.18 | 1.18 | 1.42 | 1.17 |
| Total dose (e-/Å<sup>2</sup>) | 47.07 | 56.8 | 58.8 | 58.8 | 58.8 | 56.8 | 46.77 |
| Exposure time (s) | 1.6 | 2.8 | 2 | 2 | 2 | 2.8 | 1.6 |
| Defocus range (µm) | 0.8 - 2.0 | 1.1 - 2.9 | 0.6 - 2.2 | 0.6 - 2.2 | 0.6 - 2.2 | 0.6-2.4 | 0.9 - 2.5 |
| No. of micrographs acquired | 3,850 | 207 | 4,262 | 5,854 | 19,358 | 90 | 438 |

**Table S2.** Example cryo-EM image processing details for a subset of designs.

| Image Processing | sC4 | cage_O3_10 |  | cage_T3_5 |  |  |  | cage_T3_5+2 |
| --- | --- | --- | --- | --- | --- | --- | --- | --- |
| Software | Cryo SPARC, Relion | CryoSPARC |  | CryoSPARC |  |  |  | CryoSPARC |
| No. of extracted particles | 3,781,336 | 2,000,418 |  | 959,145 |  |  |  | 5,440,898 |
| No. of final particles | 378,829 | 49,466 |  | 144,976 | 144,976 | 64,560 | 52,363 | 166,363 |
| Box size (pixels) | 256 | 512 |  | 360 | 360 | 360 | 360 | 360 |
| Symmetry | C4 | C1 | O | C1 | T | C1<br>(1 chain missing) | C1<br>(1 trimer missing) | C1 |
| Map resolution (Å):<br>FSC 0.143<br>(unmasked / masked) | 4.06 / 3.91 | 8.2 / 7.4 | 7.0 / 6.0 | 6.5 / 4.3 | 4/ 3.6 | 7.7 / 6.1 | 7.9 / 6.5 | 7.3 / 6.7 |
| Map sharpening B factor (Å<sup>2</sup>) | -224.3 | none | none | none | -165 | none | none | none |
| Sphericity from 3DFSC (unmasked/ma<br>sked) | 0.851 / 0.952 | 0.660 / 0.737 | 0.958 / 0.971 | 0.965 / 0.967 | 0.840 / 0.989 | 0.658 / 0.912 | 0.635 / 0.751 | 0.911 / 0.912 |
| EMDB ID: | EMD-29974 | EMD-40070 | EMD-40071 | EMD-40075 | EMD-40074 | EMD-40073 | EMD-40072 | EMD-40076 |
| EMPIAR |  |  |  |  |  |  |  |  |

**Table S3.** Coordinate refinement details for cryo-EM C4 model of *sC4*.

|  |  |
| --- | --- |
| **Model refinement** | sC4 |
| Map CC (mask) | 0.791 |
| Map CC (volume) | 0.764 |
| Map CC (peaks) | 0.648 |
| R.m.s. deviations (bonds) | 0.007 |
| R.m.s. deviations (angles) | 0.861 |
| Ramachandran plot values (%) |  |
| outliers | 0.00 |
| allowed | 1.62 |
| favored | 98.38 |
| Rotamer outliers (%) | 0.00 |
| C-beta deviations (%) | 0.00 |
| CaBLAM outliers (%) | 0.61 |
| Overall score (Molprobit<sup>62</sup>) | 1.81 |
| Clashscore | 20.54 |
| PDB ID | 8GEL |

**Table S4.** Cryo-EM image processing details for *cage_04_34*.

|  |  |
| --- | --- |
|  | cage_O4_34 |
| Microscope | Tundra |
| Voltage (kV) | 100 |
| Detector | CETA-F |
| Recording mode | Fractionation |
| Magnification | 110,000 |
| Movie micrograph pixel size (Å) | 1.248 |
| Dose rate (e<sup>-</sup>/Å<sup>2</sup>/s) | 27.7 |
| No. of frames per movie micrograph | 21 |
| Frame exposure time (ms) | 71.4 |
| Movie micrograph exposure time (s) | 1.5 |
| Total dose (e<sup>-</sup>/Å<sup>2</sup>) | 4.15 |
| Under focus range (μm) | -0.5 to -2.2 |
| Number of movie micrographs | 4,550 |
| Total number of picked particles | 248,248 |
| Particles in the final reconstruction | 59,904 |
| Map symmetry | O |
| Map resolution (GS-FSC) | 7.5 Å |
| EMDB ID | EMD-29915 |

**Table S5.** Crystallographic data collection and refinement.

| Data Collection | THR1 (8G9J) | THR2 (8G9K) | THR5 (8GA7) | THR6 (8GA6) |
| --- | --- | --- | --- | --- |
| Space group | $P 4_1 2_1 2$ | $P 2_1 2_1 2_1$ | $P 2_1 2_1 2_1$ | $P 2_1$ |
| Cell dimensions |  |  |  |  |
| a, b, c (Å) | 49.70, 49.70, 129.26 | 46.39, 61.19, 149.02 | 46.41, 65.06, 135.28 | 34.96, 120.42, 52.27 |
| $\alpha, \beta, \gamma$ (°) | 90, 90, 90 | 90, 90, 90 | 90, 90, 90 | 90, 105.65, 90 |
| Resolution (Å) | 46.39 - 2.5 (2.75 - 2.5) | 56.61 - 2.48 (2.56 - 2.48) | 38.27 - 2.93 (3.08 - 2.93) | 46.44 - 2.5 (2.58 - 2.5) |
| $R_{\text{merge}}$ | 0.0699 (0.431) | 0.063 (0.123) | 0.204 (1.178) | 0.137 (0.778) |
| $R_{\text{pim}}$ | 0.0698 (0.4314) | 0.025 (0.491) | 0.079 (0.439) | 0.084 (0.463) |
| $I/\sigma(I)$ | 6.62 (1.20) | 17.19 (6.68) | 7.79 (2.61) | 5.24 (1.69) |
| $CC_{1/2}$ | 0.999 (0.870) | 0.999 (0.999) | 0.997 (0.939) | 0.982 (0.722) |
| Completeness (%) | 98.64 (95.58) | 98.91 (98.79) | 98.34 (98.93) | 97.98 (95.07) |
| Redundancy | 1.7 (1.3) | 7.4 (7.6) | 7.5 (7.9) | 3.7 (3.8) |
| Refinement |  |  |  |  |
| Resolution (Å) | 46.39 - 2.5 (2.75 - 2.5) | 56.61 - 2.48 (2.56 - 2.48) | 38.27 - 2.93 (3.08 - 2.93) | 46.44 - 2.5 (2.58 - 2.5) |
| No. reflections | 6005 (1404) | 15526 (1388) | 9149 (1290) | 14161 (1330) |
| $R_{\text{work}} / R_{\text{free}}$ | 0.2333 (0.2987)/ 0.2842 (0.3763) | 0.2642 (0.3305) 0.3010 (0.3548) | 0.2756 (0.3005) 0.3144 (0.3387) | 0.2878 (0.3062) 0.3264 (0.3587) |
| No. atoms |  |  |  |  |
| Protein | 1486 | 2965 | 3438 | 3454 |
| Water | 13 | 39 | 0 | 167 |
| Ramachandran Favored/allowed Outlier (%) | 98.49/ 1.51/ 0.00 | 97.99/ 0.76/ 0.25 | 97.60/ 2.00/ 0.40 | 97.98/ 1.26/ 0.84 |
| $R.m.s. \text{ deviations}$ | | | | |
| Bond lengths (Å) | 0.003 | 0.001 | 0.002 | 0.001 |
| Bond angles (°) | 0.520 | 0.400 | 0.400 | 0.84 |
| $B_{\text{factors}}$ (Å<sup>2</sup>) | | | | |
| Protein | 46.72 | 78.57 | 58.84 | 41.70 |
| Water | 40.71 | 73.05 | n/a | 42.72 |

## References

1. Berman, H. M. et al. The Protein Data Bank. Nucleic Acids Res. 28, 235–242 (2000).

2. Thomson, A. R. et al. Computational design of water-soluble α-helical barrels. Science 346, 485– 488 (2014).

3. Wicky, B. I. M. et al. Hallucinating symmetric protein assemblies. Science 378, 56–61 (2022).

4. Fallas, J. A. et al. Computational design of self-assembling cyclic protein homo-oligomers. Nat. Chem. 9, 353–360 (2017).

5. Ljubetič, A. et al. Design of coiled-coil protein-origami cages that self-assemble in vitro and in vivo. Nat. Biotechnol. 35, 1094–1101 (2017).

6. Hsia, Y. et al. Design of multi-scale protein complexes by hierarchical building block fusion. Nat. Commun. 12, 1–10 (2021).

7. King, N. P. et al. Computational design of self-assembling protein nanomaterials with atomic level accuracy. Science 336, 1171–1174 (2012).

8. Sheffler, W. et al. Fast and versatile sequence-independent protein docking for nanomaterials design using RPXDock. PLOS Computational Biology 19(5): e1010680 (2023) doi:10.1371/journal.pcbi.1010680.

9. Bethel, N. P., et al. Precisely patterned nanofibers made from extendable protein multiplexes. bioRxiv 2022.10.14.511843 (2022) doi:10.1101/2022.10.14.511843.

10. Brodin, J. D. et al. Metal-directed, chemically tunable assembly of one-, two- and three-dimensional crystalline protein arrays. Nat. Chem. 4, 375–382 (2012).

11. Sinclair, J. C., Davies, K. M., Vénien-Bryan, C. & Noble, M. E. M. Generation of protein lattices by fusing proteins with matching rotational symmetry. Nat. Nanotechnol. 6, 558–562 (2011).

12. Ben-Sasson, A. J. et al. Author Correction: Design of biologically active binary protein 2D materials. Nature 591, E16–E16 (2021).

13. Li, Z., et al. Accurate Computational Design of 3D Protein Crystals. bioRxiv 2022.11.18.517014 (2022) doi:10.1101/2022.11.18.517014.

14. Padilla, J. E., Colovos, C. & Yeates, T. O. Nanohedra: using symmetry to design self assembling protein cages, layers, crystals, and filaments. Proc. Natl. Acad. Sci. U. S. A. 98, 2217–2221 (2001).

15. Woolfson, D. N. Understanding a protein fold: The physics, chemistry, and biology of α-helical coiled coils. J. Biol. Chem. 299, (2023).

16. Grigoryan, G. & Degrado, W. F. Probing designability via a generalized model of helical bundle geometry. J. Mol. Biol. 405, 1079–1100 (2011).

17. Brunette, T. J. et al. Exploring the repeat protein universe through computational protein design. Nature 528, 580–584 (2015).

18. Huang, P.-S. et al. High thermodynamic stability of parametrically designed helical bundles. Science 346, 481–485 (2014).

19. Alford, R. F. et al. The Rosetta All-Atom Energy Function for Macromolecular Modeling and Design. J. Chem. Theory Comput. 13, 3031–3048 (2017).

20. Dauparas, J. et al. Robust deep learning–based protein sequence design using ProteinMPNN. Science 378, 49–56 (2022).

21. Correnti, C. E. et al. Engineering and functionalization of large circular tandem repeat protein nanoparticles. Nat. Struct. Mol. Biol. 27, 342–350 (2020).

22. Coxeter, H. S. M. Regular Polytopes. (Courier Corporation, 1973).

23. Yeates, T. O. Geometric Principles for Designing Highly Symmetric Self-Assembling Protein Nanomaterials. Annu. Rev. Biophys. 46, 23–42 (2017).

24. Walshaw, J. & Woolfson, D. N. Extended knobs-into-holes packing in classical and complex coiled-coil assemblies. J. Struct. Biol. 144, 349–361 (2003).

25. Pédelacq, J.-D., Cabantous, S., Tran, T., Terwilliger, T. C. & Waldo, G. S. Engineering and characterization of a superfolder green fluorescent protein. Nat. Biotechnol. 24, 79–88 (2005).

26. Bindels, D. S. et al. mScarlet: a bright monomeric red fluorescent protein for cellular imaging. Nat. Methods 14, 53–56 (2016).

27. Kendrew, J. C. et al. A Three-Dimensional Model of the Myoglobin Molecule Obtained by X-Ray Analysis. Nature 181, 662–666 (1958).

28. Pyles, H., Zhang, S., De Yoreo, J. J. & Baker, D. Controlling protein assembly on inorganic crystals through designed protein interfaces. Nature 571, 251–256 (2019).

29. Davila-Hernandez, F. A., et al. Directing polymorph specific calcium carbonate formation with de novo protein templates. bioRxiv (2023).

30. Kibler, R. D., et al. Stepwise design of pseudosymmetric protein hetero-oligomers. bioRxiv 2023.04.07.535760 (2023) doi:10.1101/2023.04.07.535760.

31. Wintersinger, C. M. et al. Multi-micron crisscross structures grown from DNA-origami slats. Nat. Nanotechnol. 18, 281–289 (2023).

32. Bohlin, J., Turberfield, A. J., Louis, A. A. & Šulc, P. Designing the Self-Assembly of Arbitrary Shapes Using Minimal Complexity Building Blocks. ACS Nano 17, 5387–5398 (2023).

33. Sigl, C. et al. Programmable icosahedral shell system for virus trapping. Nat. Mater. 20, 1281–1289 (2021).

34. Wagenbauer, K. F., Sigl, C. & Dietz, H. Gigadalton-scale shape-programmable DNA assemblies. Nature 552, 78–83 (2017).

35. Chaudhury, S., Lyskov, S. & Gray, J. J. PyRosetta: a script-based interface for implementing molecular modeling algorithms using Rosetta. Bioinformatics 26, 689–691 (2010).

36. Jumper, J. et al. Highly accurate protein structure prediction with AlphaFold. Nature 596, 583–589 (2021).

37. Boyken, S. E. et al. De novo design of protein homo-oligomers with modular hydrogen-bond network-mediated specificity. Science 352, 680–687 (2016).

38. Studier, F. W. Protein production by auto-induction in high density shaking cultures. Protein Expr. Purif. 41, 207–234 (2005).

39. Schneidman-Duhovny, D., Hammel, M. & Sali, A. FoXS: a web server for rapid computation and fitting of SAXS profiles. Nucleic Acids Res. 38, W540–4 (2010).

40. Nannenga, B. L., Iadanza, M. G., Vollmar, B. S. & Gonen, T. Overview of electron crystallography of membrane proteins: crystallization and screening strategies using negative stain electron microscopy. Curr. Protoc. Protein Sci. Chapter 17, Unit17.15 (2013).

41. Punjani, A., Rubinstein, J. L., Fleet, D. J. & Brubaker, M. A. cryoSPARC: algorithms for rapid unsupervised cryo-EM structure determination. Nat. Methods 14, 290–296 (2017).

42. Suloway, C. et al. Automated molecular microscopy: the new Leginon system. J. Struct. Biol. 151, 41–60 (2005).

43. Zheng, S. Q. et al. MotionCor2: anisotropic correction of beam-induced motion for improved cryo-electron microscopy. Nat. Methods 14, 331–332 (2017).

44. Lander, G. C. et al. Appion: an integrated, database-driven pipeline to facilitate EM image processing. J. Struct. Biol. 166, 95–102 (2009).

45. Rohou, A. & Grigorieff, N. CTFFIND4: Fast and accurate defocus estimation from electron micrographs. J. Struct. Biol. 192, 216–221 (2015).

46. Zivanov, J. et al. New tools for automated high-resolution cryo-EM structure determination in RELION-3. Elife 7, e42166 (2018).

47. Pettersen, E. F. et al. UCSF Chimera--a visualization system for exploratory research and analysis. J. Comput. Chem. 25, 1605–1612 (2004).

48. Liebschner, D. et al. Macromolecular structure determination using X-rays, neutrons and electrons: recent developments in Phenix. Acta Crystallogr D Struct Biol 75, 861–877 (2019).

49. Emsley, P., Lohkamp, B., Scott, W. G. & Cowtan, K. Features and development of Coot. Acta Crystallogr. D Biol. Crystallogr. 66, 486–501 (2010).

50. Tan, Y. Z. et al. Addressing preferred specimen orientation in single-particle cryo-EM through tilting. Nat. Methods 14, 793–796 (2017).

51. Sanchez-Garcia, R. et al. DeepEMhancer: a deep learning solution for cryo-EM volume post-processing. Commun Biol 4, 874 (2021).

52. Pettersen, E. F. et al. UCSF ChimeraX: Structure visualization for researchers, educators, and developers. Protein Sci. 30, 70–82 (2021).

53. Kidmose, R. T. et al. Namdinator - automatic molecular dynamics flexible fitting of structural models into cryo-EM and crystallography experimental maps. IUCrJ 6, 526–531 (2019).

54. Mastronarde, D. N. Automated electron microscope tomography using robust prediction of specimen movements. J. Struct. Biol. 152, 36–51 (2005).

55. Asarnow, D., Palovcak, E. & Cheng, Y. asarnow/pyem: UCSF pyem v0.5. (2019). doi:10.5281/zenodo.3576630.

56. Kimanius, D., Dong, L., Sharov, G., Nakane, T. & Scheres, S. H. W. New tools for automated cryo-EM single-particle analysis in RELION-4.0. Biochem. J 478, 4169–4185 (2021).

57. Kabsch, W. XDS. Acta Crystallogr. D Biol. Crystallogr. 66, 125–132 (2010).

58. Winn, M. D. et al. Overview of the CCP4 suite and current developments. Acta Crystallogr. D Biol. Crystallogr. 67, 235–242 (2011).

59. McCoy, A. J. et al. Phaser crystallographic software. J. Appl. Crystallogr. 40, 658–674 (2007).

60. Adams, P. D. et al. PHENIX: a comprehensive Python-based system for macromolecular structure solution. Acta Crystallogr. D Biol. Crystallogr. 66, 213–221 (2010).

61. Emsley, P. & Cowtan, K. Coot: model-building tools for molecular graphics. Acta Crystallogr. D Biol. Crystallogr. 60, 2126–2132 (2004).

62. Williams, C. J. et al. MolProbity: More and better reference data for improved all-atom structure validation. Protein Sci. 27, 293–315 (2018).

63. Wang, Z. et al. Structure of the Marine Siphovirus TW1: Evolution of Capsid-Stabilizing Proteins and Tail Spikes. Structure 26, 238–248.e3 (2018).

